# Somatic haplotype reconstruction and variant recalibration from tumor-only long-read sequencing

**DOI:** 10.64898/2026.09.14.751225

**Authors:** Zhen-Yu Chen, Zhenxian Zheng, Ruibang Luo, Hsu-Fen Fu, Yung-Jen Yang, Yao-Ting Huang

## Abstract

Separating somatic from germline variants and reconstructing somatic haplotypes are the two central problems of tumor-only cancer genome analysis. Long reads carry the linkage needed to solve both, but chromosome-scale loss of heterozygosity (LOH) and an unknown degree of normal-cell admixture blur the distinction between somatic and germline haplotypes. Here we present LongPhase-TO, the first method to reconstruct somatic haplotypes from a tumor sample alone. LongPhase-TO co-phases germline and somatic alleles in a unified graph instead of mapping somatic variants onto germline haplotypes, and resolves LOH and tumor DNA fraction internally from heterozygosity depletion and haplotype imbalance without a copy-number and ploidy model. Across eight datasets from six cancer cell lines, LongPhase-TO increased haplotype block N50 by a median of 2.9-fold relative to germline phasers. It also consistently improved somatic single-nucleotide variant (SNV) and indel calls from ClairS-TO and DeepSomatic-TO, raising mean F1 from 0.55 to 0.62 and 0.65 for SNVs and from 0.19 to 0.23 for indels, with the largest gains at low tumor DNA fraction. Across breast, melanoma and lung cancer cell lines, LongPhase-TO improves the accuracy of existing somatic callers and reconstructs megabase-scale somatic haplotypes.

## Introduction

Cancer genomes acquire somatic mutations that distinguish tumor cells from the inherited germline and shape tumor development, diagnosis, and therapeutic response [1–3]. Cataloging these variants alone does not reveal their allelic relationships: mutations on the same chromosome copy (in cis) can have different functional and evolutionary consequences from those on opposite copies (in trans) [4, 5]. Long reads span multiple heterozygous loci, providing direct linkage information for long-range haplotype reconstruction [6, 7]. Germline phasers such as WhatsHap, HapCUT2, and LongPhase use this linkage to partition reads between the two inherited parental haplotypes [8–10]. Their diploid formulation, however, does not capture the somatic haplotype structure as mutations accumulate during clonal evolution [11].

Phasing also improves long-read variant calling by organizing reads according to local haplotypes, thereby strengthening coherent allelic signals relative to sequencing noise. Germline callers such as DeepVariant and Clair3 incorporate haplotype-aware read representations into deep neural networks to improve genotyping accuracy [12– 14]. The same principle applies to somatic small- and structural-variant calling, where genuine somatic alleles generally remain associated with one parental haplotype, whereas sequencing artifacts are less likely to show consistent haplotype support [15– 17]. In matched tumor-normal workflows, the normal sample provides complementary patient-specific germline evidence, allowing somatic variants to be distinguished from inherited alleles [18–20]. Without a matched normal, tumor-only calling must separate low-frequency somatic mutations from sequencing errors and rare germline variants [21–23]. Existing tumor-only callers rely on panels of normals (PoNs) to remove known polymorphisms and recurrent artifacts, but these resources miss private germline variation and sample-specific artifacts [18]. Tumor-only callers such as DeepSomatic-TO and ClairS-TO therefore combine learned sequence features, external filtering resources, and local phasing information to strengthen somatic variant classification [15, 24].

Local phase-aware calling, however, is not equivalent to reconstructing extended tumor-specific haplotypes. Megabase-scale loss of heterozygosity (LOH) depletes the heterozygous markers that link flanking phase blocks [25–27], while normal-cell admixture dilutes somatic alleles and weakens the evidence for their haplotype of origin [28–30]. Together these features limit the reconstruction of contiguous somatic haplotypes [31]. Recent long-read methods instead resolve the two parental haplotypes and assign somatic variants to that germline backbone. For instance, SAVANA assigns each somatic structural variant and copy-number aberration to a haplotype using reads pre-tagged by WhatsHap or LongPhase [32]. Wakhan extends pre-phased germline blocks to chromosome scale using haplotype-specific coverage [33]. TumorLens phases the tumor against germline haplotypes derived from a matched normal [34]. However, these methods indirectly map somatic variants onto germline haplotypes rather than phasing them and resolving LOH simultaneously.

Normal-cell admixture presents a second barrier, because the tumor DNA fraction dictates the allelic support expected at each somatic site. The same methods either infer this fraction from copy number or take it as input. SAVANA reads it from B-allele frequencies across LOH segments [32] and Wakhan fits it jointly with ploidy to haplotype-specific coverage [33], as ASCAT and PURPLE do from allele-specific profiles [35, 36], whereas TumorLens recovers the correct copy number only when the tumor content is provided [34]. In most methods the fraction is supplied or fitted through a copy-number and ploidy model, and never feeds back into somatic phasing to account for dilution by admixed normal DNA.

Here we present LongPhase-TO, a tumor-only method that phases germline and somatic alleles jointly and separates somatic haplotypes from germline ones. LOH and tumor DNA fraction are resolved within the same graph from heterozygosity depletion and haplotype imbalance, without a copy-number and ploidy model. Across eight datasets from six cancer cell lines, LongPhase-TO improved somatic calls from ClairS-TO and DeepSomatic-TO and produced longer haplotypes than germline phasers.

## Results

### Overview of the LongPhase-TO framework

We developed LongPhase-TO, a somatic phasing method for tumor-only long-read sequencing that reconstructs somatic haplotypes and infers tumor DNA fraction in an LOH-aware manner (Figure 1; see Methods). The workflow proceeds in four stages: somatic variant recalibration, arm-level LOH detection, LOH-aware somatic phasing, and phase-aware DNA fraction estimation. First, candidate SNVs and indels from ClairS-TO or DeepSomatic-TO are recalibrated with a triplet-graph model built on a simple biological expectation (Figure 1(a)): because a somatic mutation arises post-zygotically on one parental chromosome, reads carrying its alternate allele should track a single germline haplotype, whereas residual germline variants and sequencing artifacts scatter across both. A trained model scores this local haplotype evidence to decide whether each candidate is retained. When the alignments carry 5-methylcytosine (5mC) tags, an optional classifier further filters the retained candidates using allele-specific CpG methylation (see Methods).

**Fig. 1.**
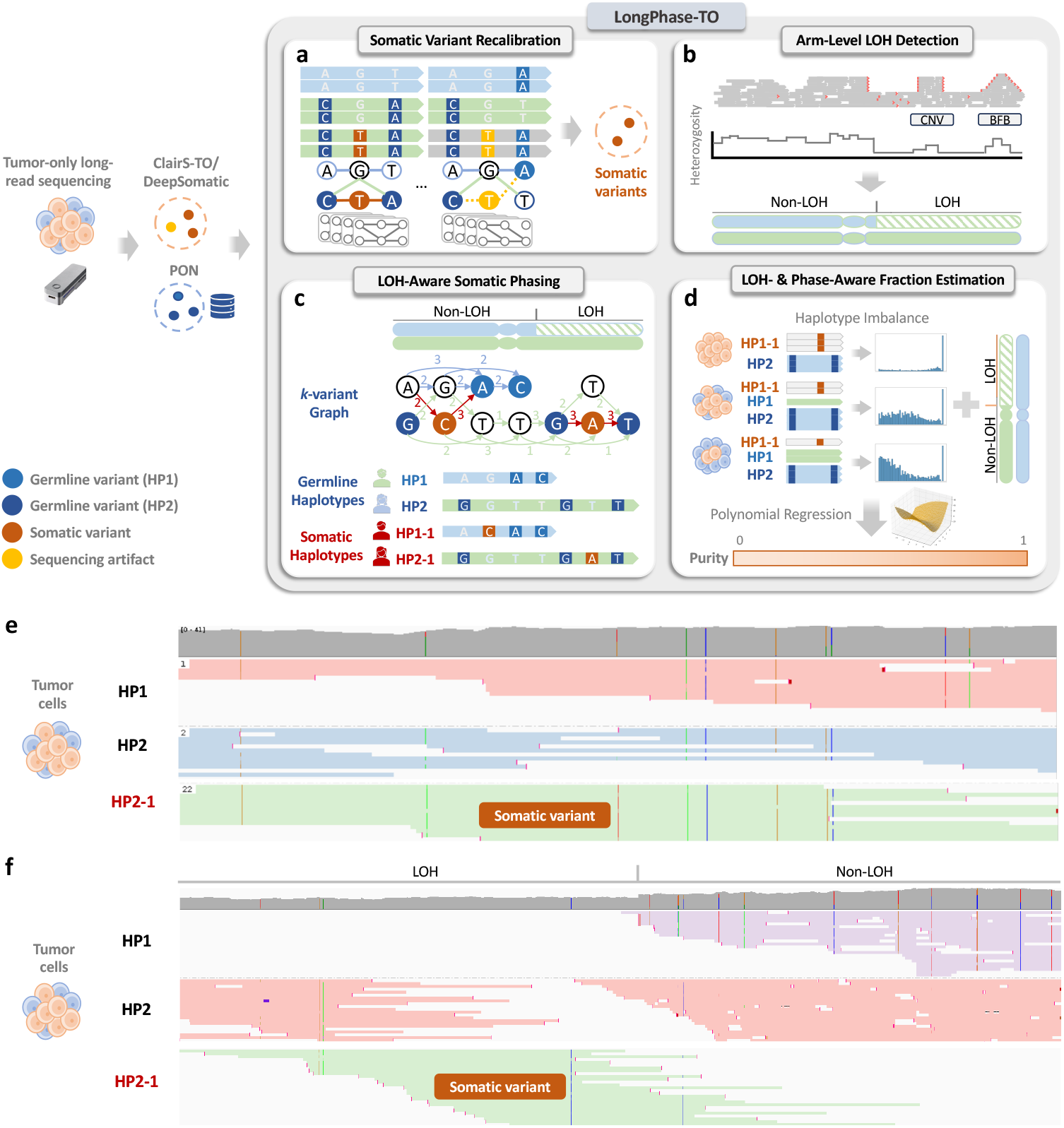
Overview of the LongPhase-TO somatic haplotype phasing framework. (a) Haplotype-aware recalibration refines initial somatic variant calls from ClairS-TO or DeepSomatic-TO by integrating PoNs and haplotype linkage information, distinguishing true somatic variants (orange) from sequencing artifacts (yellow) and germline variants (blue). (b) Chromosome-scale LOH detection identifies large-scale allelic imbalances by analyzing CNV/BFB intervals and heterozygosity ratios to distinguish LOH from non-LOH regions. (c) Somatic phasing assigns somatic variants to their originating germline haplotypes (HP1, HP2) and reconstructs derived somatic haplotypes (HP1-1, HP2-1), spanning LOH regions. (d) Tumor DNA fraction estimation quantifies tumor content based on haplotype imbalance, where deviation from balanced allelic representation reflects increasing tumor DNA fraction. (e) Haplotagged reads at a somatic SNV in HCC1395 HKU: reads are partitioned into the germline haplotypes HP1 and HP2 and the somatic haplotype HP2-1, which carries the somatic allele and descends from HP2. (f) Haplotagging across an LOH boundary in the same dataset: within the LOH segment (left), where HP1 is lost, somatic reads remain tagged HP2-1 rather than collapsing into HP2, and both parental haplotypes are recovered in the adjacent non-LOH region (right).

Second, LongPhase-TO delineates arm-level LOH with a breakpoint-guided procedure that jointly maps LOH and CNV/BFB intervals (Figure 1(b)). Small CNV/BFB islands often sit within a broad LOH tract and fragment it into many disjoint segments. LongPhase-TO reconciles the candidate intervals against the genome-wide heterozygosity profile to recover LOH at chromosome scale, where heterozygosity-depleted segments are retained as LOH and embedded high-heterozygosity CNV/BFB islands are absorbed into the surrounding LOH event.

Third, using the recalibrated variants and the LOH map, LongPhase-TO performs LOH-aware somatic phasing over a variant graph of germline and somatic alleles (Figure 1(c)). It reconstructs the two germline haplotypes (HP1, HP2) and, on reaching a somatic allele, branches a somatic haplotype (HP1-1 or HP2-1) from its parent. This lineage-aware traversal preserves clonal ancestry (e.g., HP1-1 from HP1) and extends both germline and somatic haplotypes across arm-level LOH, where conventional diploid phasers break.

Fourth, LongPhase-TO estimates tumor DNA fraction from the resulting somatic haplotagged reads (Figure 1(d)). In a normal sample, the two germline haplotypes are nearly balanced across the genome. As tumor content rises, the somatic reads (HP1-1 or HP2-1) over-represent one haplotype by an amount proportional to tumor abundance. Because arm-level LOH imposes maximal imbalance regardless of tumor fraction, a regression on the haplotype-imbalance profile together with the LOH map yields the final DNA fraction.

LongPhase-TO yields a genome-wide, phase-resolved view that partitions long reads into parental germline haplotypes and their somatic descendants. In a tumor-only dataset (HCC1395 HKU), reads carrying a somatic variant from the Sequencing Quality Control Phase 2 (SEQC2) truth set were assigned to the somatic haplotype (HP2-1) and anchored to its parental lineage HP2, cleanly separated from HP1-/HP2-tagged germline reads (Figure 1(e)). Within an LOH region of HP1-deleted haplotype, LongPhase-TO continued to accurately tag somatic reads as HP2-1 rather than collapsing them to their ancestral haplotype (HP2) or misassigning them to HP1 (Figure 1(f)). Therefore, the somatic haplotyping by LongPhase-TO preserves the clonal ancestry and is robust to LOH in heterogeneous tumors.

### LongPhase-TO improves haplotype contiguity and delineates chromosome-scale LOH

We first benchmarked LongPhase-TO against three germline phasers, WhatsHap, HapCUT2 and LongPhase, across eight datasets from six cell lines representing breast cancer, melanoma and lung cancer (Supplementary Table 1). Tumor coverage ranged from 33 to 158*×* (median, 81 *×*). Across all datasets, LongPhase-TO produced markedly longer and less fragmented haplotypes than the germline phasers (Figure 2(a), Supplementary Table 2). LongPhase-TO increased block N50 in every dataset, by 1.1–48*×* (median, 2.9 *×*), to a maximum of 25.2 Mb in HCC1937_UCSC, and reduced the number of blocks 1.7–5.1-fold (median, 3.7-fold). The gain scaled with LOH burden, the fraction of the autosomal genome affected by LOH. It was smallest in HCC1954_UCSC, with 10% LOH and a 1.1*×* increase, and largest in the LOH-rich HCC1395 and HCC1937 lines, with 51–56% LOH and increases of up to 48*×* . The total length of phased blocks also increased, by 3–38% (median, 11%, Figure 2(b)), and the fraction of phased SNVs increased in seven of eight datasets (e.g., 0.40 to 0.71 in HCC1937_UCSC).

**Fig. 2.**
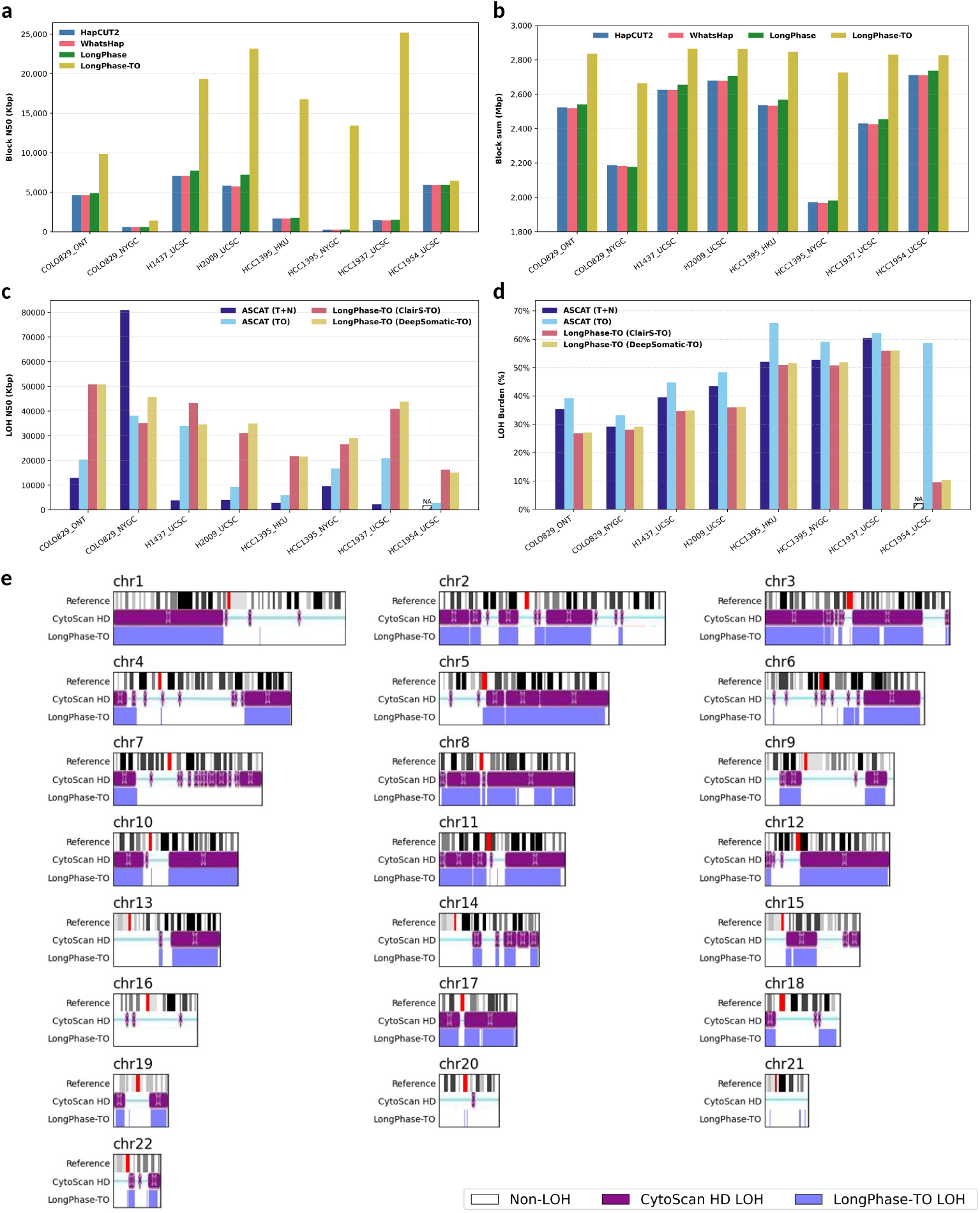
Comparison of LOH-aware phasing and LOH calling. (a) Haplotype block N50 for three germline-based phasing tools (HapCUT2, WhatsHap, LongPhase) and the LOH- and somatic-aware phaser LongPhase-TO across eight datasets from six cancer cell lines. (b) Total length of phased haplotypes for the same methods and datasets. (c) LOH N50 of ASCAT in tumor-normal (T+N) and tumor-only (TO) modes and of LongPhase-TO using somatic variants from ClairS-TO or DeepSomatic-TO. (d) LOH burden (fraction of the autosomal genome with LOH) of ASCAT (T+N), ASCAT (TO), and LongPhase-TO (ClairS-TO and DeepSomatic-TO). (e) Orthogonal comparison in HCC1395, the only cell line with an available independent LOH profile: chromosome ideograms show reference cytobands (top), LOH segments reported by CytoScan HD microarray (middle, purple), and LOH segments inferred by LongPhase-TO (bottom, blue).

We next compared arm-level LOH segments from LongPhase-TO with those from ASCAT, a CNV-based method [35]. ASCAT was run in tumor-only (TO) and tumor-normal (T+N) modes, and LongPhase-TO delineated LOH using somatic variants called by ClairS-TO or DeepSomatic-TO. Because independent LOH annotations were unavailable for most cell lines, these comparisons assess segmentation contiguity and cross-method agreement rather than accuracy against a ground truth. LongPhase-TO produced markedly more contiguous LOH segments than ASCAT (Figure 2(c), Supplementary Table 3). With either somatic caller, LongPhase-TO reached a median LOH N50 of 33–35 Mb, versus 18.6 Mb for ASCAT (TO) and 4.1 Mb for ASCAT (T+N). In HCC1937_UCSC, for example, LongPhase-TO reached 41–44 Mb, versus 20.9 Mb for ASCAT (TO) and 2.2 Mb for ASCAT (T+N). This advantage held in seven of eight datasets. The exception was COLO829_NYGC, where ASCAT (T+N) reached an N50 of 80.9 Mb. Consistent with reduced fragmentation at CNV/BFB events within LOH tracts, LongPhase-TO produced a median of 138–162 LOH segments, versus 558 for ASCAT (TO) and 1,948 for ASCAT (T+N). The two LongPhase-TO configurations gave similar per-dataset LOH N50 values (median difference, *∼*8%), indicating robustness to the choice of somatic caller.

Finally, we compared LOH burdens estimated by LongPhase-TO and ASCAT. LongPhase-TO burdens ranged from 10% in HCC1954_UCSC to 56% in HCC1937_UCSC and agreed to within *∼*1% between the two somatic callers (Figure 2(d)). ASCAT (T+N) reproduced this ranking of cell lines whereas ASCAT (TO) did not, and both modes reported systematically higher burdens (median, 43.4% for T+N and 53.5% for TO, versus 35% for LongPhase-TO with either caller). The largest discrepancy occurred in HCC1954_UCSC, where ASCAT (TO) estimated 58.7% and ASCAT (T+N) failed to converge, against 10% for LongPhase-TO. This discrepancy may reflect the whole-genome-duplication structure of HCC1954. For the two cell lines sequenced independently twice, COLO829 (ONT and NYGC) and HCC1395 (HKU and NYGC), LongPhase-TO burdens differed by at most 2% between the two runs, whereas ASCAT burdens differed by up to 6% in T+N mode and 7% in TO mode. In HCC1395, the LongPhase-TO chromosome ideograms recapitulated the arm-level LOH architecture of the published CytoScan HD microarray profile across the autosomes (Figure 2(e)) [37].

### Haplotype-aware recalibration improves somatic SNV and indel calls

To test whether somatic haplotyping can sharpen existing callers, we used LongPhase-TO to recalibrate the somatic SNVs of ClairS-TO and DeepSomatic-TO across eight tumor-only datasets, comparing each caller alone against the same caller followed by recalibration over an *in silico* dilution series spanning 20–100% tumor DNA fraction (see Methods). Averaged over all datasets and fractions, recalibration raised the mean SNV F1-score from 0.55 to 0.62 for ClairS-TO and from 0.55 to 0.65 for DeepSomatic-TO (Figure 3(a), Supplementary Table 4). The gain was consistently precision-driven. Mean precision rose by 0.15 for ClairS-TO at near-constant recall and by 0.24 for DeepSomatic-TO at a recall cost of 0.08, as expected for a filter that removes candidates lacking single-haplotype support.

**Fig. 3.**
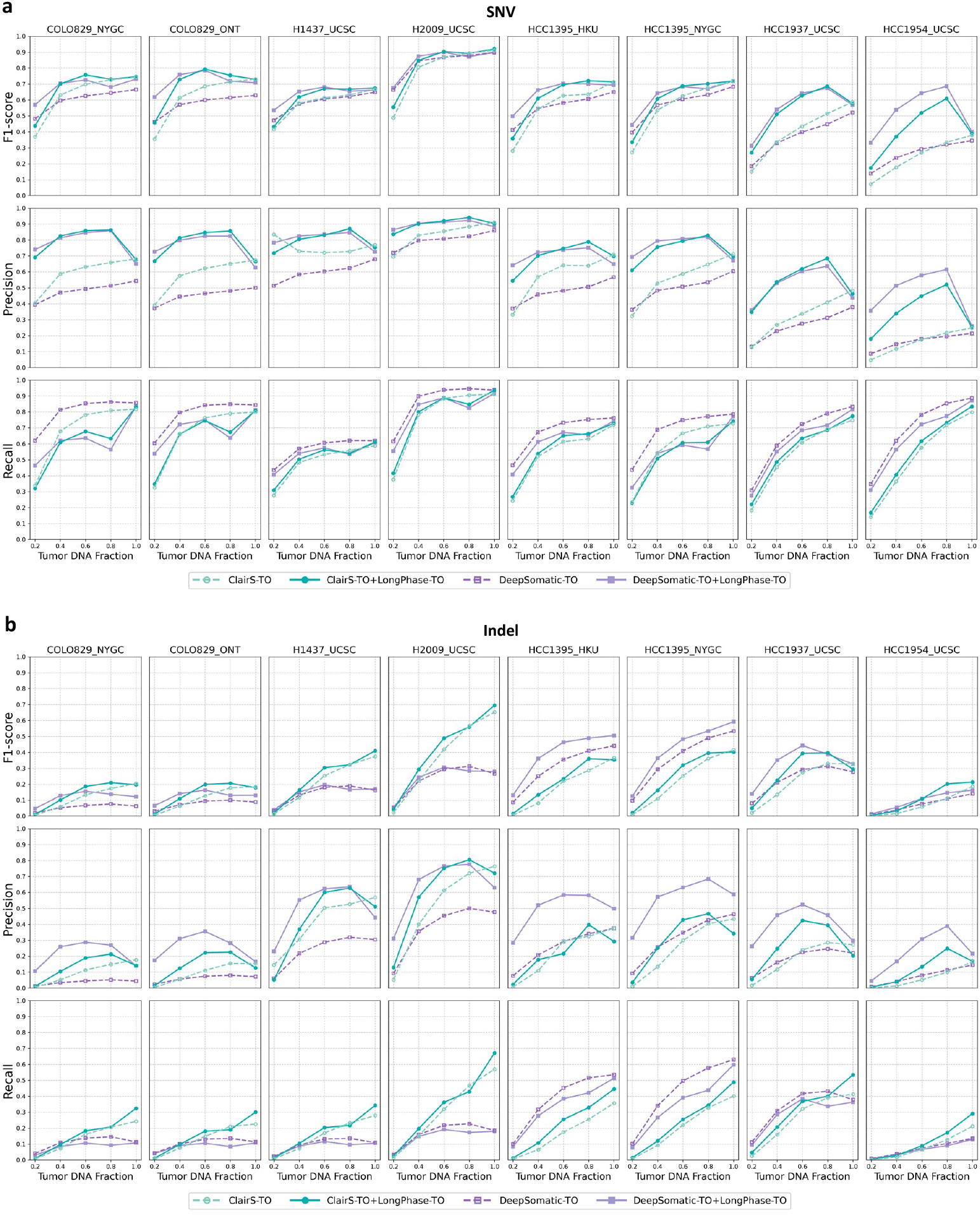
LongPhase-TO improves somatic SNV and indel calling accuracy across tumor DNA fractions. (a) F1-score, precision and recall of ClairS-TO and DeepSomatic-TO SNV calling, with or without LongPhase-TO recalibration, are shown for eight tumor cell line datasets across DNA fractions from 20% to 100%. LongPhase-TO consistently increases F1, with the largest gains at 20-80% DNA fraction by improving precision while maintaining similar or modestly reduced recall for ClairS-TO and DeepSomatic-TO. (b) F1-score, precision and recall of ClairS-TO and DeepSomatic-TO indel calling, with or without LongPhase-TO recalibration, are shown for eight tumor cell line datasets across DNA fractions from 20% to 100%.

We next examined whether the benefit of recalibration depended on tumor DNA fraction. The standalone F1-scores of ClairS-TO and DeepSomatic-TO fell monotonically as tumor DNA fraction decreased, and LongPhase-TO recalibration improved F1 at every level while blunting this decline (Figure 3(a)). Absolute F1 gains of 0.08– 0.15 were obtained across 20–80% DNA fraction. At 20% fraction, for instance, mean F1 rose from 0.30 to 0.38 for ClairS-TO and from 0.40 to 0.50 for DeepSomatic-TO, with the corresponding precision gain over this range averaging *∼*0.18 and *∼*0.28, respectively. Although the average benefit was greatest at low-to-intermediate tumor DNA fractions, large improvements were also observed in difficult individual datasets at higher fractions. In HCC1954_UCSC at 80% DNA fraction, recalibration raised F1 from 0.33 to 0.61 for ClairS-TO and from 0.32 to 0.69 for DeepSomatic-TO. At 100% DNA fraction, where the standalone callers were already strongest, the benefit diverged between callers: mean F1 was essentially unchanged for ClairS-TO (0.68 to 0.68) but still improved for DeepSomatic-TO (0.63 to 0.67), consistent with recalibration acting mainly where the baseline caller is least precise. The same fraction-dependent profile was reproduced with a ClairS-TO model trained on simulated data (ClairS-TO-ss, Supplementary Fig. 1).

We next asked whether recalibration offers comparable gains for somatic indels. Indels were substantially harder to detect than SNVs, with a baseline mean F1 of only 0.19 for both callers versus 0.55 for SNVs (Figure 3(b), Supplementary Table 5), a gap inherited from the lower baseline accuracy of both callers on long-read indels. Recalibration nonetheless improved mean indel F1 to 0.23 for each caller, again through precision (0.23 to 0.28 for ClairS-TO; 0.19 to 0.41 for DeepSomatic-TO), with recall changing by less than 0.04 in either direction (a small gain for ClairS-TO and a small loss for DeepSomatic-TO). As with SNVs, the indel gains concentrated at low-to-intermediate fraction. In HCC1937_UCSC at 60% fraction, for example, F1 rose from 0.27 to 0.39 for ClairS-TO and from 0.29 to 0.44 for DeepSomatic-TO. Even at 100% fraction, mean indel F1 stayed above baseline for both callers.

Because tumor and admixed normal cells differ in CpG methylation [29, 38, 39], a somatic alternate allele carries a single methylation state while the reference allele mixes both (see Methods). LongPhase-TO can optionally exploit this allele-specific 5-methylcytosine (5mC) asymmetry as a second signal beyond haplotype linkage. We tested its incremental value on the five datasets with methylation calls, comparing for each caller the original calls, haplotype-aware recalibration alone, and recalibration with the added methylation-aware step (Supplementary Figs. 2 and 3, Supplementary Table 6). Relative to haplotype-aware recalibration alone, the methylation-aware step raised mean precision from 0.68 to 0.72 for SNVs and from 0.40 to 0.49 for indels, each at a mean recall cost of 0.01, leaving F1 nearly unchanged (0.01 for SNVs and negligible for indels). The precision gain was nonetheless consistent, improving 34 of 50 caller-dataset-fraction combinations for SNVs and 31 of 50 for indels, with largest gains of 0.14 in HCC1937 UCSC. Across datasets and variant types, the gain increased as haplotype-aware precision decreased, indicating that allele-specific methylation compensates for weak haplotype evidence.

### Somatic haplotagging estimates tumor DNA fraction across dilution series

LongPhase-TO derives tumor DNA fraction, the fraction of sequenced molecules of tumor origin, from the haplotagged reads produced by phasing (see Methods), whereas ASCAT and PURPLE infer purity jointly with ploidy from allele-specific copy-number profiles (Supplementary Table 7), and their purities were converted to DNA fraction with the fitted ploidy for comparison (see Methods). We evaluated the haplotagging-based estimator on *in silico* dilution series of eight nanopore datasets at five DNA fractions each, with somatic variants called by DeepSomatic-TO or ClairS-TO. As comparators we ran PURPLE in tumor-only mode and ASCAT in both tumor-only (TO) and tumor-normal (T+N) modes.

Across the panel, LongPhase-TO recovered the designed DNA fraction with a mean absolute error of 0.04 using DeepSomatic-TO variants and 0.06 using ClairS-TO variants, and with Lin’s concordance correlation coefficients of 0.96 and 0.90 (Figure 4(c), Supplementary Tables 8 and 9). Estimates fell within 0.05 of the achieved fraction in 29 and 27 of 40 mixtures and within 0.10 in 35 and 34, and the replicate datasets of COLO829 and HCC1395 from different sequencing centers gave concordant estimates. PURPLE was comparably accurate (mean absolute error 0.08, concordance 0.79). Although PURPLE and LongPhase-TO share no modeling assumptions, their estimates for the same mixture differed by a median of 0.05 with either variant caller. ASCAT was markedly less accurate in tumor-normal and tumor-only modes (mean absolute error 0.20 and 0.33, concordance 0.24 and *−*0.11) and returned no estimate for 7 of 40 mixtures. The DNA fraction obtained as a by-product of phasing thus matched the accuracy of estimators that fit a full copy-number and ploidy model.

**Fig. 4.**
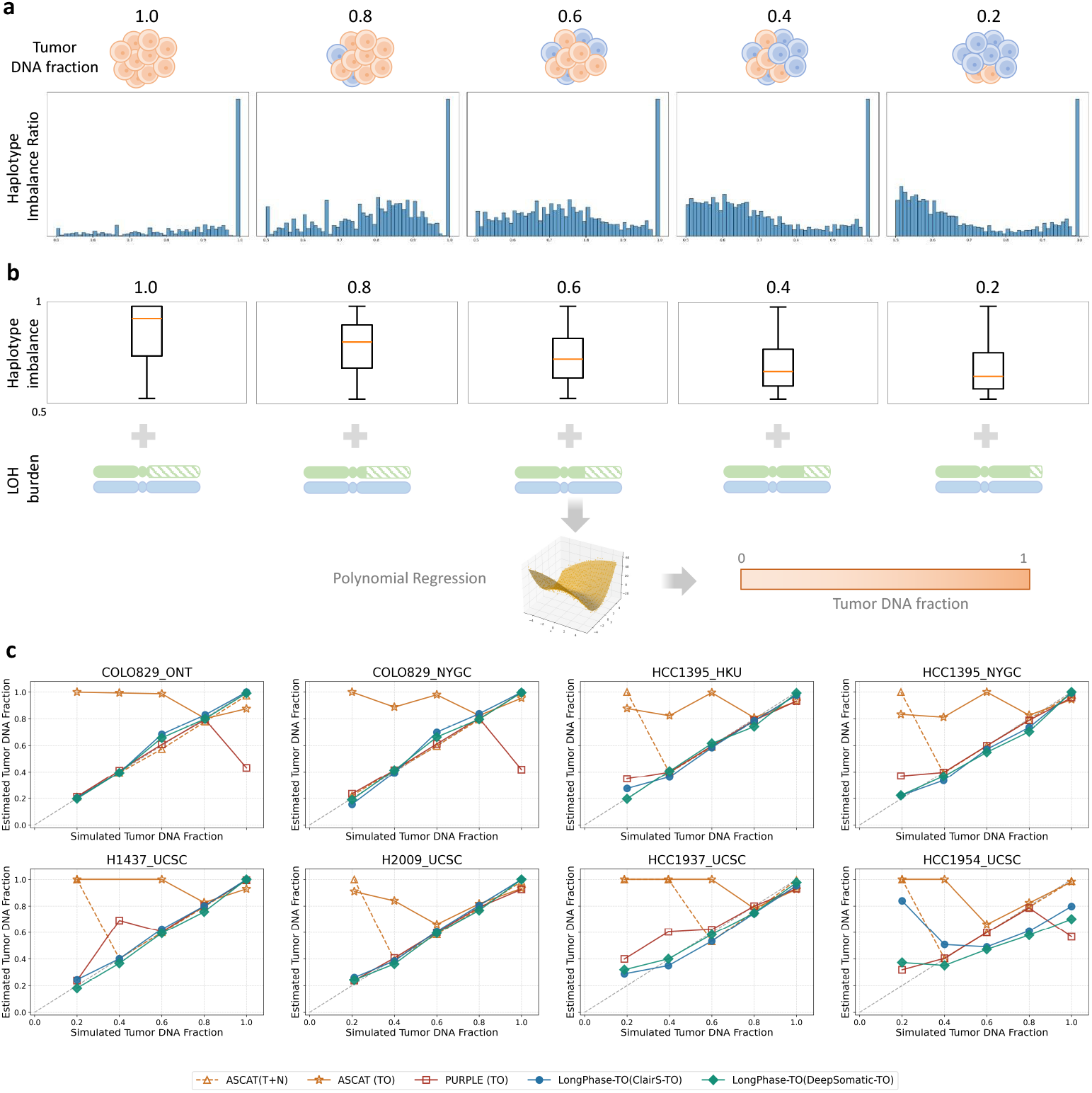
Tumor DNA fraction estimation by LongPhase-TO across in-silico dilution series. (a) Genome-wide distribution of the haplotype imbalance ratio at simulated tumor DNA fractions from 1.0 to 0.2. As DNA fraction decreases, the distribution shifts away from 1.0 toward the balanced value of 0.5. (b) Summary statistics of the haplotype imbalance distribution (boxplots) together with the LOH burden are used as features in a polynomial regression to estimate tumor DNA fraction. (c) Estimated DNA fraction (y-axis) versus simulated DNA fraction (x-axis) for ASCAT (T+N), ASCAT (TO), PURPLE (TO), LongPhase-TO (ClairS-TO), and LongPhase-TO (DeepSomatic-TO) across eight cancer cell line datasets; the diagonal indicates the ideal prediction (*y* = *x*). LongPhase-TO tracks the designed dilution across the whole range without a copy-number or ploidy model, whereas ASCAT overestimates DNA fraction at low tumor content and PURPLE underestimates it in the near-pure samples.

The errors of the three methods concentrated in different dilution regimes and cell lines (Supplementary Tables 8 and 10). Across dilution levels, the error of ASCAT ranged from 0.01 to 0.65 in tumor-normal mode and from 0.05 to 0.79 in tumor-only mode, peaking in the most dilute mixtures (DNA fraction 0.2). The error of PURPLE ranged from 0.01 to 0.23, peaking in the pure samples (DNA fraction 1.0). The error of LongPhase-TO ranged from 0.02 to 0.12 with either variant caller, with no level standing out. Across cell line datasets, the error of ASCAT ranged from 0.03 to 0.70 in tumor-normal mode and from 0.25 to 0.55 in tumor-only mode, peaking in HCC1954_UCSC. The error of PURPLE ranged from 0.02 to 0.13, peaking in the two COLO829 datasets. The error of LongPhase-TO ranged from 0.01 to 0.25 with either variant caller, also peaking in HCC1954_UCSC (0.17 and 0.25 for DeepSomatic-TO and ClairS-TO). This cell line carries a whole-genome duplication and had the lowest LOH burden in the panel, the condition under which haplotype imbalance carries the least information (see Discussion).

### Variant recalibration and tumor DNA fraction estimation in PacBio HiFi data

Both trained components of LongPhase-TO, the triplet-graph coefficients and the DNA fraction regression, were fitted once on nanopore data and held fixed. To test whether either depends on the sequencing platform, we applied them without refitting to six PacBio HiFi datasets of the same six cell lines [15] (see Methods), with ClairS-TO and DeepSomatic-TO run with their PacBio HiFi models (see Methods). Recalibration raised mean SNV F1 from 0.58 to 0.67 for ClairS-TO and from 0.60 to 0.69 for DeepSomatic-TO, and mean indel F1 from 0.24 to 0.27 and from 0.28 to 0.32 (Supplementary Figs. 4 and 5, Supplementary Tables 11 and 12). F1 improved at 25 and 24 of 30 SNV points and at 26 and 23 of 30 indel points, with gains again concentrated at low-to-intermediate DNA fraction. Every decrease larger than 0.01 occurred at 0.8 or 1.0 DNA fraction, the largest for DeepSomatic-TO on the pure H2009 sample (SNV F1 from 0.88 to 0.31, indel F1 from 0.39 to 0.18), the condition furthest from the 0.6 DNA fraction at which the coefficients were fitted.

The DNA fraction regression transferred equally well. With its coefficients unchanged, LongPhase-TO gave mean absolute errors of 0.06 and 0.08 for the DeepSomatic-TO and ClairS-TO configurations on the PacBio panel, against 0.04 and 0.06 on nanopore, with concordance coefficients of 0.94 and 0.89 (Supplementary Fig. 6, Supplementary Tables 9 and 13). PURPLE was again comparably accurate (mean absolute error 0.07, concordance 0.87) and again peaked in the pure samples, and ASCAT (mean absolute error 0.09 and 0.24 in tumor-normal and tumor-only modes, concordance 0.59 and *−*0.04) again peaked in the most dilute mixtures. HCC1954 remained the least accurate dataset for LongPhase-TO, with mean absolute errors of 0.18 and 0.29 against 0.17 and 0.25 on nanopore. Both components therefore transfer across long-read platforms without platform-specific retraining.

## Discussion

LongPhase-TO represents parental germline haplotypes and their somatic descendants within one LOH-aware graph. Across eight nanopore cell-line datasets, this shared representation yielded longer and less fragmented haplotypes than germline phasers, delineated contiguous arm-level LOH, and improved somatic SNV and indel calls through precision-driven recalibration. It also enabled tumor DNA fraction to be inferred from germline haplotype imbalance without a matched normal sample. Both trained components transferred to six PacBio HiFi datasets without refitting. Concordance with CytoScan HD supported the biological plausibility of the LOH calls in HCC1395. Agreement across somatic callers and independently sequenced datasets established technical reproducibility across the remainder of the panel.

These results distinguish LongPhase-TO from methods that consume haplotypes rather than construct them. Germline phasers such as WhatsHap, HapCUT2 and LongPhase partition reads between two inherited haplotypes [8–10]. Extended LOH removes the heterozygous markers that connect those haplotypes and fragments phase. The long-read cancer suites SAVANA, Wakhan and TumorLens inherit this limitation because they take phase from germline phasing and describe somatic events relative to the resulting germline haplotypes [32–34]. LongPhase-TO instead makes the somatic alleles nodes of the phasing graph, tracks them as descendants of their parental haplotypes, and reconciles chromosome-scale LOH before graph traversal, thereby maintaining phase continuity across LOH regions.

The same local haplotype structure provides a biological constraint for refining calls from ClairS-TO and DeepSomatic-TO [15, 24]. A genuine somatic allele remains confined to one parental lineage. Residual germline alleles and sequencing errors are less likely to satisfy this constraint. The reproducible precision gains across both callers show that this haplotype evidence complements caller-specific models. Allele-specific CpG methylation supplies a second constraint of the same kind, because a somatic allele arises in tumor cells whose methylation state differs from that of admixed normal cells [29, 38, 39], and its contribution was largest where haplotype evidence was weakest. Precision-driven recalibration has a cost near full tumor content. The triplet-graph coefficients were fitted on a single mixture at 0.6 DNA fraction, where admixed normal reads populate the parental haplotype alongside its somatic descendant. In a pure clonal sample no reads carry the parental haplotype without the somatic allele, and the triplet collapses to a pair. On the nanopore panel the mean gain at 1.0 DNA fraction shrank to zero for ClairS-TO. On the PacBio panel every F1 decrease larger than 0.01 occurred at 0.8 or 1.0 DNA fraction, most severely on the pure H2009 sample. Because LongPhase-TO estimates the DNA fraction itself, that estimate could gate the recalibration or select fraction-specific coefficients.

The tumor DNA fraction reported by LongPhase-TO is not tumor cellular purity. ASCAT [35], PURPLE [36] and Battenberg [40] model aneuploidy and copy-number alterations explicitly and report cellular purity. LongPhase-TO estimates the proportion of sequenced DNA molecules of tumor origin from germline haplotype imbalance alone. The two quantities are similar in near-diploid genomes with limited copy-number change. They diverge under widespread aneuploidy or whole-genome duplication, because tumor and normal cells then contribute different amounts of DNA per cell. All comparisons in this study were made in DNA fraction (see Methods).

The HCC1954 series highlights a limitation of this haplotype-based estimator. At phased somatic sites, imbalance among reference-supporting reads can increase with tumor DNA fraction even without LOH; LOH and allele-specific copy number can modify this relationship. HCC1954 carries a whole-genome duplication [41] and had the lowest LOH burden in the panel. Loss of one chromosomal copy from the duplicated genome leaves an imbalanced but still heterozygous 2:1 state rather than eliminating a parental allele, and extensive aneuploidy coexists with little true LOH [42]. The estimates were compressed at high DNA fraction, reaching only 0.65 to 0.80 in the pure samples across both variant callers and both sequencing platforms, and the ClairS-TO configuration also overestimated the most dilute mixture by 0.4 or more (Supplementary Tables 8 and 13). These errors may reflect the effects of genome duplication and allele-specific copy number on the haplotype-imbalance profile. PURPLE, which models copy number explicitly, tracked the intermediate mixtures of this genome closely. Whether LOH burden can serve as an indicator of estimation confidence requires validation.

Copy-number estimators have a boundary of their own. A purity and ploidy fit can be weakly identified as tumor DNA fraction approaches 1.0, because some copynumber profiles admit both a high-purity solution and a lower-purity solution at a multiple of the ploidy [35, 36]. The largest errors of PURPLE occurred in the pure mixtures, with estimates near half the expected fraction, and recurred for the same genome in independently sequenced datasets (Supplementary Tables 8 and 13). This pattern is consistent with the selection of an alternative purity and ploidy solution. LongPhase-TO instead estimates DNA fraction from haplotype imbalance and LOH burden without explicitly fitting ploidy. The contrasting error profiles observed for the two approaches suggest that haplotype and copy-number information could provide complementary constraints on tumor DNA fraction estimation. Allele-specific copynumber modeling could stabilize the haplotype-based estimate in genome-doubled, low-LOH samples, and the haplotype-imbalance signal could in turn constrain copynumber-based estimators near full tumor content. SAVANA and Wakhan, which already fit such models without a matched normal [32, 33], are candidate frameworks for this integration.

Several limitations bound these conclusions. The benchmarks are based on cell-line dilution series and do not establish performance in primary tumors, highly subclonal lesions or diagnostic specimens. The results establish methodological feasibility and technical accuracy under controlled conditions. Validation of the LOH calls is further limited by the scarcity of independent chromosome-scale references. A published orthogonal profile was available only for HCC1395. The remaining datasets establish contiguity and technical reproducibility rather than biological accuracy across distinct tumor genomes. Primary tumors with matched allele-specific array or sequencing references, complemented by controlled simulations with known LOH boundaries, will be required to establish generalizability across event sizes, coverage levels and tumor DNA fractions.

## Methods

Before LOH-aware somatic phasing, LongPhase-TO independently recalibrates somatic variant candidates by a triplet graph model and detects arm-level LOH by breakpoint analysis. The resulting somatic call set and LOH map are then integrated for phasing, after which tumor DNA fraction is estimated (Figure 1).

### Triplet-graph recalibration of somatic variants

Recalibration of somatic variant candidates from tumor-only long-read data proceeds in two steps: candidate enrichment by filtering, followed by triplet-graph evidence modeling that exploits local haplotype linkage (Figure 1(a)).

### Somatic candidate filtering using external resources and allele concordance

In the absence of a matched normal sample, LongPhase-TO screens candidate variants against four external germline and normal-panel resources adopted from the ClairS-TO workflow. The 1000 Genomes panel of normals (PoN) and CoLoRSdb GRCh38 v1.1.0 sites with allele frequency *≥*0.001 are queried by genomic position, whereas dbSNP build 138 non-somatic sites and gnomAD r2.1 sites with allele frequency *≥*0.001 require concordance of both position and alternate allele. Candidates matching either set are excluded from somatic classification before triplet-graph recalibration.

Systematic sequencing and alignment artifacts can generate clusters of nearby non-reference observations on the same reads. LongPhase-TO therefore assesses position-level allele concordance within 100 aligned positions on either side of each candidate *i*. Let *N*_*A*_(*i*) denote the number of reads supporting the candidate alternate allele, and let *n*_*A*_(*i, j*) and *n*_*R*_(*i, j*) denote the numbers of candidate-alternate and candidate-reference reads, respectively, carrying a non-reference observation at neighboring position *j*. Position *j* is considered concordant with the candidate when

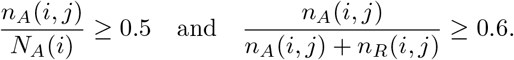

Thus, the non-reference observation at *j* must occur in at least 50% of candidate-alternate reads, with at least 60% of reads carrying that observation also supporting the candidate alternate allele. A candidate is removed as a clustered artifact when at least two distinct neighboring positions satisfy both criteria. This procedure is based on the recurrence position rather than the identity of the non-reference nucleotide, and all thresholds were fixed and applied unchanged across datasets.

### Candidate evaluation with the triplet-graph model

#### Triplet-graph construction and pattern classification

A somatic allele arises on one parental haplotype and is subsequently inherited by descendant tumor subclones (Supplementary Fig. 7(a)). In contrast, sequencing and alignment artifacts may generate apparent alternate alleles but generally lack consistent linkage to a parental haplotype. LongPhase-TO leverages this local haplotype context using a triplet graph spanning three variant positions (Supplementary Fig. 7(b)). Nodes represent reference or alternate alleles, and weighted paths record the observed three-variant haplotypes and their read support. Each locus can carry a reference (R) or alternate (A) allele, giving eight possible paths in a triplet. A true somatic mutation is expected to produce two parental germline paths and a third path that diverges from one parent at the candidate allele. Each candidate-containing triplet contributes at most one of three mutually exclusive evidence categories: *V*_*H*_ denotes high-confidence three-dominant evidence comprising two parental germline paths and one somatic-descendant path; *V*_*L*_ denotes a lower-confidence two-dominant-path pattern that remains compatible with a single haplotype-specific somatic origin; and *V*_*N*_ denotes any other topology accepted as neither *V*_*H*_ nor *V*_*L*_. These categories showed distinct enrichment among validated true- and false-positive calls (Supplementary Fig. 8). The details are given below.

#### Regression-based scoring of triplet patterns

For each triplet, the read supports of the eight possible paths are ranked as *r*_1_ *≥ r*_2_ *≥ r*_3_. LongPhase-TO first evaluates 12 predefined high-confidence (*V*_*H*_) topologies, comprising four evolutionary configurations for each placement of the somatic candidate at the left, middle or right node (Supplementary Fig. 7(c)). The read supports of the two parental paths (*x* and *y*) and the somatic-descendant path (*z*) are scored by two pre-trained models: a linear predictor for indels and a third-order polynomial predictor in *x, y* and *z* for SNVs (see Supplementary Methods). Both models were fitted on the HCC1395_HKU mixture at tumor DNA fraction and then held fixed for all other datasets and DNA-fraction levels. An eligible topology contributes a *V*_*H*_ vote when its model-estimated probability is *≥*0.5. If no *V*_*H*_ configuration is accepted for that triplet, LongPhase-TO further considers lower-confidence, two-dominant-path configurations compatible with a haplotype-specific somatic origin. In LOH or locally homozygous regions, a somatic mutation is expected to produce two dominant paths, although admixed normal DNA may contribute a weak third path. LongPhase-TO therefore evaluates nine *V*_*L*_ topologies that accommodate this residual signal (*r*_3_ *<* 0.5*r*_2_), while candidate-level voting limits the associated loss of specificity. The weak third path must be either the all-reference path (RRR) or one of two paths that bypass the candidate somatic allele (Supplementary Fig. 7(d)). If neither *V*_*H*_ nor *V*_*L*_ is accepted, the triplet contributes a *V*_*N*_ vote when *r*_2_ *≥*2 (Supplementary Fig. 7(e)). Other triplets with less read support are not considered.

#### Candidate-level integration of triplet evidence

The hierarchical classification described above is applied separately to every candidate-containing triplet. A candidate is evaluated in the triplets in which it occupies the left, middle or right node; with each edge spanning up to *k* neighboring variants, the two accompanying loci generate up to *k*^2^ triplets per placement and 3*k*^2^ in total. Different triplets for the same candidate may therefore contribute *V*_*H*_, *V*_*L*_ or *V*_*N*_ votes, even though each individual triplet contributes at most one vote. Let *H, L* and *N* denote the numbers of accepted *V*_*H*_, accepted *V*_*L*_ and non-somatic votes across these triplets. For each accepted *V*_*H*_ triplet *g* we record the somatic-path proportion *s*_*g*_, defined as the support for the selected somatic-descendant path relative to the total support for the paths carrying the candidate alternate allele, and summarize the high-confidence evidence by its mean *S*_*H*_ over the *H* accepted triplets. The candidate is retained through either the high-confidence or low-confidence evidence route:

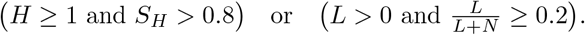

The two thresholds were fixed from the corresponding score distributions in the training sample (Supplementary Figs. 9 and 10; Supplementary Methods). Mutation-type-specific regression thus governs whether an individual triplet contributes evidence, whereas the site-level decision integrates the accepted high- and low-confidence evidence across the local graph ensemble.

### Methylation-aware recalibration of somatic variants

When the input alignments carry 5-methylcytosine (5mC) tags, LongPhase-TO applies an optional recalibration of the candidates retained by the triplet-graph model that exploits allele-specific CpG methylation; the step is skipped, and the triplet-graph output returned unchanged, for data without methylation calls. Because a somatic variant arises within a tumor subpopulation whose CpG methylation state differs from that of admixed normal cells, reads carrying its alternate allele tend to share a single methylation state, whereas reads carrying the reference allele mix tumor- and normal-derived molecules; germline variants, present in both compartments, produce methylation heterogeneity on both alleles, and sequencing or alignment artifacts show no consistent allelic asymmetry. For both somatic SNV and indel candidates, LongPhase-TO summarizes this signal with ten features computed from the CpG sites flanking each candidate, spanning three complementary views of the methylation neighborhood: allelic methylation coverage, joint haplotype methylation proportions and methylation-cloud geometry (Supplementary Figs. 11–14). A gradient-boosted decision-tree classifier (XGBoost), selected through held-out comparisons with six alternative classifiers (Supplementary Fig. 15), uses these features to label each candidate as somatic, germline or artifact, retaining only candidates classified as somatic. The classifier was trained once on the HCC1395_NYGC mixture at 0.6 tumor DNA fraction and held fixed thereafter (Supplementary Methods). This step’s incremental effect on somatic SNV and indel calling accuracy, beyond haplotype-aware recalibration alone, is evaluated in Results (Supplementary Figs. 2 and 3).

### Arm-level LOH detection

LongPhase-TO detects arm-level LOH, defined as the loss of one parental allele across a large genomic region, from the depletion of heterozygous sites. Small copy-number variants (CNVs) and breakage–fusion–bridge (BFB) events embedded within an LOH tract can locally restore heterozygosity and fragment a broad LOH call. To reduce this fragmentation, LongPhase-TO identifies the corresponding short intervals from read-clipping signals and omits variants within them when calculating regional heterozygosity. The remaining breakpoint signals partition the chromosome into regions that are classified as LOH or non-LOH according to their heterozygosity ratios (Supplementary Fig. 16).

### Directional clipping and genomic-event interval definition

Directional clipping is used to distinguish localized breakpoint-associated interruptions from boundaries delimiting broader genomic regions (Supplementary Fig. 17). Soft- and hard-clipping operations longer than five bases are counted separately at the left and right alignment ends. At the ordered clip-bearing position *i, F*_*i*_ and *B*_*i*_ denote the numbers of these qualifying clipping operations at the front (left) and back (right) alignment ends, respectively; they are alignment-end counts, not forward- and reverse-strand read counts. Their difference, *d*_*i*_ = *F*_*i*_*− B*_*i*_, defines the directional clipping signal. Breakpoints of small CNV events often produce concentrated clipping support at a single genomic position and are therefore retained directly. By contrast, the complex junctions generated by BFB events can distribute clipping support across several nearby positions, producing lower-amplitude signals that may not satisfy a single-position threshold. To recover these dispersed BFB-associated boundaries, LongPhase-TO aggregates weak clipping signals of the same orientation across neighboring positions using rolling windows applied in both genomic directions. Peaks in the aggregated profiles define candidate boundaries, and each candidate is assigned a support value by comparing the smoothed cumulative clipping signal on its two sides; the sign of this change preserves the breakpoint orientation. Full signal-processing definitions are provided in Supplementary Methods (Section 2.1.2).

Candidate clipping signals are then paired according to their orientation and genomic proximity. Nearby signals of opposite orientation are paired to delimit short intervals associated with local CNV or BFB events, whereas unpaired signals are retained as candidate boundaries of broader LOH regions. Variants within these short CNV/BFB-associated intervals are omitted during heterozygosity calculation, whereas the candidate LOH boundaries define the regions evaluated. Detailed pairing rules and fixed parameter values are provided in Supplementary Methods (Section 2.1.2) and Supplementary Table 14; all parameters were applied unchanged across datasets.

### LOH classification after exclusion of small-event intervals

For each region bounded by consecutive candidate LOH boundaries, variants falling within a short CNV/BFB-associated interval are omitted before heterozygosity calculation. Each remaining variant is counted as homozygous when its variant allele fraction (VAF) is at least 0.8 or when it is marked as homozygous in the input; all other variants are counted as heterozygous. Let *N*_het_ and *N*_hom_ denote the resulting counts. The regional heterozygosity ratio is

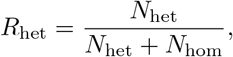

and a region is classified as LOH when *R*_het_ *<* 0.09. The VAF and *R*_het_ thresholds were selected from their empirical distributions within the HCC1395 LOH and non-LOH reference intervals published by SEQC2 [37] (Supplementary Fig. 18; Supplementary Methods, Section 2.1.1). Both thresholds were then fixed and applied unchanged across all sequencing datasets. Because variants within short CNV/BFB-associated intervals are excluded before this calculation, localized restoration of heterozygosity associated with small CNV or BFB events does not split the encompassing chromosome-scale LOH call (Supplementary Fig. 16).

### LOH-aware somatic phasing

#### Joint reconstruction of germline and somatic haplotypes

LongPhase-TO jointly phases recalibrated germline and somatic variants using a local allele-linkage graph (Supplementary Fig. 19). Each allele is represented as a node, and alleles observed on the same read are connected across a fixed number of downstream informative variants. Edge support reflects both the number and quality of the contributing read observations. For each pair of variants, LongPhase-TO compares evidence for parallel linkage (reference–reference and alternate–alternate) with that for crossed linkage (reference–alternate and alternate–reference). Within each phase block, HP1 and HP2 are initialized with an arbitrary orientation at the first informative variant, and weighted linkage evidence from preceding variants determines the phase of each subsequent variant. Ambiguous evidence initiates a new phase block rather than forcing an unsupported connection.

Somatic candidates are phased according to the linkage of their alternate alleles with the surrounding germline haplotypes. A somatic alternate allele supported by reads assigned to one parental haplotype is represented as a derived somatic haplotype, for example, HP2-1 descending from HP2, whereas candidates without sufficient haplotype-specific support remain unresolved. After this initial graph traversal, reads are assigned to HP1 or HP2 according to their agreement with the provisional phase, and high-confidence read assignments are used to refine the allele phase calls.

At an LOH boundary, linkage between heterozygous variants outside the LOH segment and homozygous variants within it is used to identify the parental haplotype retained in the LOH region. The retained haplotype and its somatic descendants are then propagated through the segment, whereas the lost parental path is recorded as absent. Linkage at the distal boundary reconnects the retained path to the subsequent diploid region, thereby preserving phase continuity across chromosome-scale LOH (Supplementary Fig. 19). Exact graph ranges, weighting rules and confidence thresholds are provided in Supplementary Methods (Section 2.3).

### Tumor DNA fraction estimation from haplotype imbalance

Using the somatic haplotagged reads from the phasing stage, LongPhase-TO estimates the tumor DNA fraction, defined as the proportion of sequencing reads originating from tumor cells, directly from germline haplotype imbalance (HI) at phased somatic variants (Figure 1(d)). At low tumor DNA fraction, reference-supporting reads at these sites are distributed approximately evenly between the two germline haplotypes, giving HI values near 0.5. As the tumor DNA fraction increases, these reads become progressively concentrated on one parental haplotype and the HI distribution tends to shift toward 1.0. At each phased somatic variant *i* with at least one informative reference-supporting read, HI is defined as

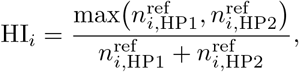

where 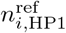 and 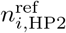 are the numbers of reference-supporting reads assigned to HP1 and HP2, respectively, at variant *i*. The site-level HI values are aggregated into a genome-wide distribution that forms the basis of the regression model below (Figure 4(a,b)).

The regression model uses three genome-wide features: the first and third quartiles (Q1 and Q3) of the site-level HI distribution and the LOH burden. Q1 and Q3 jointly summarize the shift and dispersion of the HI distribution. The LOH burden, defined as the total length of detected LOH regions divided by the total analyzed chromosome length, summarizes the genome-wide extent of parental haplotype loss and complements the site-level HI measurements.

The three features enter a complete third-degree polynomial regression containing all 20 monomials of total degree at most three. The resulting estimate is restricted to the interval [0, 1] (Figure 4(b)). Model coefficients were fitted using seven of the eight nanopore datasets, with HCC1954 UCSC reserved for independent testing. After fitting, the coefficients were fixed and applied without caller-specific refitting to somatic variants supplied by either ClairS-TO or DeepSomatic-TO, and without platform-specific refitting to the PacBio HiFi panel; the complete model is provided in Supplementary Methods (Section 2.4).

The estimator was compared with ASCAT in tumor-only and tumor-normal modes and with PURPLE in tumor-only mode, using the command lines given in the Supplementary Information. Both comparators report cellular purity together with a fitted tumor ploidy, whereas LongPhase-TO estimates tumor DNA fraction. To place all methods on the same axis, the cellular purity *p* reported by each comparator was converted to tumor DNA fraction *f* using its own fitted ploidy *κ*, assuming diploid normal-cell contamination:

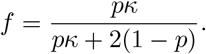

All ASCAT and PURPLE values reported in this study are these converted DNA fractions. Accuracy is reported as the mean absolute error against the achieved read-level mixing fraction, together with Lin’s concordance correlation coefficient, which unlike a Pearson correlation is sensitive to bias as well as to scale. Linearity over the dilution range is summarized by the slope and intercept of an ordinary least-squares fit of the estimate on the achieved fraction. Because the five DNA-fraction levels within a dataset are downsampled from a single library and are therefore not independent, the unit of analysis is the cell-line dataset, and the sensitivity of each summary statistic to individual cell lines is reported as the range obtained when any one dataset is excluded.

### High-DNA-fraction phasing refinement

The VAF of a heterozygous somatic variant increases with tumor DNA fraction, whereas that of a heterozygous germline variant remains near 0.5. At high tumor DNA fraction, overlap between these distributions reduces the reliability of the initial haplotype-based candidate recalibration (Supplementary Fig. 20).

LongPhase-TO therefore uses the tumor DNA fraction estimated from the initial phasing result to determine whether a refinement pass is required. When the estimate exceeds 0.9, candidate variants not flagged by the external germline or normal-panel resources are treated as somatic, and phasing is repeated once using the existing LOH map. Otherwise, the initial phasing result is retained. This additional pass refines phasing only; the tumor DNA fraction is not re-estimated (Supplementary Fig. 21).

### Benchmark datasets and experimental design

We analyzed eight publicly available nanopore long-read datasets spanning six cancer cell lines (HCC1395, COLO829, HCC1937, HCC1954, H1437 and H2009) sequenced by the University of Hong Kong (HKU), the New York Genome Center (NYGC), Oxford Nanopore Technologies (ONT) and the University of California, Santa Cruz (UCSC) (Supplementary Table 1). For somatic variant benchmarking, the HCC1395 truth set was obtained from SEQC2 [37], the COLO829 truth set from NYGC [43], and the HCC1937, HCC1954, H1437 and H2009 truth sets from Google DeepSomatic [15]. Tumor coverage ranged from 33.45*×* to 158.44*×* and matched normal coverage from 25.68*×* to 49.28*×* ; normal data were used only to generate the *in silico* tumor-normal mixtures and to run ASCAT in tumor-normal mode as a comparator.

LongPhase-TO is compared with the long-read somatic variant callers listed in Supplementary Table 15. We generated *in silico* mixtures by downsampling high-coverage reads from each pure tumor cell line and its matched normal and combining them at a fixed total coverage of 50*×* (for example, 40*×* tumor plus 10*×* normal for a 0.8 fraction). The target tumor DNA fractions were 1.0, 0.8, 0.6, 0.4 and 0.2, with the pure tumor cell line assigned a fraction of 1.0. Where the available coverage prevented a target from being met exactly, the achieved fraction was used in place of the nominal one (0.1875 to 0.2105 for the 0.2 target, 0.3953 to 0.3958 for the 0.4 target and 0.8049 for one 0.8 target). These achieved fractions served as ground truth for evaluating variant calling and for training and validating the tumor DNA fraction model.

To assess cross-platform generalization of the somatic recalibration, we additionally analyzed six PacBio Revio HiFi datasets (Supplementary Table 16), using the same dilution design, the same benchmark truth sets, and the same evaluation procedure. Five of the cell lines (HCC1395, HCC1937, HCC1954, H1437 and H2009) belong to the tumor-normal cohort assembled for DeepSomatic [15], four of them obtained from the CASTLE release and HCC1395 from the public PacBio Revio 2023Q2 release; COLO829 is an independent PacBio Revio benchmark from the same public release. ClairS-TO and DeepSomatic-TO were run with their PacBio HiFi models, whereas every trained LongPhase-TO model was that fitted on the nanopore panel and was applied without refitting or platform-specific tuning: the triplet-graph coefficients and site-level thresholds for somatic recalibration, and the polynomial coefficients of the tumor DNA fraction regression. Both components were therefore evaluated off their training platform. The corresponding command lines are given in the Command-line arguments section of the Supplementary Information.

## Data availability

All sequencing data and benchmark truth sets analyzed in this study are publicly available; dataset-specific sequencing and benchmark sources are listed in Supplementary Table 1 for the nanopore panel and Supplementary Table 16 for the PacBio HiFi panel. The PacBio HiFi datasets for HCC1937, HCC1954, H1437 and H2009 were obtained from CASTLE (NCBI BioProject PRJNA1086849), and those for HCC1395 and COLO829 from the public PacBio Revio 2023Q2 release (https://downloads.pacbcloud.com/public/revio/2023Q2/). The CytoScan HD microarray LOH profile of HCC1395 used for the orthogonal comparison was obtained from previously published data [37]. No new sequencing data were generated in this study.

## Code availability

The source code of LongPhase-TO is freely available at https://github.com/CCU-Bioinformatics-Lab/longphase-to. The exact command lines and parameters used to run LongPhase-TO and every tool it was benchmarked against are provided in the Command-line arguments section of the Supplementary Information, enabling all reported statistics to be reproduced.

## Competing interests

R.L. receives research funding from Oxford Nanopore Technologies. The other authors declare no competing interests.

## Author contributions

Y.-T.H. conceived and supervised the study and designed the algorithms. Z.-Y.C. implemented LongPhase-TO and performed the experiments and analyses. H.-F.F. implemented the regression models for somatic SNV and indel recalibration. Y.-J.Y. conducted the PacBio experiments. Z.Z. and R.L. contributed the somatic variant calling tools and datasets and advised on the benchmarking design. Z.-Y.C. and Y.-T.H. wrote the manuscript. All authors read and approved the final manuscript.

