## Supplementary material for "Somatic haplotype reconstruction and variant recalibration from tumor-only long-read sequencing"

**3 Supplementary Figures** **9**

**4 Supplementary Tables** **26**

**5 Command-line arguments** **45**

### 1 Supplementary Results

#### Robustness of somatic recalibration to the ClairS-TO model

To test whether the recalibration depends on the particular ClairS-TO model, we repeated the SNV benchmark with a ClairS-TO model trained on simulated data (ClairS-TO-ss), with and without LongPhase-TO, across the same eight datasets and DNA fraction levels (Supplementary Fig. 1).

The fraction-dependent profile observed for the default model was reproduced. Recalibration raised precision across 20 to 80% DNA fraction in nearly every dataset, at a modest cost to recall, and the F1 gain was largest at low DNA fraction and in the two datasets where the standalone precision of ClairS-TO-ss was lowest (HCC1937_UCSC and HCC1954_UCSC). At 100% DNA fraction, where ClairS-TO-ss was already most precise, recalibration lowered F1 in several datasets. The recalibration therefore acts as a conservative precision filter whose benefit concentrates where the standalone caller is least reliable, regardless of the ClairS-TO model used.

#### Cross-platform generalization of somatic recalibration to PacBio HiFi

Every model used by the triplet-graph recalibration was fitted once on nanopore data and then held fixed: the linear (indel) and third-order polynomial (SNV) predictors were trained on the HCC1395_HKU mixture at 0.6 tumor DNA fraction, and the site-level decision thresholds ($S_H > 0.8$; $L/(L+N) \geq 0.2$) were calibrated on the same sample. To test whether this haplotype-based recalibration depends on the sequencing platform, we applied the identical models and thresholds to six PacBio HiFi datasets (Supplementary Table 16), using the same *in silico* dilution design spanning 20–100% tumor DNA fraction. Five of these cell lines (HCC1395, HCC1937, HCC1954, H1437 and H2009) belong to the PacBio Revio tumor-normal cohort assembled for DeepSomatic, four of them obtained here from the CASTLE release and HCC1395 from the public PacBio Revio 2023Q2 release; COLO829 is an independent PacBio Revio benchmark from the same public release. The panel is therefore not the complete DeepSomatic cohort, which also includes Hs578T.

Every LongPhase-TO model was held off-platform. Both somatic callers were run with their PacBio HiFi models, so that the baselines against which recalibration is measured are platform-matched, whereas LongPhase-TO was run in its nanopore mode so that the nanopore-fitted triplet-graph coefficients were applied under exactly the settings in which they were trained. No coefficient was refitted and no threshold was retuned for PacBio, and the same applies to the tumor DNA fraction regression assessed in the following subsection (Supplementary Fig. 6). The experiment therefore measures transfer of the trained models alone, against callers that carry no comparable platform handicap.

Recalibration improved PacBio HiFi somatic calls to a degree comparable with nanopore. Averaged over the six datasets and five DNA-fraction levels, mean SNV F1 rose from 0.58 to 0.67 for ClairS-TO and from 0.60 to 0.69 for DeepSomatic-TO, improving 25 and 24 of 30 individual points, respectively (Supplementary Fig. 4). Mean indel F1 rose from 0.24 to 0.27 for ClairS-TO and from 0.28 to 0.32 for DeepSomatic-TO, improving 26 and 23 of 30 points (Supplementary Fig. 5). The gain was precision-driven, as on nanopore. Mean SNV precision rose by 0.19 for ClairS-TO and by 0.21 for DeepSomatic-TO, against mean recall changes of −0.02 and −0.13, and mean indel precision rose by 0.11 and 0.15 against mean recall changes of −0.01 and −0.05. The asymmetry between the two callers was systematic rather than incidental: DeepSomatic-TO recall decreased at all 30 points in both variant classes, whereas its indel precision increased at all 30. Per-mixture precision, recall and F1 for every caller and configuration are given in Supplementary Tables 11 and 12.

The dependence on tumor DNA fraction also reproduced the nanopore pattern. Mean SNV F1 gains were largest at low-to-intermediate fraction, reaching +0.11, +0.13 and +0.13 for ClairS-TO and +0.12, +0.14 and +0.15 for DeepSomatic-TO at fractions of 0.2, 0.4 and 0.6, and were smallest at 1.0. Restricted to mixtures below 1.0, mean SNV F1 increased by 0.12 for ClairS-TO and 0.13 for DeepSomatic-TO, with 23 and 22 of 24 points improved. The dataset-level ordering was preserved as well. HCC1954, the most difficult genome in the nanopore panel, again showed the largest gains: at 0.8 DNA fraction its SNV F1 rose from 0.36 to 0.66 for ClairS-TO and from 0.35 to 0.71 for DeepSomatic-TO, closely matching the corresponding nanopore values of 0.33 to 0.61 and 0.32 to 0.69.

One point departed from this pattern. In H2009 at 1.0 DNA fraction, recalibration of DeepSomatic-TO calls reduced SNV F1 from 0.88 to 0.31 and indel F1 from 0.39 to 0.18. This single point accounts for the only negative fraction-level averages in the analysis: at 1.0 DNA fraction the mean change in DeepSomatic-TO F1 was −0.07 for SNVs and −0.03 for indels with H2009 included, and +0.03 and 0.00 with it excluded. Pure tumor samples are the regime in which the fixed triplet-graph coefficients operate furthest from their training conditions, because the variant allele frequencies of heterozygous somatic and germline variants converge as tumor DNA fraction approaches 1.0 (Supplementary Fig. 20), and coefficients fitted on a nanopore sample at 0.6 carry no calibration for the corresponding PacBio HiFi error profile. Refitting the triplet-graph coefficients on PacBio data is the direct remedy and was not attempted here, because the purpose of this analysis was to measure transfer without retraining. Taken together, these results indicate that the haplotype evidence exploited by the triplet graph is a property of read-backed linkage rather than of a particular sequencing chemistry, while identifying near-pure samples as the boundary at which platform-specific recalibration becomes worthwhile.

##### Cross-platform generalization of tumor DNA fraction estimation to PacBio HiFi

The tumor DNA fraction regression was trained in the same single-fit manner as the recalibration models. The complete third-degree polynomial in the two haplotype-imbalance quartiles and the LOH burden was fitted on seven of the eight nanopore datasets, with HCC1954_UCSC held out as an independent test set (Supplementary Section 2.4), and its coefficients were then held fixed. To assess whether this estimator depends on the sequencing platform, the identical coefficients were applied without refitting to the six PacBio HiFi datasets, using the same dilution design and the same comparators as the nanopore panel: ASCAT in tumor-only (TO) and tumor-normal (T+N) modes, and PURPLE in tumor-only mode.

The quantity estimated is not the same across methods. ASCAT and PURPLE infer cellular purity jointly with ploidy from allele-specific copy-number profiles, whereas LongPhase-TO infers the fraction of sequenced molecules of tumor origin from germline haplotype imbalance alone, without copy-number segmentation, a global ploidy fit, or any purity and ploidy search (Supplementary Table 7). To place the methods on a common axis, the cellular purity $p$ reported by each comparator was converted to tumor DNA fraction with its own fitted ploidy $\kappa$ as $f = p\kappa/(p\kappa + 2(1 - p))$, assuming diploid normal cells (see Methods in the main text), and all ASCAT and PURPLE values reported here are these converted fractions. The unit of analysis is the cell-line dataset rather than the individual mixture, because the five DNA-fraction levels within a dataset are downsampled from a single library and are therefore not independent.

Accuracy on PacBio HiFi was comparable to the training platform. Mean absolute error was 0.06 for the DeepSomatic-TO configuration and 0.08 for ClairS-TO, with jackknife ranges of 0.03 to 0.06 and 0.04 to 0.09 when any single cell line was excluded, against 0.04 and 0.06 on nanopore (Supplementary Table 9). Lin's concordance correlation coefficient was 0.94 and 0.89. The estimator also remained close to a linear response over the dilution series, with slopes of 0.93 for both configurations and intercepts indistinguishable from zero, the closest approach to the identity relation among the five methods on either platform. Per-mixture estimates are given in Supplementary Table 13 and plotted in Supplementary Fig. 6.

Stratifying by dilution level shows that the methods fail in different regimes rather than by different amounts (Supplementary Table 10). LongPhase-TO with DeepSomatic-TO gave errors of 0.04, 0.03, 0.05, 0.08 and 0.09 from 0.2 to 1.0 DNA fraction, varying by a factor of about two across the range. PURPLE was more accurate than any other method at intermediate fractions, reaching 0.00 at 0.6 and 0.8, but its error rose to 0.16 in the pure samples, with individual estimates of 0.567 for HCC1954 and 0.619 for HCC1937 where the true fraction was 1.0. A purity and ploidy fit likely becomes weakly identified as tumor content approaches 100%, because a given allele-specific copy-number profile can be explained by high purity at one ploidy or lower purity at a multiple of it. ASCAT showed the opposite behavior, with errors of 0.41 (T+N) and 0.78 (TO) at 0.2 DNA fraction; ASCAT (T+N) then fell to 0.01 at every higher level, whereas ASCAT (TO) still reached 0.26 at 0.4 and 0.12 in the pure samples, and returned no estimate for the pure COLO829 sample.

ASCAT in tumor-normal mode reached 0.01 at four of the five levels but saturated toward 1.0 in the most dilute HCC1395 mixture, giving an overall mean absolute error of 0.09 against 0.06 for LongPhase-TO with DeepSomatic-TO. With six independent cell lines this difference is not statistically established. LongPhase-TO nonetheless matched or exceeded the accuracy of a matched-normal copy-number method across the whole dilution range, including the dilute mixtures in which both ASCAT modes fail, from haplotagging that the phasing stage had already computed, without a matched normal sample and without a copy-number or ploidy model.

The one dataset that departs from this agreement is the same one on both platforms. HCC1954 gave a mean absolute error of 0.18 for DeepSomatic-TO and 0.29 for ClairS-TO on PacBio, against 0.17 and 0.25 on nanopore, so the difficulty reproduces almost exactly across sequencing chemistries and is a property of the genome rather than of the platform. HCC1954 carries a whole-genome duplication and had the lowest LOH burden in the panel, which is the regime in which haplotype imbalance carries least information about tumor content. Consistent with this interpretation, PURPLE, which models copy number explicitly, achieved 0.10 on HCC1954 and was the more accurate method there. One caveat applies symmetrically to all methods on this dataset: the ground truth in these series is a read-level mixing fraction created by downsampling reads, and in a genome-duplicated sample the read-level fraction and the cellular fraction diverge, so HCC1954 scores each method against a quantity that at least one of them does not target. We therefore do not rank methods on this dataset.

### 2 Supplementary Methods

#### 2.1 Somatic variant recalibration and LOH detection

The following methods provide additional details on empirical threshold calibration, directional-clipping interval definition and regression-calibrated triplet-graph recalibration of somatic variant candidates.

##### 2.1.1 Empirical calibration of LOH detection thresholds

The two thresholds used for LOH classification were calibrated against the HCC1395 LOH and non-LOH reference intervals published by SEQC2. Variant allele frequency (VAF) distributions within these intervals showed a heterozygous peak near 0.5 in non-LOH regions that was depleted in LOH regions; variants with VAF $\geq 0.8$, or marked as homozygous in the input, were therefore counted as homozygous, and all remaining variants were counted as heterozygous (Supplementary Fig. 18a). After variants within short CNV/BFB-associated intervals were excluded, the regional heterozygosity ratio was calculated as $R_{\text{het}} = N_{\text{het}}/(N_{\text{het}} + N_{\text{hom}})$. The resulting distributions separated the SEQC2 reference LOH and non-LOH regions, supporting classification of a region as LOH when $R_{\text{het}} < 0.09$ (Supplementary Fig. 18b). Both thresholds were then held fixed and applied unchanged across all sequencing datasets.

##### 2.1.2 Directional clipping and genomic-event interval definition

At each ordered clip-bearing position $i$, LongPhase-TO counts soft- and hard-clipping operations longer than five bases at the left and right alignment ends as $F_i$ and $B_i$, respectively, and defines the signed directional signal as $d_i = F_i - B_i$. Positions with $F_i \geq 5$ or $B_i \geq 5$ are retained as direct clipping signals. To recover boundaries supported by dispersed lower-amplitude clipping, two successive directional rolling means, each spanning 100 clip-bearing positions, are applied to $d_i$ in each direction; the reverse profile is sign-inverted so that the two clipping orientations can be processed symmetrically. Local peaks with a transformed value of at least 0.25 are retained, and peaks separated by no more than 100 clip-bearing positions are represented by the larger peak. A forward candidate is discarded when a direct signal $d_j \geq 5$ occurs within the following 200 clip-bearing positions; the reciprocal criterion, $d_j \leq -5$ within the preceding 200 positions, is applied to reverse candidates. This prevents a strong localized breakpoint from being represented again as a distributed-signal candidate.

Support for each retained distributed-signal candidate is estimated from the cumulative directional signal, $C_i = \sum_{k=1}^{i} d_k$. After $C_i$ is smoothed over 100 clip-bearing positions to obtain $\widetilde{C}_i$, support at candidate position $p$ is defined as $A_p = \widetilde{C}_{p+10} - \widetilde{C}_{p-10}$. The sign of $A_p$ preserves clipping orientation, and its magnitude is assigned as the amplified clipping support (Supplementary Fig. 17).

Direct and amplified signals are scanned in genomic order. A signal with at least five supporting clips initiates a candidate event. An opposite-orientation signal within 10 kb completes an SGE interval when its support is at least $\lfloor c/4 \rfloor$, where $c$ is the maximum support observed for the initiating orientation. If no qualifying partner is present, the initiating signal is retained as an LGE boundary. The first and last clip-bearing positions are added as terminal LGE boundaries. SGE intervals are excluded from allele-state counting, whereas LGE boundaries partition the chromosome for LOH evaluation. These labels describe operational interval classes and do not assign a molecular CNV or BFB subtype. Supplementary Table 14 lists the fixed implementation parameters and their roles.

##### 2.1.3 Regression-calibrated triplet-graph recalibration

LongPhase-TO evaluates each candidate in the context of two neighboring variants. The reference and alternate alleles at the three loci define eight possible paths, whose weights are the numbers of spanning reads. Each candidate-containing triplet contributes at most one of three mutually exclusive evidence categories: $V_H$ denotes high-confidence three-path evidence comprising two parental germline paths and one somatic-descendant path; $V_L$ denotes lower-confidence, effectively two-path evidence that remains compatible with a single haplotype-specific somatic origin; and $V_N$ denotes a sufficiently supported topology accepted as neither $V_H$ nor $V_L$. The 12 predefined $V_H$ patterns represent four evolutionary configurations for each placement of the candidate at the left, middle or right node. Empirically, these patterns are enriched among validated true positive calls, whereas discordant $V_N$ topologies are more common among false positive calls (Supplementary Fig. 8).

***Regression-based scoring of high-confidence triplets***

For each triplet, let $r_1 \geq r_2 \geq r_3$ denote the read supports of the three highest-ranked paths. LongPhase-TO first evaluates the 12 predefined $V_H$ configurations. For a candidate $V_H$ configuration, the read counts of the two parental paths ($x$ and $y$) and the somatic-descendant path ($z$) are used as model features. The configuration is eligible only when its constituent paths are the three highest-ranked paths and both parental paths have at least two supporting reads. LongPhase-TO then applies a fixed, mutation-type-specific logistic model whose linear predictor $\eta$ is mapped to a probability $p = \{1+\exp(-\eta)\}^{-1}$; the triplet contributes a $V_H$ vote when $p \geq 0.5$. For indels, $\eta$ is linear in the path supports,

$$\eta_{\text{indel}} = -0.2337 + 0.7643x + 0.7979y - 0.7473z,$$

whereas for SNVs $\eta$ is a third-order polynomial in $x$, $y$ and $z$,

$$\begin{aligned}\eta_{\text{SNV}} = {} & 0.003754 + 0.014223x + 0.015017y + 0.011169z \\ & - 0.000769x^2 + 0.000674xy + 0.007199xz \\ & - 0.004500y^2 + 0.041002yz - 0.009079z^2 \\ & + 0.000015x^3 + 0.000010x^2y - 0.000177x^2z \\ & - 0.000033xy^2 - 0.000075xyz - 0.000055xz^2 \\ & + 0.000111y^3 - 0.000754y^2z - 0.000201yz^2 + 0.000092z^3.\end{aligned}$$

The coefficients were fitted on a single training sample, the HCC1395_HKU mixture at 0.6 tumor DNA fraction, and then held fixed and applied to all other datasets, which were used only for evaluation. Regression therefore calibrates the evidence contributed by each high-confidence triplet; it does not by itself make the final candidate-site call.

###### *Classification of low-confidence triplet patterns*

If no $V_H$ configuration is accepted for that triplet, LongPhase-TO considers $V_L$ evidence only when $r_3 < 0.5r_2$, indicating that the graph is effectively supported by two paths. The nine $V_L$ patterns comprise three candidate positions crossed with three permitted configurations for the weak third path: RRR and two other paths that do not carry the candidate alternate allele. A weak third path carrying the candidate alternate allele is incompatible with a single haplotype-specific somatic origin and therefore cannot contribute a $V_L$ vote. If neither $V_H$ nor $V_L$ is accepted, the triplet contributes a $V_N$ vote when $r_2 \geq 2$; triplets with less support do not vote. This subdivision prevents all nominal two-path configurations from being treated as equivalent.

###### *Candidate-level integration of triplet evidence*

The hierarchical classification described above is applied separately to every candidate-containing triplet. Different triplets for the same candidate may therefore contribute $V_H$, $V_L$ or $V_N$ votes, even though each individual triplet contributes at most one vote. Let $H$, $L$ and $N$ be the numbers of accepted $V_H$, accepted $V_L$ and non-somatic triplet votes for a candidate. For each accepted high-confidence triplet $g$, the somatic-path proportion $s_g$ is the support for the selected somatic-descendant path divided by the total support for paths carrying the candidate alternate allele, and $S_H = H^{-1}\sum_g s_g$. Let $\tau = 0.8$ and $\theta = 0.2$ denote the high- and low-confidence aggregation thresholds, respectively. The candidate is retained through either the high-confidence or low-confidence evidence route:

$$(H \geq 1 \text{ and } S_H > \tau) \quad \text{or} \quad \left(L > 0 \text{ and } \frac{L}{L+N} \geq \theta\right).$$

The OR combines alternative candidate-level evidence routes; it does not replace the hierarchical classification of each individual triplet. The distributions of $S_H$ and $L/(L+N)$ among truth-labelled calls support the selected values of $\tau = 0.8$ and $\theta = 0.2$, respectively (Supplementary Figs. 9 and 10). Thus, the upgraded procedure combines mutation-type-specific regression at the individual-graph level with deterministic evidence aggregation at the candidate-site level.

#### 2.2 Methylation-aware recalibration of somatic variants

Long-read platforms report 5-methylcytosine (5mC) status per read, allowing the CpG methylation neighborhood of each candidate to be examined at single-read resolution. When the input alignments carry 5mC tags, LongPhase-TO performs an additional recalibration of the somatic candidates retained by the triplet-graph model; when methylation calls are absent, the step is skipped and the triplet-graph output is returned unchanged. The rationale is developmental: a somatic variant arises post-zygotically within a tumor subpopulation that carries a characteristic CpG methylation state, whereas admixed normal cells retain the germline state. Consequently, reads carrying the somatic alternate allele tend to share a single methylation state (methylation-"pure"), whereas reads carrying the reference allele are a mixture of tumor- and normal-derived molecules and are therefore methylation-"mixed". Germline variants, which segregate in both compartments, yield methylation heterogeneity on both alleles, and sequencing or alignment artifacts produce no reproducible allelic asymmetry. This module is applied to both somatic SNV and indel candidates; the accompanying schematics illustrate the indel case.

For each candidate, reads are partitioned by the allele they carry at the candidate locus, reference-allele reads and alternate-allele reads, and the CpG sites falling within a symmetric window of $\pm d$ base pairs around the candidate position are analyzed. Ten features are computed and grouped into three complementary views of the local methylation pattern (Supplementary Fig. 11).

***Allelic methylation coverage.***

At each flanking CpG site, the number of methylated reads is counted separately for the reference and alternate alleles. Three features summarize these counts across the window: the mean reference-allele methylated-read count, the mean alternate-allele methylated-read count, and the mean absolute difference between the per-site reference and alternate counts. Because the somatic alternate allele is confined to the uniformly methylated tumor fraction, a somatic candidate exhibits a large allelic count difference, whereas germline variants and artifacts do not (Supplementary Fig. 12).

***Joint haplotype methylation proportions.***

At each flanking CpG site, the reads of each allele are labeled "pure" when they share a single methylation state (all methylated or all unmethylated) and "mixed" when both states are present. Pairing the reference- and alternate-allele labels assigns each site to one of four joint states: reference-pure/alternate-pure, reference-pure/alternate-mixed, reference-mixed/alternate-pure and reference-mixed/alternate-mixed. Three features record the fraction of covered CpG sites in the reference-mixed/alternate-mixed, reference-pure/alternate-mixed and reference-mixed/alternate-pure states; the fourth fraction is omitted as redundant, since the four proportions sum to one. The somatic signature of a methylation-pure alternate allele against a methylation-mixed reference allele is captured by an elevated reference-mixed/alternate-pure fraction (Supplementary Fig. 13).

***Methylation-cloud geometry.***

At each flanking CpG site, an allele methylation ratio is defined as the number of methylated reads divided by the total number of methylated and unmethylated reads, computed separately for the two alleles. Plotting the per-site alternate-allele ratio against the reference-allele ratio yields a cloud of points whose centroid, at coordinates (mean alternate ratio, mean reference ratio), locates the candidate in this methylation-ratio space. Four features are derived: the two centroid coordinates, the perpendicular distance of the centroid from the diagonal $y = x$, which measures the overall imbalance between reference- and alternate-allele methylation, and the mean absolute per-site difference between the reference and alternate ratios. Balanced, symmetric methylation places the centroid on the diagonal, whereas the allele-specific methylation expected of a somatic variant displaces it (Supplementary Fig. 14).

Together these ten features are supplied to a gradient-boosted decision-tree classifier (XGBoost), which assigns each candidate to the somatic, germline or artifact class; only candidates classified as somatic are retained. Consistent with the training strategy used elsewhere in LongPhase-TO, the classifier was trained on a single sample, the HCC1395_NYGC mixture at 0.6 tumor DNA fraction, and then held fixed and applied unchanged to all other cell-line datasets and DNA-fraction levels, which were used only for evaluation.

***Classifier and training-set selection.***

The choice of a gradient-boosted tree model was made empirically (Supplementary Fig. 15). Seven candidate classifiers (XGBoost, histogram-based gradient boosting, random forests, extremely randomized trees, logistic regression, a linear support-vector machine and linear discriminant analysis) were compared by held-out ROC-AUC on the binary somatic-versus-non-somatic indel task under two cross-validation schemes, each applied both across cell lines and across tumor DNA fractions: leave-one-out, in which the model was trained on all groups but one and evaluated on the held-out group, and leave-four-out, in which it was trained on a single group and evaluated on the remaining four. The four tree-ensemble methods consistently outperformed the three linear methods, and XGBoost attained the highest or tied-highest held-out ROC-AUC in every scheme (median approximately 0.93 for leave-one-cell-line-out and approximately 0.96 for leave-one-fraction-out), motivating its selection. Under the more stringent leave-four-out setting, in which a single group supplied the training data, the held-out ROC-AUC of XGBoost remained at approximately 0.90 or above, confirming that the single-sample training strategy generalizes robustly across both cell lines and tumor DNA fractions.

#### 2.3 LOH-aware joint phasing of germline and somatic variants

LongPhase-TO constructs a local allele-linkage graph from the recalibrated variants and the detected LOH segments (Supplementary Fig. 19). Reference and alternate alleles are represented as separate nodes. For each read, an allele observation is connected to observations at up to the next 35 informative non-homozygous variants; traversal is interrupted across a gap of more than 300 kb between consecutive informative variants.

At an LOH transition, the graph additionally retains a limited set of connections between heterozygous variants outside the segment and informative homozygous variants within it. Each ordinary read observation contributes an edge weight of 1. An edge involving an allele observation with base quality below 12 is assigned a weight of 0.1, and observations from alignments marked as unreliable contribute a weight of 0.01.

Variants are visited in genomic order. For two non-somatic variants, parallel support is the sum of the reference–reference and alternate–alternate edge weights, whereas crossed support is the sum of the reference–alternate and alternate–reference weights. When either site is somatic, only linkages involving the somatic alternate allele are used to determine its parental haplotype. A pair is not propagated when the ratio of the smaller to the larger of the parallel and crossed supports exceeds 0.7, because the two configurations are insufficiently distinguishable. Accepted pairwise comparisons cast weighted votes for the HP1 or HP2 assignment of downstream variants; strongly concordant comparisons are upweighted, and potentially unreliable indel comparisons are downweighted. If the accumulated HP1 and HP2 support is equal, the current variant starts a new phase block rather than being connected without directional evidence.

The provisional phase is refined through the reads. For each read, allele observations consistent with HP1 and HP2 are counted, with unreliable observations downweighted or omitted. A read is assigned to a haplotype when the larger haplotype count accounts for more than 0.65 of its informative support and the read contributes more than one informative observation. Haplotype-specific allele counts from these assigned reads are then used to retain or revise a variant phase when one configuration accounts for more than 0.75 of the support. At a somatic site, this comparison uses the alternate-allele support in the two read haplotypes and records the supported somatic allele as a descendant of HP1 or HP2.

At each LOH boundary, LongPhase-TO compares read links between variants on opposite sides of the heterozygous-to-homozygous transition. A connection is accepted when one allele configuration accounts for at least 0.8 of the cross-boundary support. Votes from the retained connections identify whether HP1 or HP2 continues through the LOH segment. The other parental haplotype is represented as absent within the segment, whereas somatic alleles remain assigned to descendants of the retained haplotype. Connections at the distal boundary restore the corresponding parental label and phase-set continuity in the following diploid region.

#### 2.4 Tumor DNA fraction regression from haplotype imbalance and LOH

The tumor DNA fraction is predicted from three genome-wide features. The first and third quartiles of the haplotype-imbalance distribution, $q_1$ and $q_3$, capture its lower and upper portions and jointly describe its location and spread. The LOH burden $r$, defined as the total detected LOH length divided by the total analyzed chromosome length, provides complementary information on the genome-wide extent of parental haplotype loss. A single complete third-degree (cubic) polynomial in these features, comprising all 20 monomials of total degree at most three, was fitted on seven of the eight cell-line datasets, with HCC1954_UCSC held out as an independent test set, and then held fixed; the same coefficients are applied whether somatic variants are supplied by ClairS-TO or DeepSomatic-TO. The fitted predictor is

$$\begin{aligned} f = & -40.7226 + 194.2179\,q_1 - 13.4902\,q_3 - 8.6631\,r \\ & - 169.4964\,q_1^2 - 161.1149\,q_1q_3 + 71.1042\,q_1r + 74.7865\,q_3^2 - 38.3745\,q_3r + 22.1523\,r^2 \\ & + 6.5747\,q_1^3 + 145.5591\,q_1^2q_3 + 25.5427\,q_1^2r - 4.7007\,q_1q_3^2 - 127.0712\,q_1q_3r + 19.2526\,q_1r^2 \\ & - 30.2605\,q_3^3 + 74.4106\,q_3^2r - 33.5680\,q_3r^2 - 4.5039\,r^3, \end{aligned}$$

and the reported tumor DNA fraction is $f$ clamped to the interval $[0, 1]$. These coefficients were also applied unchanged to the PacBio HiFi panel, without refitting or platform-specific tuning, so that the regression was evaluated off its training platform in the same manner as the recalibration models (Supplementary Fig. 6).

### 3 Supplementary Figures

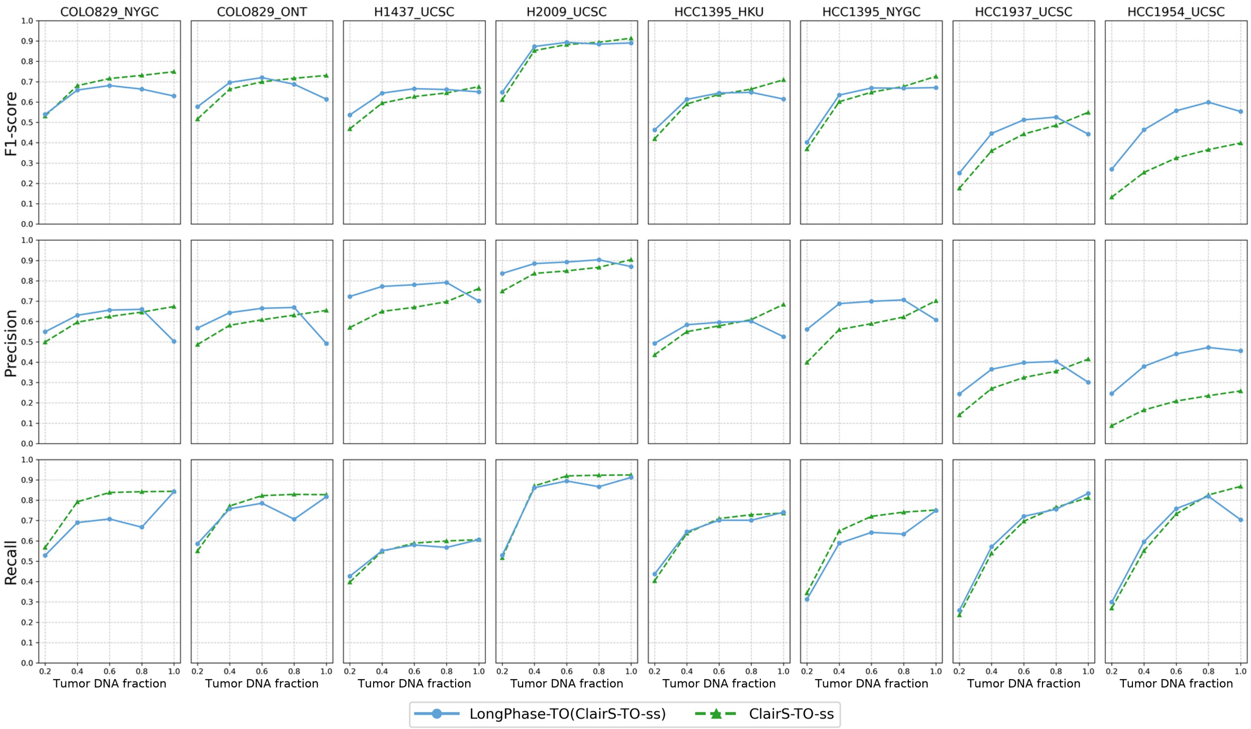

**Supplementary Fig. 1**: Somatic variant calling performance of ClairS-TO-ss with and without LongPhase-TO across eight cancer cell line datasets at tumor DNA fractions from 20% to 100%. LongPhase-TO generally improved precision, while recall was reduced to varying degrees.

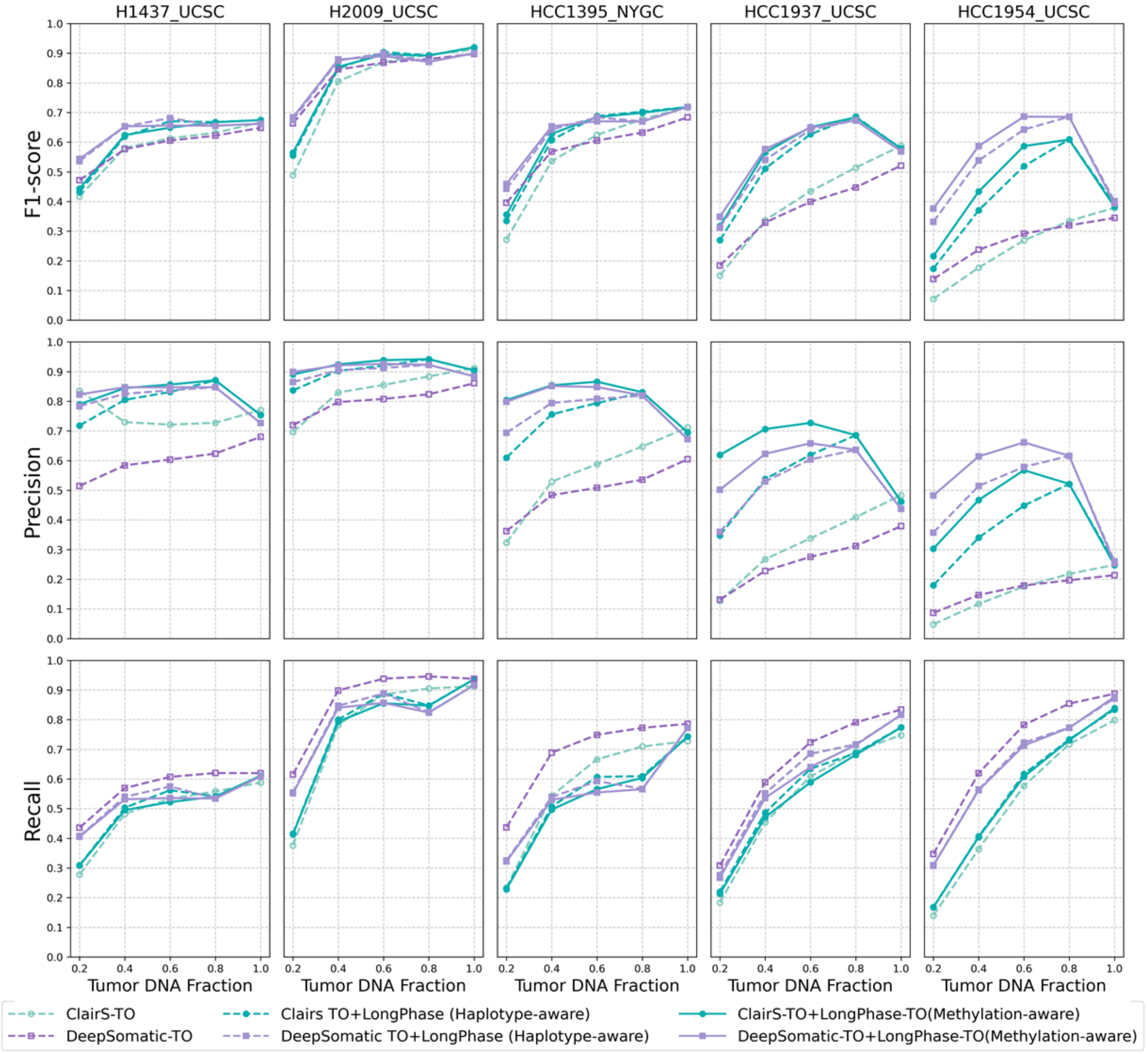

**Supplementary Fig. 2**: Methylation-aware recalibration improves somatic SNV calling beyond haplotype-aware recalibration alone. F1-score, precision and recall of ClairS-TO and DeepSomatic-TO SNV calling are shown for the five tumor cell line datasets with 5-methylcytosine (5mC) tags available (H1437_UCSC, H2009_UCSC, HCC1395_NYGC, HCC1937_UCSC and HCC1954_UCSC) across DNA fractions from 20% to 100%, comparing each caller alone, the caller after haplotype-aware (triplet-graph) recalibration, and the caller after the additional methylation-aware recalibration step. Averaged over the five datasets and both callers, the methylation-aware step raised mean SNV precision by 0.05 (0.68 to 0.72) at a mean recall cost of 0.01, giving a mean F1 gain of 0.01. Precision improved in 34 of the 50 caller-dataset-fraction combinations and declined in 2, with the largest gains in HCC1937_UCSC (0.08) and HCC1954_UCSC (0.07), the two datasets with the lowest haplotype-aware accuracy.

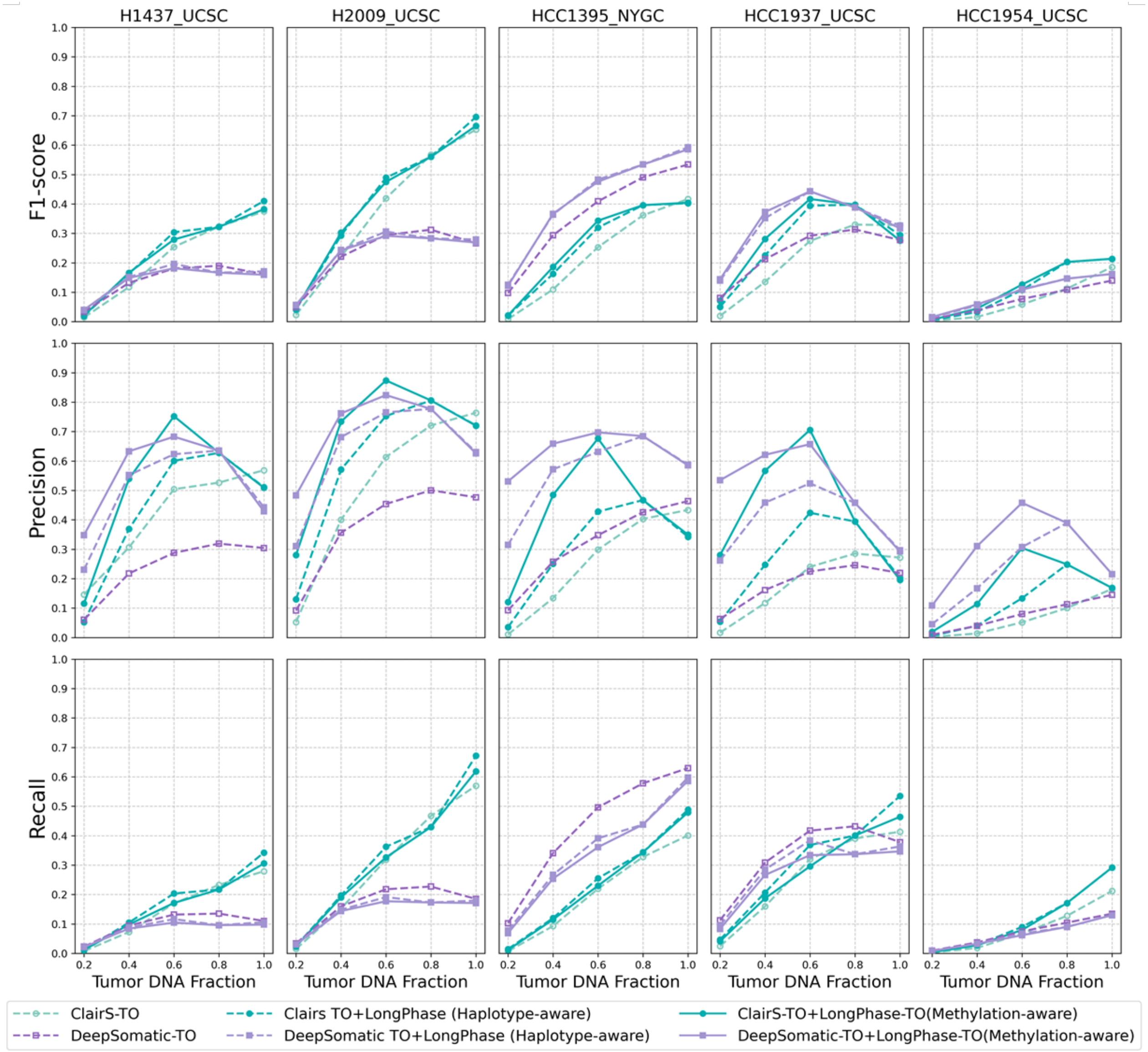

**Supplementary Fig. 3**: Methylation-aware recalibration improves the precision of somatic indel calling beyond haplotype-aware recalibration alone. F1-score, precision and recall of ClairS-TO and DeepSomatic-TO indel calling are shown for the five tumor cell line datasets with 5-methylcytosine (5mC) tags available (H1437_UCSC, H2009_UCSC, HCC1395_NYGC, HCC1937_UCSC and HCC1954_UCSC) across DNA fractions from 20% to 100%, comparing each caller alone, the caller after haplotype-aware (triplet-graph) recalibration, and the caller after the additional methylation-aware recalibration step. Averaged over the five datasets and both callers, the methylation-aware step raised mean indel precision by 0.09 (0.40 to 0.49) at a mean recall cost of 0.01, leaving mean F1 essentially unchanged. Precision improved in 31 of the 50 caller-dataset-fraction combinations and declined in 7, with the largest gains in HCC1937_UCSC (0.14) and HCC1395_NYGC (0.09).

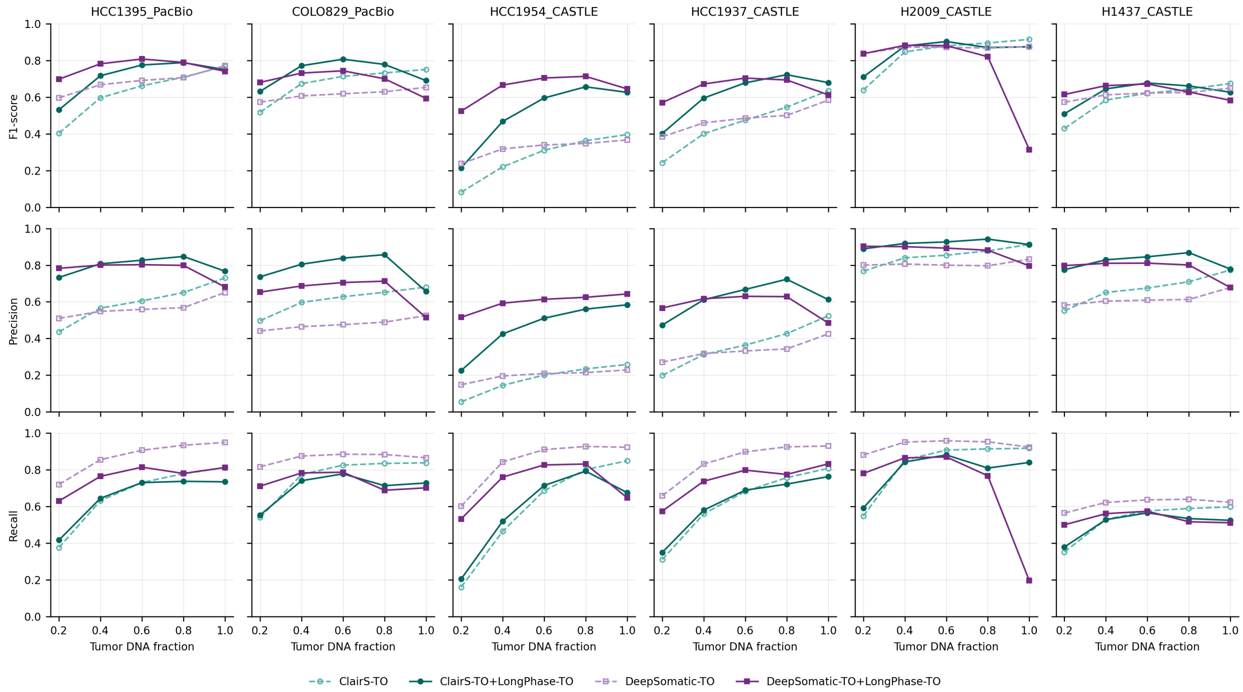

**Supplementary Fig. 4**: Cross-platform generalization of haplotype-aware somatic SNV recalibration to PacBio HiFi. F1-score, precision and recall of ClairS-TO and DeepSomatic-TO SNV calling, with and without LongPhase-TO recalibration, are shown for six PacBio HiFi cancer cell line datasets across tumor DNA fractions from 20% to 100%. Both callers were run with their PacBio HiFi models, whereas all triplet-graph regression coefficients and site-level thresholds were those fitted on nanopore data (HCC1395_HKU at 0.6 tumor DNA fraction) and were applied without refitting or platform-specific tuning. Recalibration raised mean SNV F1 from 0.58 to 0.67 for ClairS-TO and from 0.60 to 0.69 for DeepSomatic-TO, improving 25 and 24 of 30 points, with the largest gains at 20–60% DNA fraction as observed on nanopore data. The exception is H2009 at 100% DNA fraction with DeepSomatic-TO, where F1 decreased from 0.88 to 0.31.

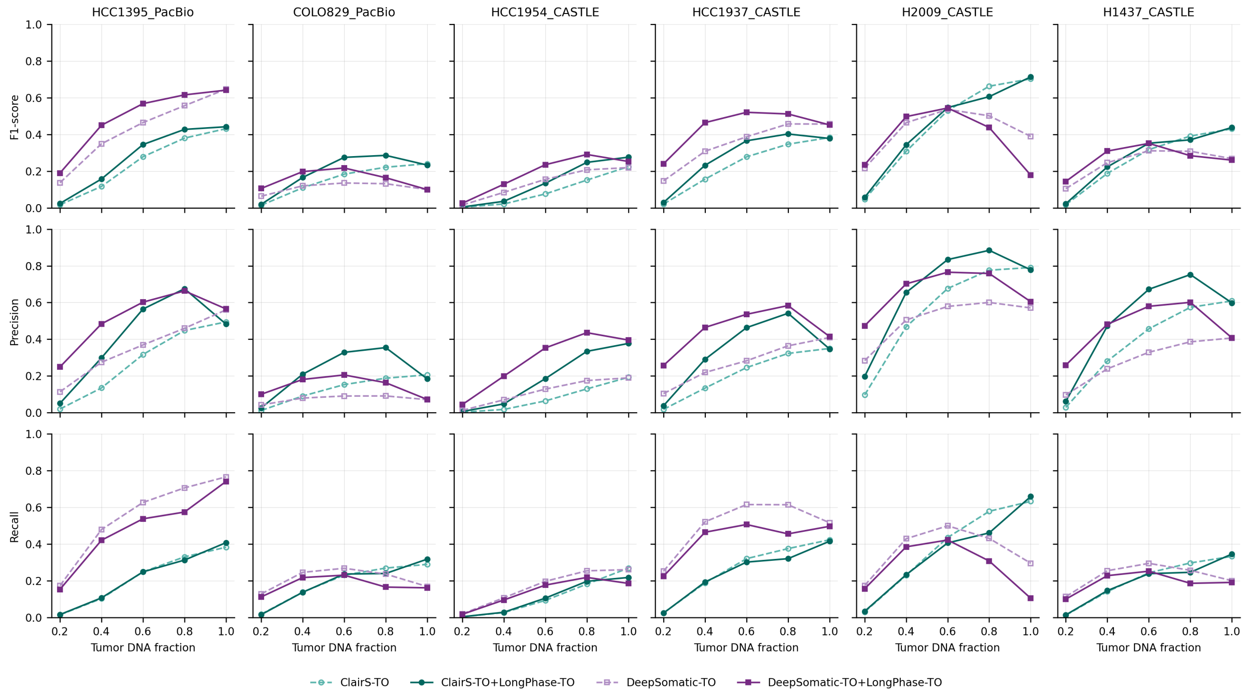

**Supplementary Fig. 5**: Cross-platform generalization of haplotype-aware somatic indel recalibration to PacBio HiFi. F1-score, precision and recall of ClairS-TO and DeepSomatic-TO indel calling, with and without LongPhase-TO recalibration, are shown for the same six PacBio HiFi cancer cell line datasets and tumor DNA fractions as Supplementary Fig. 4, again using the nanopore-fitted models without refitting. Recalibration raised mean indel F1 from 0.24 to 0.27 for ClairS-TO and from 0.28 to 0.32 for DeepSomatic-TO, improving 26 and 23 of 30 points. As for SNVs, the gains concentrated at low-to-intermediate DNA fraction, and the largest decrease occurred in H2009 at 100% DNA fraction with DeepSomatic-TO (0.39 to 0.18).

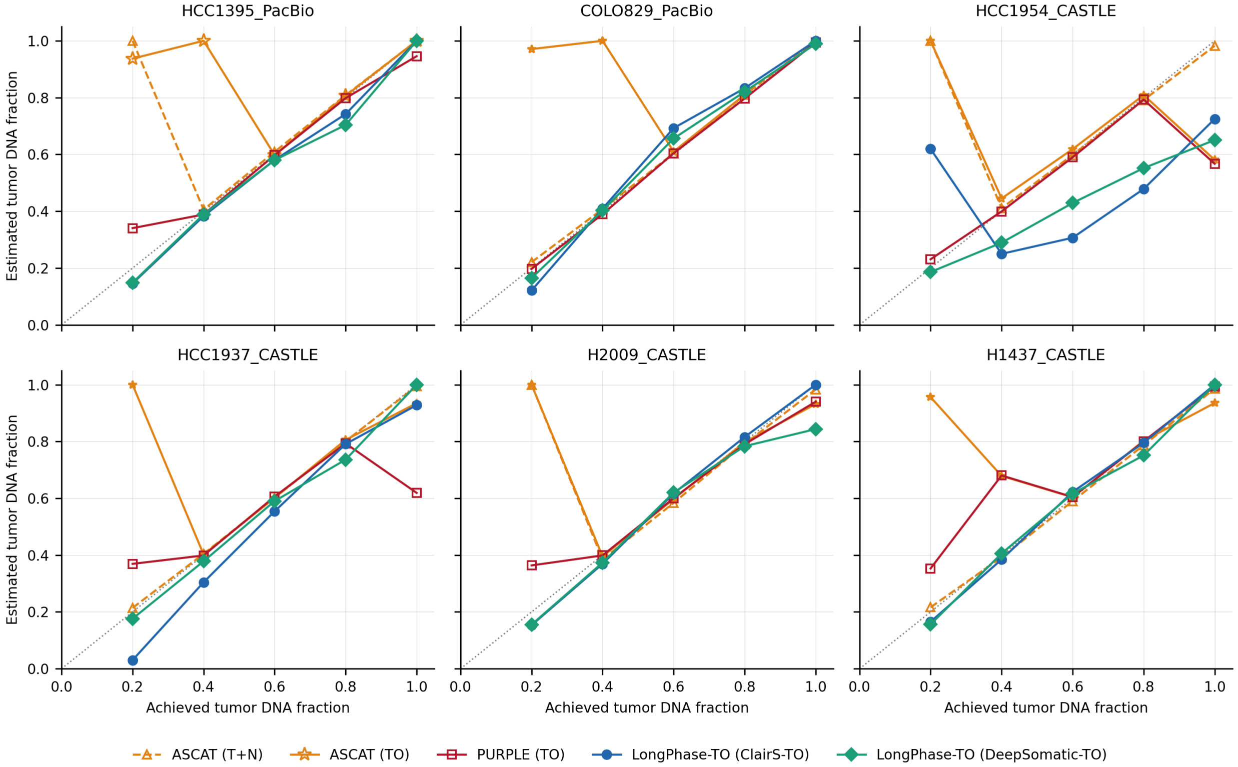

**Supplementary Fig. 6**: Tumor DNA fraction estimation on PacBio HiFi with nanopore-fitted coefficients. Estimated tumor DNA fraction is plotted against the achieved read-level mixing fraction for six PacBio HiFi cell line datasets across five dilution levels, comparing ASCAT in tumor-normal (T+N) and tumor-only (TO) modes, PURPLE in tumor-only mode, and LongPhase-TO using somatic variants from ClairS-TO or DeepSomatic-TO; the diagonal indicates the ideal prediction ($y = x$). The LongPhase-TO regression coefficients were those fitted on the nanopore panel and were applied without refitting or platform-specific tuning. Mean absolute error was 0.06 for the DeepSomatic-TO configuration and 0.08 for ClairS-TO, with slopes of 0.93 and intercepts indistinguishable from zero. The methods degrade in different regimes: ASCAT is least accurate in the most dilute mixtures, PURPLE in the near-pure samples, and LongPhase-TO is comparatively uniform across the range. HCC1954, which carries a whole-genome duplication and the lowest LOH burden in the panel, is the least accurate dataset for LongPhase-TO on both platforms. Per-mixture values are given in Supplementary Table 13.

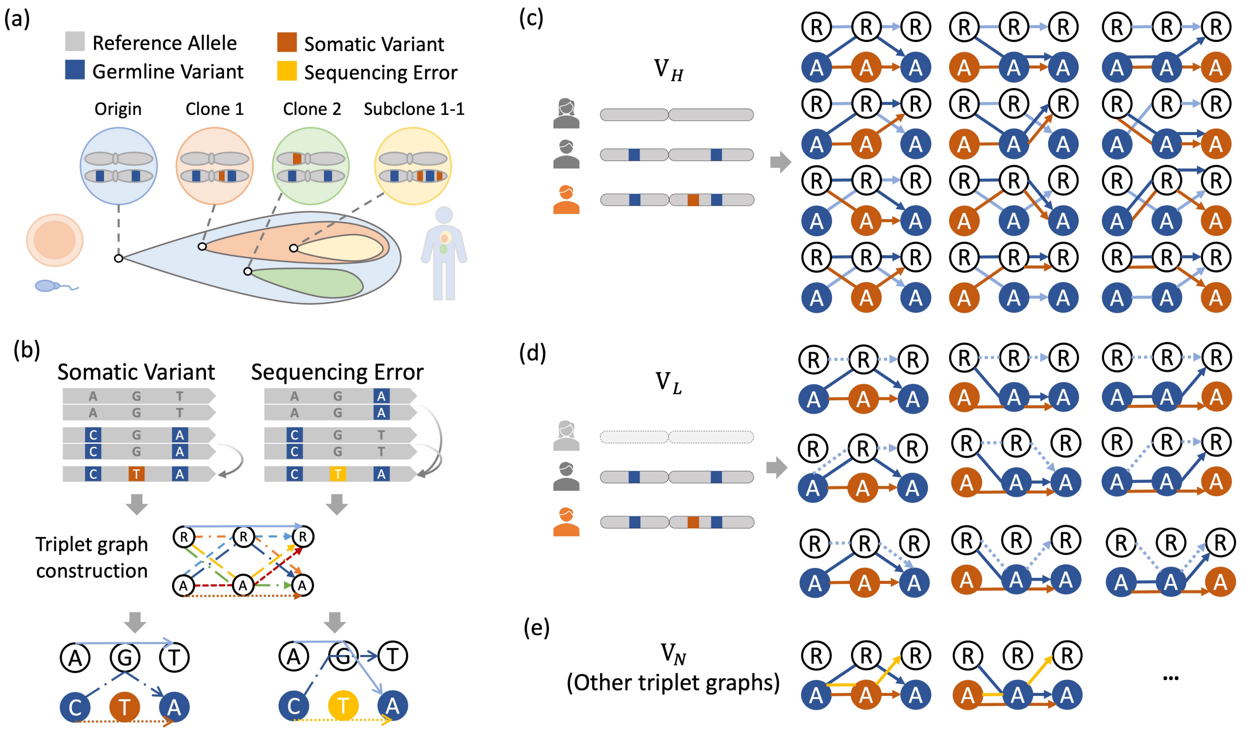

**Supplementary Fig. 7**: Triplet-graph construction and pattern classification. (a) A somatic allele arises on one parental haplotype and is inherited through clonal evolution. (b) Reads spanning three adjacent variant positions define a triplet graph containing up to eight reference/alternate haplotype paths; the examples contrast the consistent haplotype linkage of a somatic variant with the discordant linkage of a sequencing error. (c) The 12 high-confidence ($V_H$) configurations comprise four evolutionary configurations for each candidate position (middle, left and right columns), each containing two parental germline paths and one somatic-descendant path. (d) The nine lower-confidence ($V_L$) configurations comprise three candidate positions crossed with three permitted weak third paths: RRR and two paths that bypass the candidate alternate allele. (e) Sufficiently supported topologies classified as neither $V_H$ nor $V_L$ contribute a $V_N$ vote; triplets with insufficient support do not vote.

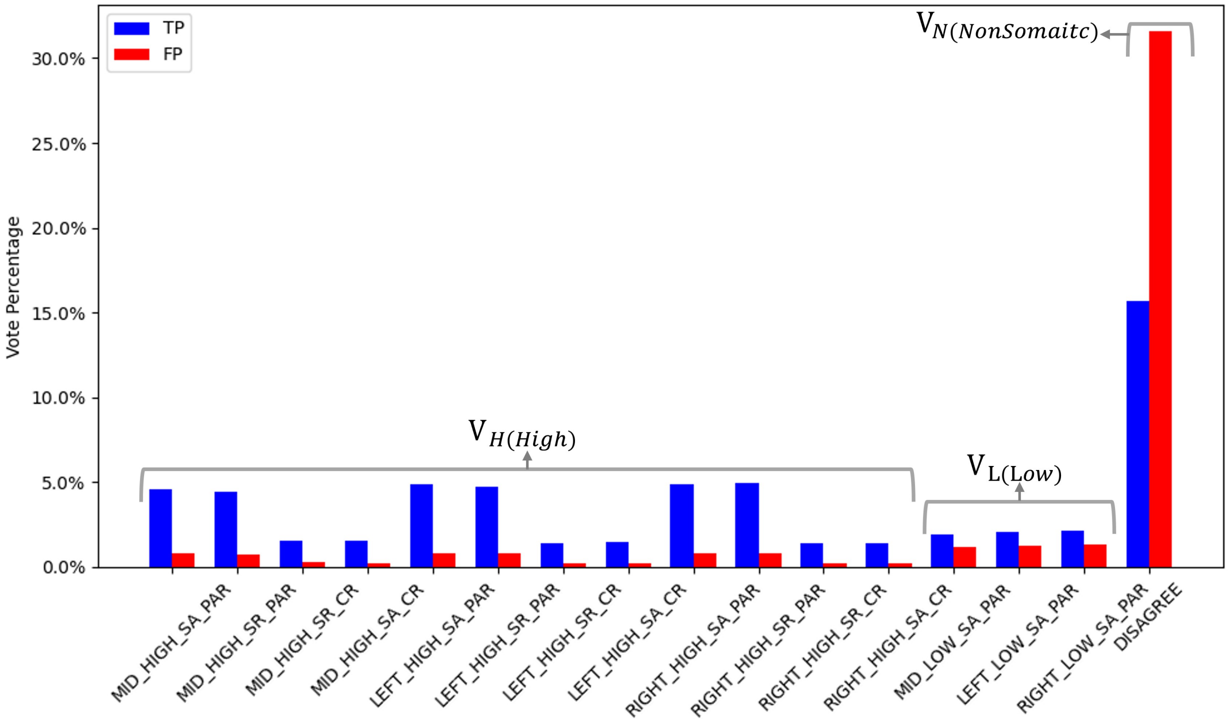

**Supplementary Fig. 8**: Comparison of True Positive (TP, blue) and False Positive (FP, red) variant calls across predefined evidence patterns. The patterns are grouped into three categories: high-confidence ($V_H$), low-confidence ($V_L$), and non-somatic ($V_N$). $V_H$ patterns are strongly enriched for TPs with minimal FP contribution, $V_L$ patterns occur less frequently but remain reliable indicators of true variants, and $V_N$ patterns are dominated by the DISAGREE category, which contributes substantially to both TPs and FPs and is therefore ambiguous and unreliable. These distributions provide the rationale for adopting a filtering strategy that prioritizes variants supported by $V_H$ and $V_L$ patterns.

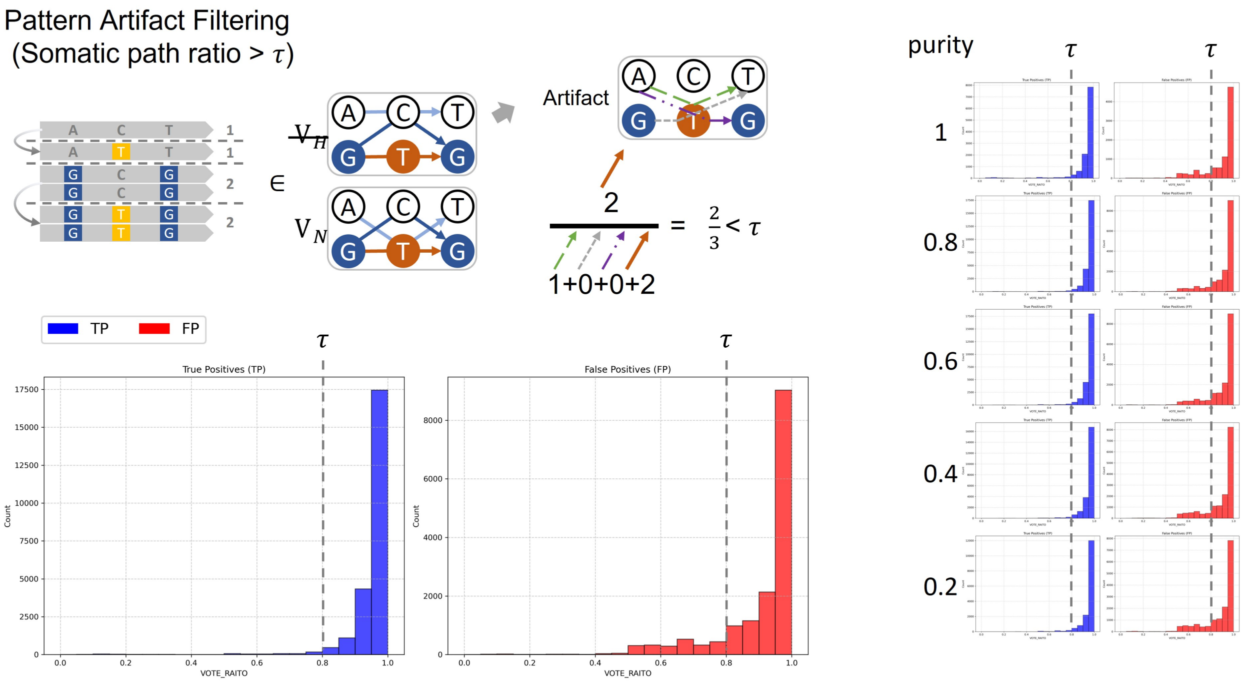

**Supplementary Fig. 9**: Distribution of the mean high-confidence somatic-path proportion, $S_H$, used during site-level evidence aggregation. True positive candidates (TP, blue) are enriched near 1.0, whereas false positive candidates (FP, red) have lower and more dispersed values. The separation across tumor DNA fractions from 1.0 to 0.2 supports the retained threshold $S_H > \tau$, where $\tau = 0.8$, for candidates with at least one regression-accepted $V_H$ triplet.

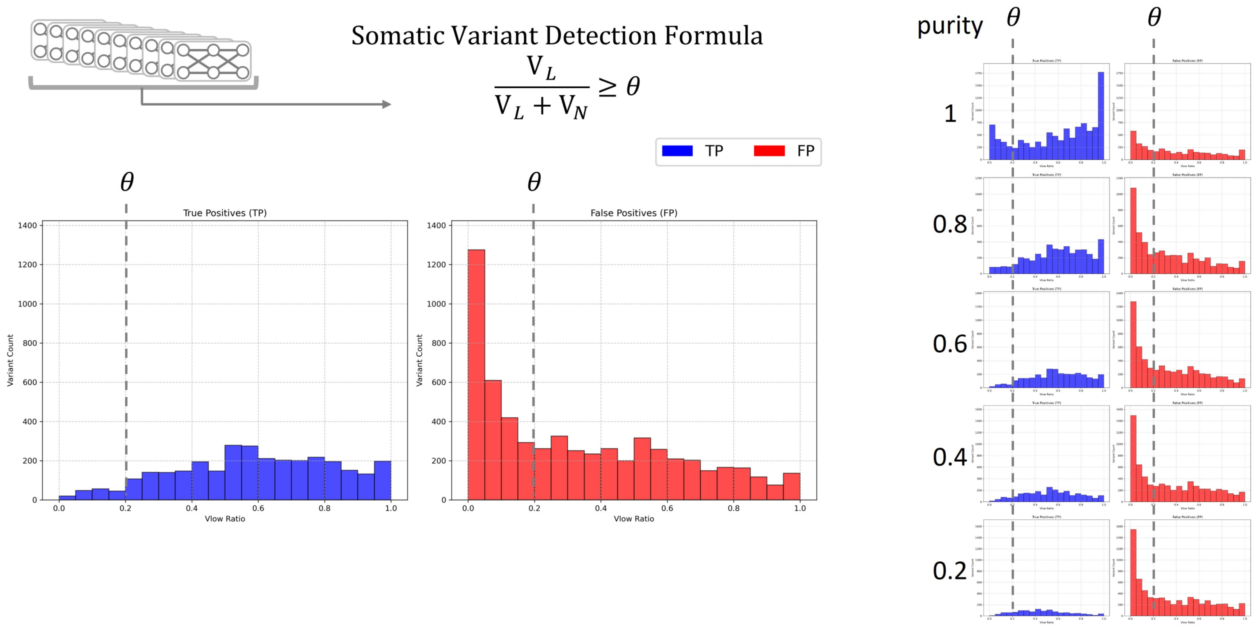

**Supplementary Fig. 10**: Distribution of the site-level low-confidence evidence fraction, $L/(L+N)$, for validated true positive (TP, blue) and false positive (FP, red) candidates. False positive candidates are concentrated near zero, whereas true positive candidates extend toward higher values. The distributions across tumor DNA fractions from 1.0 to 0.2 support the retained threshold $L/(L+N) \geq \theta$, where $\theta = 0.2$, when no high-confidence route is satisfied.

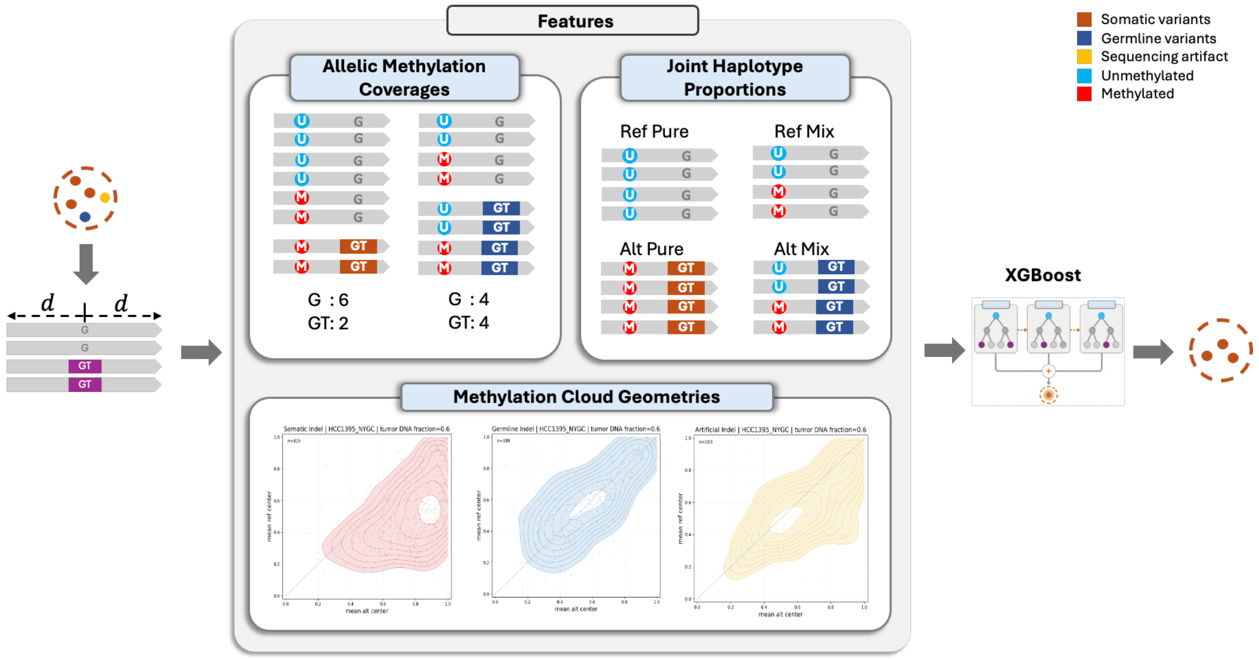

**Supplementary Fig. 11**: Overview of methylation-aware recalibration of somatic variant candidates. When the input alignments carry 5-methylcytosine (5mC) tags, reads spanning each candidate are partitioned by the allele they carry at the candidate locus (reference, "G"; alternate, e.g. "GT"), and the CpG sites within a symmetric window of $\pm d$ base pairs are analyzed at single-read resolution (methylated, red "M"; unmethylated, cyan "U"). Ten features are computed across three complementary views: allelic methylation coverage (per-site methylated-read counts on each allele), joint haplotype methylation proportions (per-site classification of each allele as methylation-pure or methylation-mixed), and methylation-cloud geometry (per-site allele methylation ratios and the geometry of their centroid; kernel-density clouds are shown for somatic, germline and artifact indels in HCC1395_NYGC at 0.6 tumor DNA fraction). A gradient-boosted decision-tree classifier (XGBoost) uses these features to label each candidate as a somatic variant, germline variant or sequencing artifact, retaining only candidates classified as somatic.

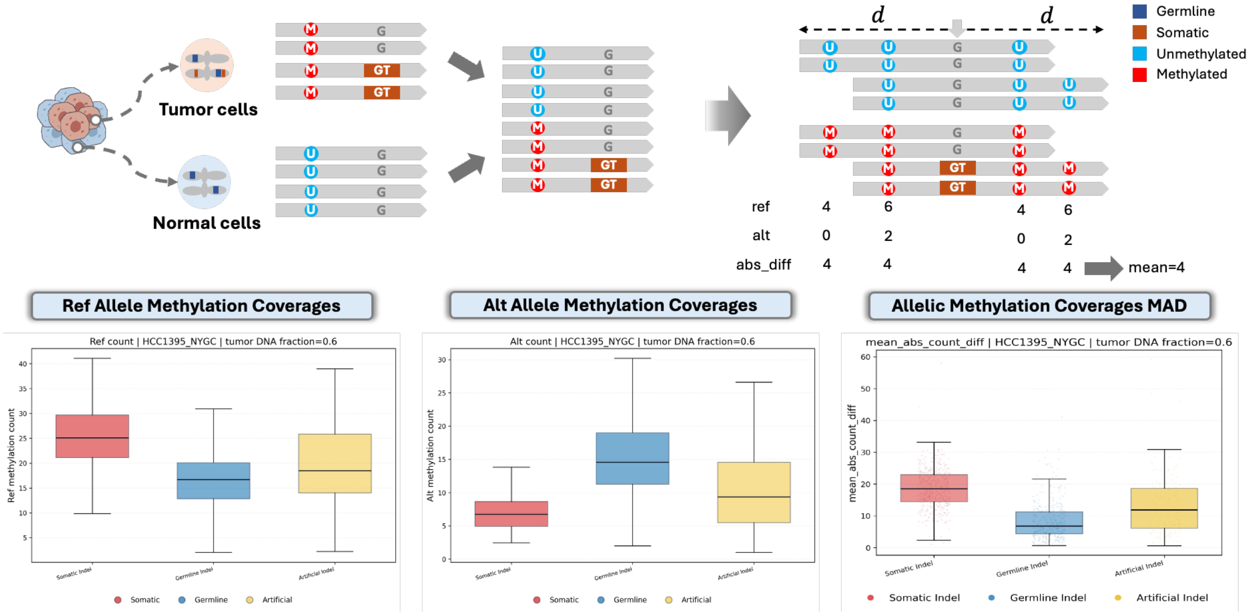

**Supplementary Fig. 12**: Allelic methylation coverage features. Reads spanning a candidate are split by the allele they carry (reference, “G”; alternate, e.g. “GT”), and at each CpG site within the flanking window the number of methylated reads is counted separately for the two alleles (methylated, red “M”; unmethylated, cyan “U”). The per-site reference and alternate methylated-read counts and their absolute difference are averaged over the window to give the mean reference count, the mean alternate count and the mean absolute count difference (MAD). Because the somatic alternate allele is confined to the uniformly methylated tumor fraction, whereas the reference allele mixes tumor- and normal-derived reads, somatic indels exhibit the largest allelic count difference; box plots (HCC1395_NYGC at 0.6 tumor DNA fraction) contrast somatic, germline and artifact indels for each feature.

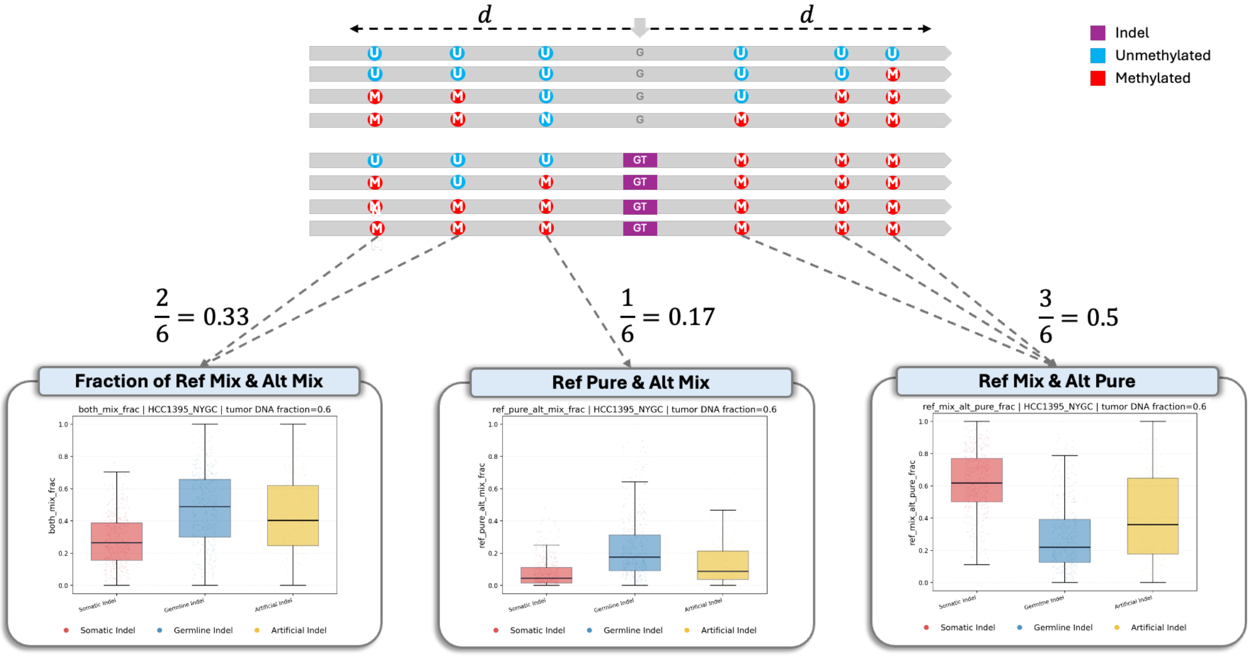

**Supplementary Fig. 13**: Joint haplotype methylation proportion features. At each CpG site within the flanking window, the reads of each allele are labeled "pure" when they share a single methylation state (all methylated, red "M"; or all unmethylated, cyan "U") and "mixed" when both states are present. Pairing the reference- and alternate-allele labels assigns each site to one of four joint states, and the fraction of covered CpG sites in each state is recorded; three of the four proportions are retained as features (the fourth is redundant, as they sum to one). In the illustrated window of six covered CpG sites, the reference-mixed/alternate-mixed, reference-pure/alternate-mixed and reference-mixed/alternate-pure fractions are $2/6 = 0.33$, $1/6 = 0.17$ and $3/6 = 0.5$, respectively. The somatic signature of a methylation-pure alternate allele against a methylation-mixed reference allele is captured by an elevated reference-mixed/alternate-pure fraction; box plots (HCC1395_NYGC at 0.6 tumor DNA fraction) contrast somatic, germline and artifact indels for each feature.

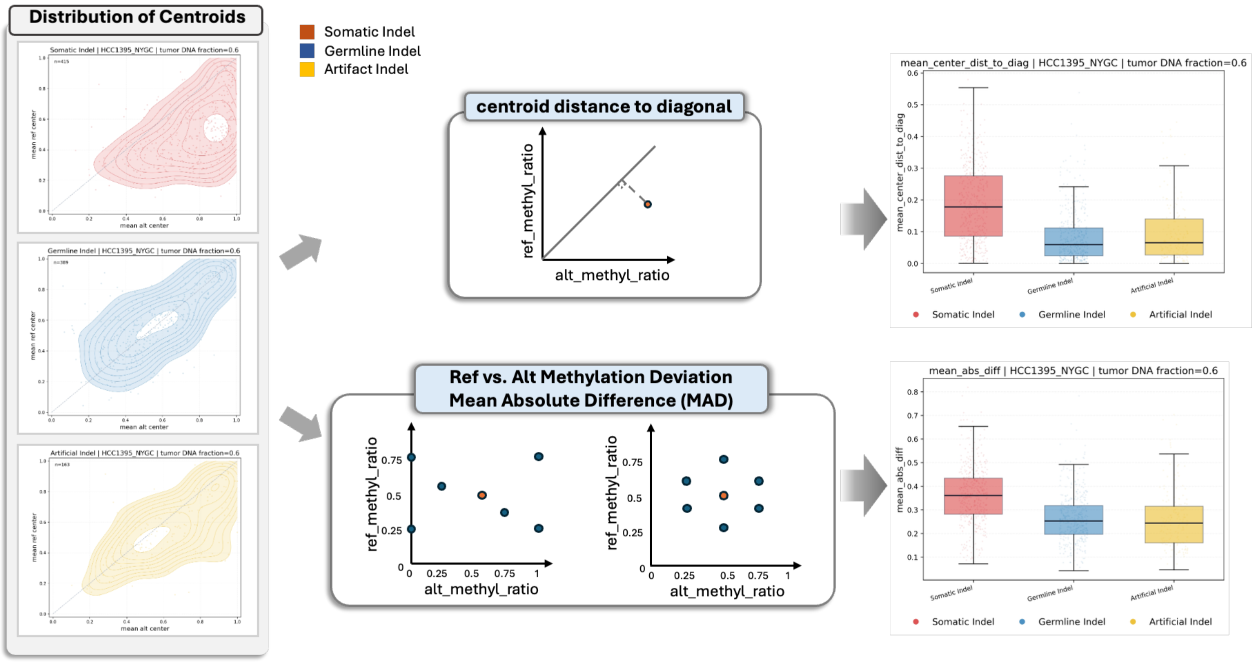

**Supplementary Fig. 14**: Methylation-cloud geometry features. At each CpG site within the flanking window, an allele methylation ratio is computed for the reference and alternate alleles, and the per-site (alternate ratio, reference ratio) points form a cloud whose centroid (mean alternate ratio, mean reference ratio) is retained as two features; the kernel-density distributions of these centroids differ between somatic, germline and artifact indels (left). Two further features summarize the cloud geometry: the perpendicular distance of the centroid from the diagonal $y = x$, which quantifies the overall imbalance between reference- and alternate-allele methylation, and the mean absolute per-site difference between the reference and alternate ratios (MAD). Balanced methylation places the centroid near the diagonal, whereas the allele-specific methylation of a somatic variant displaces it; box plots (HCC1395_NYGC at 0.6 tumor DNA fraction) show that somatic indels have the largest centroid-to-diagonal distance and ratio MAD.

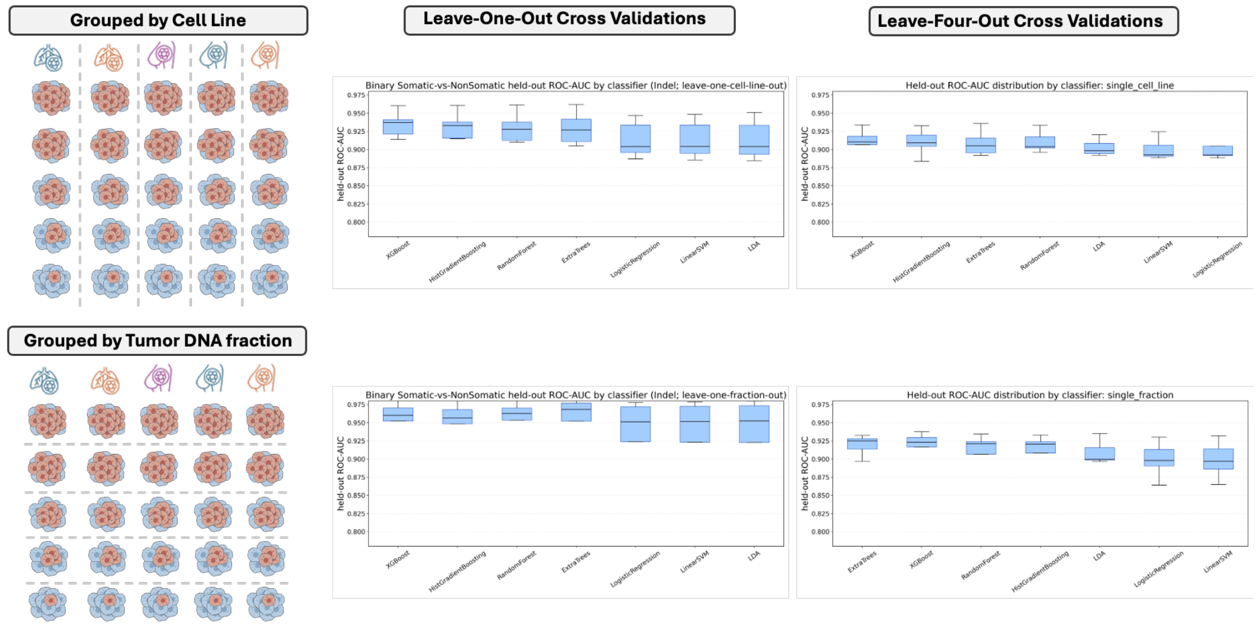

**Supplementary Fig. 15**: Selection of the classifier for methylation-aware recalibration. Seven candidate classifiers were compared by held-out ROC-AUC on the binary somatic-versus-non-somatic indel task under two cross-validation schemes, each applied both across cell lines and across tumor DNA fractions. Left, leave-one-out cross-validation, in which the model is trained on all groups but one and evaluated on the held-out group; right, leave-four-out cross-validation, in which it is trained on a single group and evaluated on the remaining four. The four tree-ensemble methods (XGBoost, histogram-based gradient boosting, random forests and extremely randomized trees) outperform the three linear methods (logistic regression, a linear support-vector machine and linear discriminant analysis), and XGBoost is consistently the top or tied-top performer. Its held-out ROC-AUC remains high even under the more stringent leave-four-out setting, supporting both the choice of XGBoost and the single-sample training strategy. The box plots convey the quantitative results; the cell-line and tumor-fraction icons are schematic.

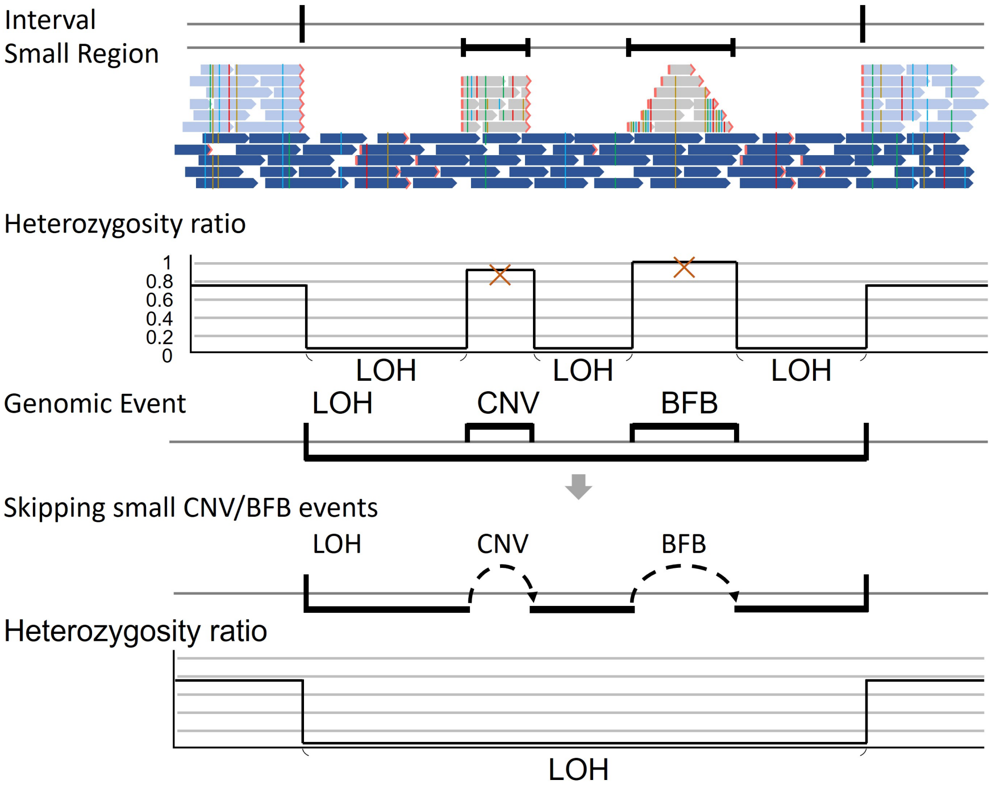

**Supplementary Fig. 16**: Chromosome-scale LOH detection after exclusion of short CNV/BFB-associated intervals. Candidate LOH boundaries define the region evaluated. Small CNV or BFB events embedded within a broader LOH tract can locally increase heterozygosity and split an initial LOH call. Omitting variants within these short intervals before calculating the regional heterozygosity ratio allows the flanking heterozygosity-depleted segments to be classified as one continuous arm-level LOH event.

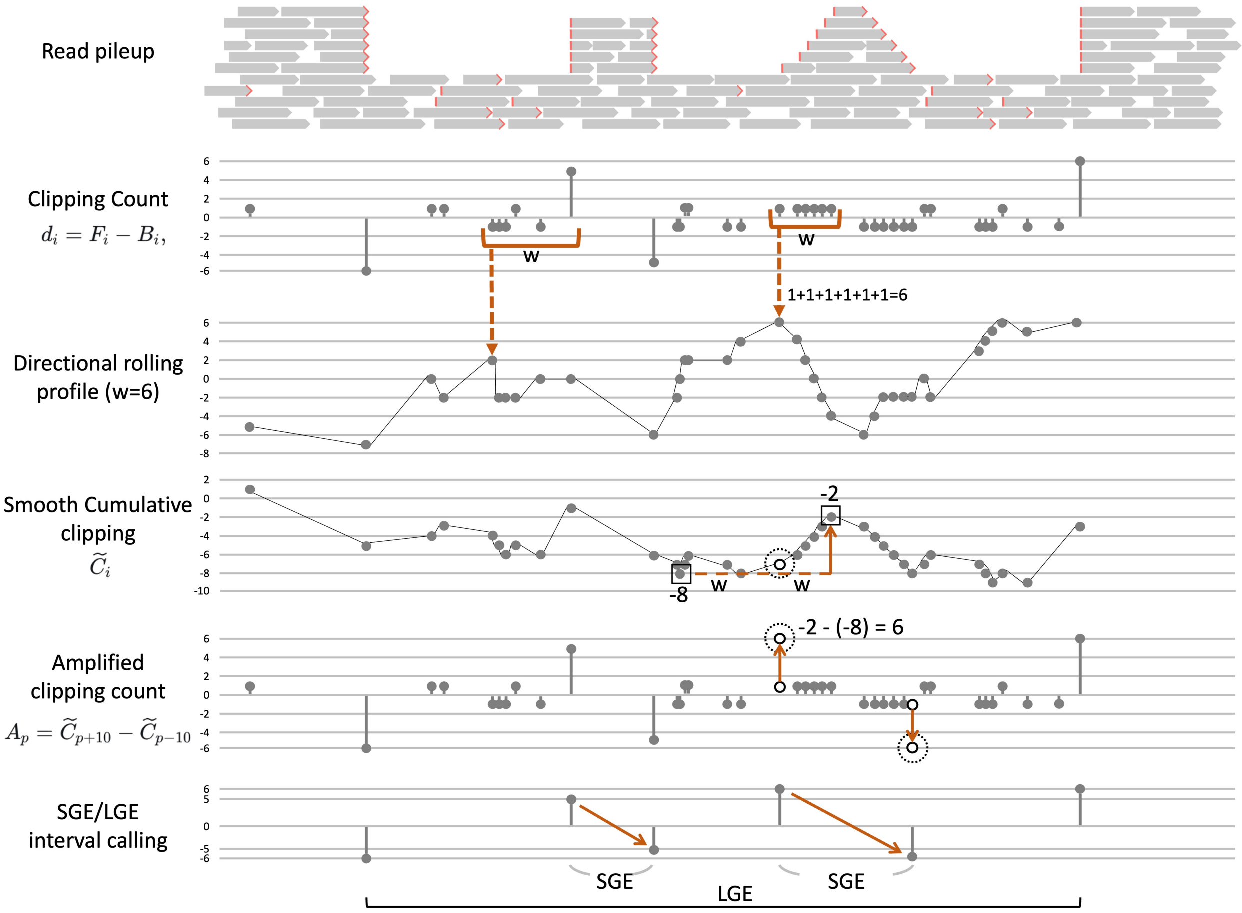

**Supplementary Fig. 17**: Directional clipping and genomic-event interval definition. Soft- and hard-clipping operations longer than five bases are counted at the left and right alignment ends as $F_i$ and $B_i$, respectively, and their difference defines the signed directional signal, $d_i = F_i - B_i$. Strong localized signals are retained without smoothing, whereas directional rolling profiles consolidate dispersed lower-amplitude support. For each resulting candidate at position $p$, support is estimated from the change in the smoothed cumulative clipping signal across the candidate, $A_p = \widetilde{C}_{p+10} - \widetilde{C}_{p-10}$; its sign preserves clipping orientation. Nearby signals of opposite orientation are paired to delimit short intervals associated with local CNV or BFB events (SGE), whereas unpaired signals are retained as candidate boundaries of broader LOH regions (LGE). The schematic uses a six-position rolling window for clarity; full definitions and fixed implementation parameters are provided in Supplementary Methods (Section 2.1.2) and Supplementary Table 14. SGE and LGE are operational labels and do not assign a molecular event subtype.

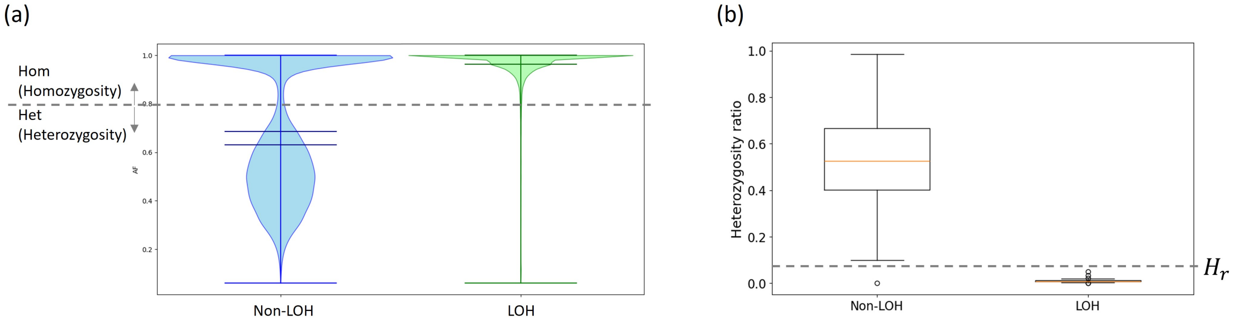

**Supplementary Fig. 18**: Empirical calibration of LOH classification thresholds using the HCC1395 LOH and non-LOH reference intervals published by SEQC2. (a) Variant allele frequency (VAF) distributions for variants in non-LOH (blue) and LOH (green) regions. The depletion of the heterozygous VAF peak in LOH regions supports classifying variants with VAF $\geq 0.8$ as homozygous. (b) Distributions of the regional heterozygosity ratio, $R_{\text{het}} = N_{\text{het}}/(N_{\text{het}} + N_{\text{hom}})$, after exclusion of short CNV/BFB-associated intervals. The separation between non-LOH and LOH regions supports classifying regions with $R_{\text{het}} < 0.09$ as LOH. Both thresholds were fixed and applied unchanged across all sequencing datasets.

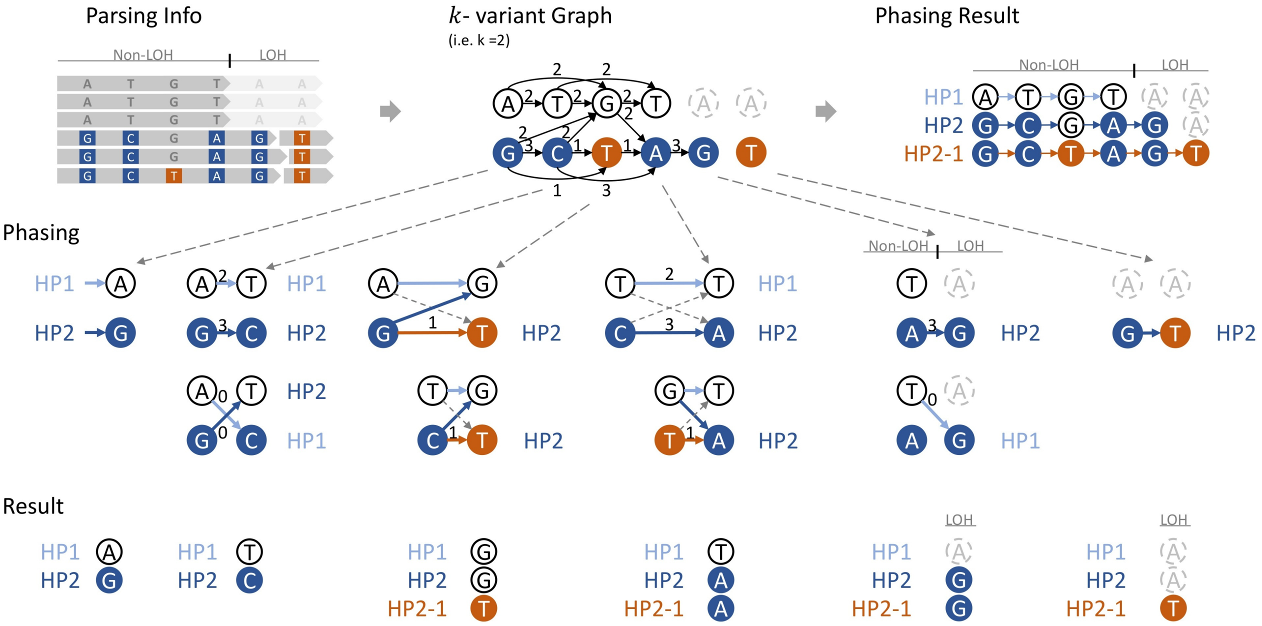

**Supplementary Fig. 19**: Joint germline and somatic phasing across LOH. Read-level allele observations are converted into a local allele-linkage graph in which nodes represent reference or alternate alleles and weighted edges represent their co-occurrence on reads. Parallel and crossed linkage evidence is combined across neighboring variants to assign alleles to the parental haplotypes HP1 and HP2. A somatic alternate allele is assigned to its supporting parental haplotype and represented as a derived somatic haplotype, such as HP2-1. At an LOH boundary, read connections between flanking heterozygous variants and homozygous variants within the LOH segment identify the retained parental haplotype; the lost path is shown in grey, whereas the retained parental and somatic paths continue through the LOH region. The schematic uses $k = 2$ for clarity; implementation details are provided in Supplementary Methods (Section 2.3).

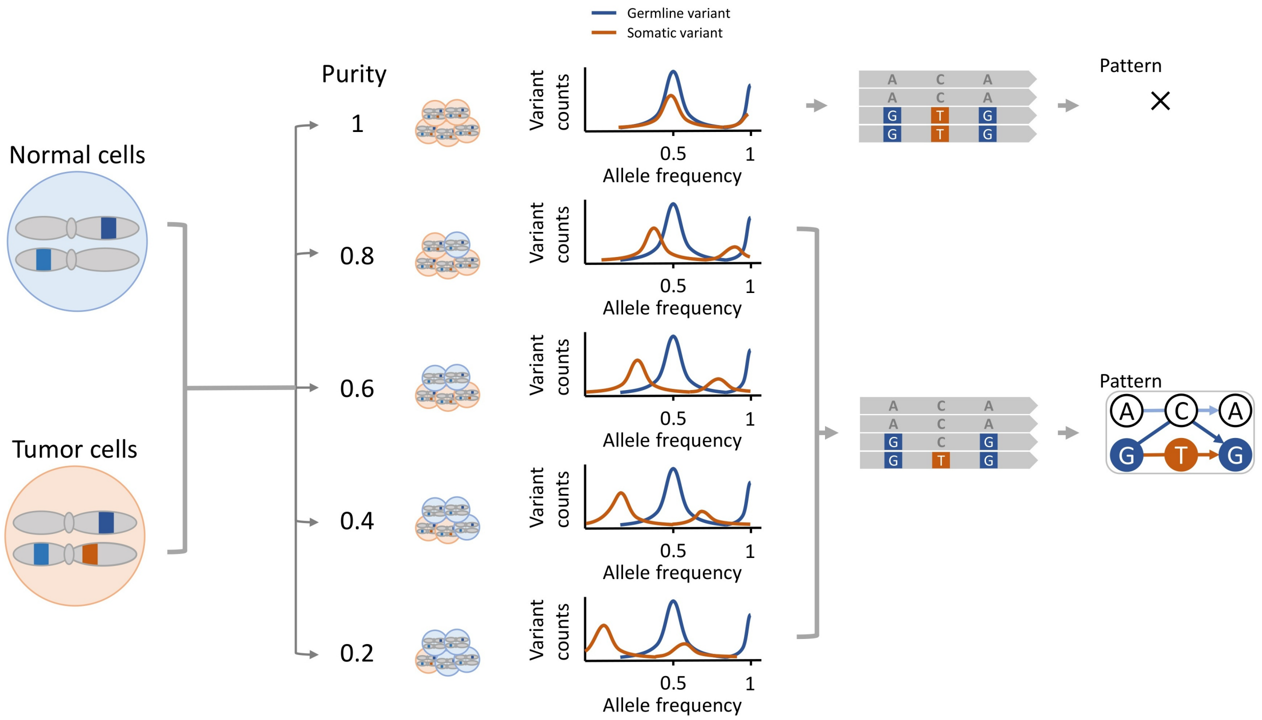

**Supplementary Fig. 20**: Effect of tumor DNA fraction on variant allele frequencies (VAFs). The VAF of a heterozygous germline variant remains stable at approximately 0.5 across DNA fraction levels, whereas the VAF of a heterozygous somatic variant decreases proportionally with decreasing DNA fraction. At high DNA fraction, the distributions of somatic and germline variants overlap, complicating statistical separation.

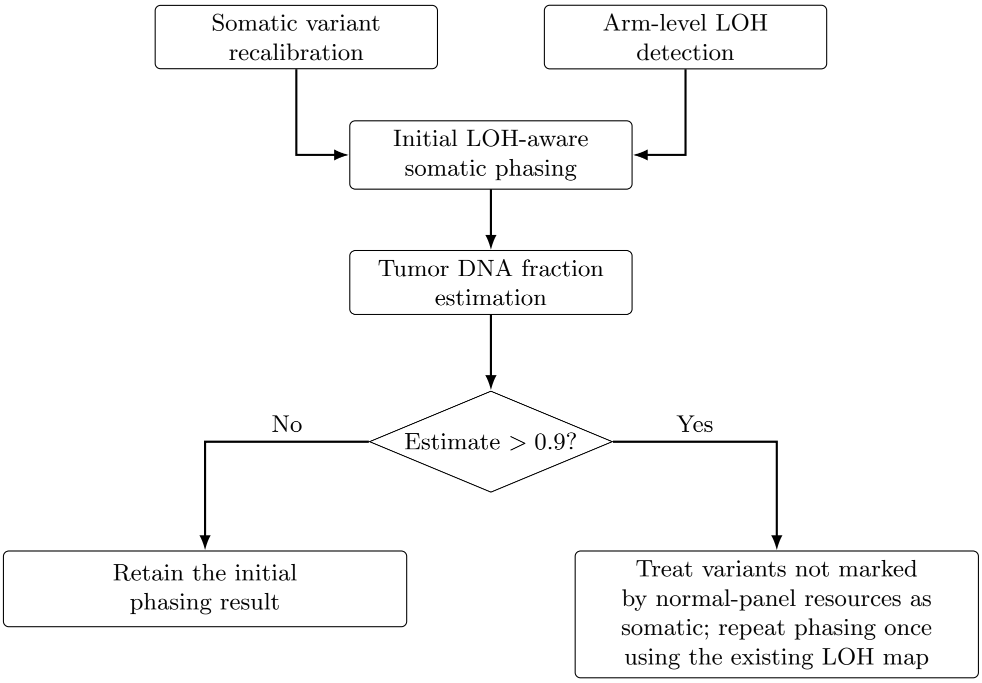

**Supplementary Fig. 21**: Conditional second phasing pass for high-DNA-fraction samples. Somatic variant recalibration and arm-level LOH detection are completed before the initial LOH-aware phasing pass. When the resulting tumor DNA fraction estimate exceeds 0.9, variants not marked by the normal-panel resources are treated as somatic and phasing is repeated once using the existing LOH map; otherwise, the initial phasing result is retained. The tumor DNA fraction is not re-estimated.

### 4 Supplementary Tables

**Supplementary Table 1**: Nanopore cell line datasets and corresponding benchmark truth sets, with normal and tumor coverages. All eight datasets were generated by Oxford Nanopore sequencing. The Source column names the group that produced the reads (HKU, NYGC, ONT or UCSC) and not the sequencing platform, and the Benchmark column names the origin of the somatic truth set (SEQC2, NYGC or DeepSomatic). In particular, COLO829_NYGC denotes nanopore reads produced by NYGC, evaluated against the NYGC somatic truth set.

| Dataset | Cancer Type | Source | Benchmark | Normal | Tumor |
|---|---|---|---|---|---|
| HCC1395_HKU | Breast ductal carcinoma | HKU | SEQC2 | 45.70x | 76.27x |
| HCC1395_NYGC | Breast ductal carcinoma | NYGC | SEQC2 | 29.14x | 79.52x |
| COLO829_ONT | Malignant melanoma | ONT | NYGC | 39.83x | 33.45x |
| COLO829_NYGC | Malignant melanoma | NYGC | NYGC | 45.85x | 108.49x |
| HCC1954_UCSC | Breast ductal carcinoma | UCSC | DeepSomatic | 34.03x | 82.44x |
| HCC1937_UCSC | Breast ductal carcinoma | UCSC | DeepSomatic | 25.68x | 158.44x |
| H2009_UCSC | Non-small-cell lung cancer | UCSC | DeepSomatic | 30.07x | 106.92x |
| H1437_UCSC | Non-small-cell lung cancer | UCSC | DeepSomatic | 49.28x | 79.10x |

COLO829 and HCC1395 each contribute two independent nanopore datasets to this panel, produced by different groups, and COLO829 and the other five cell lines also appear in the separate PacBio HiFi panel of Supplementary Table 16. Datasets sharing a cell line are independent sequencing runs and were analyzed separately throughout; no reads were pooled across them.

**Supplementary Table 2**: Comparison of conventional phasing tools with LongPhase-TO.

| Sample | Phasing tool | Phased ratio | Blocks | Block N50 (bp) | Block sum (bp) |
|---|---|---|---|---|---|
| COLO829_ONT | HapCUT2 | 0.531 | 3569 | 4645832 | 2523260676 |
| COLO829_ONT | WhatsHap | 0.530 | 3686 | 4635320 | 2518385299 |
| COLO829_ONT | LongPhase | 0.498 | 3477 | 4887010 | 2540883450 |
| COLO829_ONT | LongPhase-TO | 0.594 | 1110 | 9852371 | 2835034563 |
| COLO829_NYGC | HapCUT2 | 0.557 | 15016 | 584415 | 2187808909 |
| COLO829_NYGC | WhatsHap | 0.556 | 15261 | 575997 | 2182781406 |
| COLO829_NYGC | LongPhase | 0.521 | 15017 | 570504 | 2176223422 |
| COLO829_NYGC | LongPhase-TO | 0.607 | 6576 | 1403687 | 2664715137 |
| H1437_UCSC | HapCUT2 | 0.540 | 2376 | 7048644 | 2625509787 |
| H1437_UCSC | WhatsHap | 0.539 | 2398 | 7048644 | 2624731180 |
| H1437_UCSC | LongPhase | 0.502 | 2287 | 7729758 | 2654323753 |
| H1437_UCSC | LongPhase-TO | 0.698 | 529 | 19323318 | 2864794192 |
| H2009_UCSC | HapCUT2 | 0.557 | 2252 | 5825292 | 2677986524 |
| H2009_UCSC | WhatsHap | 0.556 | 2292 | 5723430 | 2676922405 |
| H2009_UCSC | LongPhase | 0.517 | 2204 | 7217802 | 2706782369 |
| H2009_UCSC | LongPhase-TO | 0.682 | 638 | 23169348 | 2863280796 |
| HCC1395_HKU | HapCUT2 | 0.602 | 4553 | 1678506 | 2536199864 |
| HCC1395_HKU | WhatsHap | 0.600 | 4662 | 1667777 | 2532455755 |
| HCC1395_HKU | LongPhase | 0.561 | 4581 | 1754642 | 2569265774 |
| HCC1395_HKU | LongPhase-TO | 0.576 | 893 | 16768123 | 2847723783 |
| HCC1395_NYGC | HapCUT2 | 0.499 | 21138 | 275106 | 1972073127 |
| HCC1395_NYGC | WhatsHap | 0.498 | 21510 | 270769 | 1966798050 |
| HCC1395_NYGC | LongPhase | 0.468 | 21057 | 279702 | 1980748528 |
| HCC1395_NYGC | LongPhase-TO | 0.693 | 5227 | 13441849 | 2726465382 |
| HCC1937_UCSC | HapCUT2 | 0.402 | 4537 | 1447908 | 2430726898 |
| HCC1937_UCSC | WhatsHap | 0.401 | 4650 | 1420186 | 2425582425 |
| HCC1937_UCSC | LongPhase | 0.375 | 4517 | 1513047 | 2454370721 |
| HCC1937_UCSC | LongPhase-TO | 0.713 | 1135 | 25207324 | 2830053891 |
| HCC1954_UCSC | HapCUT2 | 0.652 | 1969 | 5927639 | 2711566888 |
| HCC1954_UCSC | WhatsHap | 0.650 | 1999 | 5908073 | 2710095182 |
| HCC1954_UCSC | LongPhase | 0.610 | 1923 | 5912510 | 2736740481 |
| HCC1954_UCSC | LongPhase-TO | 0.659 | 1115 | 6477286 | 2828110080 |

**Supplementary Table 3**: Loss of heterozygosity (LOH) summary metrics across samples and methods.

| Sample | LOH Methods | LOH Burden (%) | No. Blocks | LOH Sum (bp) | LOH N50 Size (bp) |
|---|---|---|---|---|---|
| COLO829_ONT | ASCAT (T+N) | 35.33 | 575 | 1015662001 | 12864314 |
| COLO829_ONT | ASCAT (TO) | 39.31 | 290 | 1130080160 | 20362317 |
| COLO829_ONT | LongPhase-TO (ClairS-TO) | 26.85 | 90 | 771811167 | 50760310 |
| COLO829_ONT | LongPhase-TO (DeepSomatic-TO) | 27.10 | 105 | 779127888 | 50760310 |
| COLO829_NYGC | ASCAT (T+N) | 29.11 | 45 | 836961836 | 80866668 |
| COLO829_NYGC | ASCAT (TO) | 33.24 | 192 | 955772886 | 38166624 |
| COLO829_NYGC | LongPhase-TO (ClairS-TO) | 28.11 | 119 | 808144053 | 35105161 |
| COLO829_NYGC | LongPhase-TO (DeepSomatic-TO) | 29.05 | 159 | 835138542 | 45643569 |
| H1437_UCSC | ASCAT (T+N) | 39.51 | 1948 | 1135872591 | 3829891 |
| H1437_UCSC | ASCAT (TO) | 44.70 | 531 | 1285175995 | 34097583 |
| H1437_UCSC | LongPhase-TO (ClairS-TO) | 34.62 | 128 | 995275823 | 43336933 |
| H1437_UCSC | LongPhase-TO (DeepSomatic-TO) | 34.79 | 125 | 1000311306 | 34605403 |
| H2009_UCSC | ASCAT (T+N) | 43.39 | 1553 | 1247558196 | 4110937 |
| H2009_UCSC | ASCAT (TO) | 48.26 | 721 | 1387574137 | 9209663 |
| H2009_UCSC | LongPhase-TO (ClairS-TO) | 35.93 | 127 | 1033104502 | 31122631 |
| H2009_UCSC | LongPhase-TO (DeepSomatic-TO) | 36.09 | 147 | 1037482232 | 34937899 |
| HCC1395_HKU | ASCAT (T+N) | 52.01 | 3068 | 1495294320 | 2803775 |
| HCC1395_HKU | ASCAT (TO) | 65.72 | 1392 | 1889479858 | 6023307 |
| HCC1395_HKU | LongPhase-TO (ClairS-TO) | 50.84 | 198 | 1461681845 | 21772604 |
| HCC1395_HKU | LongPhase-TO (DeepSomatic-TO) | 51.53 | 202 | 1481487680 | 21597183 |
| HCC1395_NYGC | ASCAT (T+N) | 52.69 | 2134 | 1514982034 | 9642863 |
| HCC1395_NYGC | ASCAT (TO) | 59.07 | 584 | 1698371776 | 16795656 |
| HCC1395_NYGC | LongPhase-TO (ClairS-TO) | 50.74 | 220 | 1458689344 | 26501707 |
| HCC1395_NYGC | LongPhase-TO (DeepSomatic-TO) | 51.86 | 253 | 1490967218 | 29068243 |
| HCC1937_UCSC | ASCAT (T+N) | 60.44 | 4483 | 1737671187 | 2214193 |
| HCC1937_UCSC | ASCAT (TO) | 62.07 | 389 | 1784605418 | 20879693 |
| HCC1937_UCSC | LongPhase-TO (ClairS-TO) | 55.86 | 169 | 1606116347 | 40921298 |
| HCC1937_UCSC | LongPhase-TO (DeepSomatic-TO) | 56.05 | 170 | 1611305118 | 43800261 |
| HCC1954_UCSC | ASCAT (T+N) | n.a. | n.a. | n.a. | n.a. |
| HCC1954_UCSC | ASCAT (TO) | 58.72 | 1341 | 1688216927 | 2812944 |
| HCC1954_UCSC | LongPhase-TO (ClairS-TO) | 9.55 | 147 | 274660981 | 16263599 |
| HCC1954_UCSC | LongPhase-TO (DeepSomatic-TO) | 10.28 | 165 | 295496102 | 15126962 |

**Supplementary Table 4**: Somatic SNV calling performance across DNA fraction levels for eight tumor cell line datasets. Fraction is the tumor DNA fraction.

| Sample | Fraction | Variant caller | Precision | Recall | F1-score |
|---|---|---|---|---|---|
| COLO829_ONT | 1 | ClairS-TO | 0.674 | 0.799 | 0.731 |
| COLO829_ONT | 1 | ClairS-TO+LongPhase-TO | 0.666 | 0.808 | 0.730 |
| COLO829_ONT | 1 | DeepSomatic-TO | 0.501 | 0.844 | 0.629 |
| COLO829_ONT | 1 | DeepSomatic-TO+LongPhase-TO | 0.628 | 0.813 | 0.709 |
| COLO829_ONT | 0.8 | ClairS-TO | 0.651 | 0.790 | 0.714 |
| COLO829_ONT | 0.8 | ClairS-TO+LongPhase-TO | 0.857 | 0.674 | 0.755 |
| COLO829_ONT | 0.8 | DeepSomatic-TO | 0.481 | 0.849 | 0.614 |
| COLO829_ONT | 0.8 | DeepSomatic-TO+LongPhase-TO | 0.825 | 0.637 | 0.719 |
| COLO829_ONT | 0.6 | ClairS-TO | 0.623 | 0.764 | 0.686 |
| COLO829_ONT | 0.6 | ClairS-TO+LongPhase-TO | 0.848 | 0.746 | 0.794 |
| COLO829_ONT | 0.6 | DeepSomatic-TO | 0.466 | 0.842 | 0.600 |
| COLO829_ONT | 0.6 | DeepSomatic-TO+LongPhase-TO | 0.825 | 0.749 | 0.785 |
| COLO829_ONT | 0.4 | ClairS-TO | 0.576 | 0.659 | 0.615 |
| COLO829_ONT | 0.4 | ClairS-TO+LongPhase-TO | 0.814 | 0.661 | 0.730 |
| COLO829_ONT | 0.4 | DeepSomatic-TO | 0.445 | 0.797 | 0.571 |
| COLO829_ONT | 0.4 | DeepSomatic-TO+LongPhase-TO | 0.799 | 0.722 | 0.759 |
| COLO829_ONT | 0.2 | ClairS-TO | 0.392 | 0.326 | 0.356 |
| COLO829_ONT | 0.2 | ClairS-TO+LongPhase-TO | 0.668 | 0.348 | 0.458 |
| COLO829_ONT | 0.2 | DeepSomatic-TO | 0.373 | 0.604 | 0.461 |
| COLO829_ONT | 0.2 | DeepSomatic-TO+LongPhase-TO | 0.727 | 0.538 | 0.618 |
| COLO829_NYGC | 1 | ClairS-TO | 0.682 | 0.817 | 0.744 |
| COLO829_NYGC | 1 | ClairS-TO+LongPhase-TO | 0.677 | 0.832 | 0.747 |
| COLO829_NYGC | 1 | DeepSomatic-TO | 0.543 | 0.857 | 0.665 |
| COLO829_NYGC | 1 | DeepSomatic-TO+LongPhase-TO | 0.651 | 0.835 | 0.732 |
| COLO829_NYGC | 0.8 | ClairS-TO | 0.659 | 0.809 | 0.726 |
| COLO829_NYGC | 0.8 | ClairS-TO+LongPhase-TO | 0.863 | 0.633 | 0.730 |
| COLO829_NYGC | 0.8 | DeepSomatic-TO | 0.514 | 0.863 | 0.644 |

| Sample | Fraction | Variant caller | Precision | Recall | F1-score |
|---|---|---|---|---|---|
| COLO829_NYGC | 0.8 | DeepSomatic-TO+LongPhase-TO | 0.860 | 0.564 | 0.681 |
| COLO829_NYGC | 0.6 | ClairS-TO | 0.632 | 0.782 | 0.699 |
| COLO829_NYGC | 0.6 | ClairS-TO+LongPhase-TO | 0.859 | 0.678 | 0.758 |
| COLO829_NYGC | 0.6 | DeepSomatic-TO | 0.494 | 0.853 | 0.625 |
| COLO829_NYGC | 0.6 | DeepSomatic-TO+LongPhase-TO | 0.846 | 0.636 | 0.726 |
| COLO829_NYGC | 0.4 | ClairS-TO | 0.588 | 0.680 | 0.631 |
| COLO829_NYGC | 0.4 | ClairS-TO+LongPhase-TO | 0.826 | 0.610 | 0.702 |
| COLO829_NYGC | 0.4 | DeepSomatic-TO | 0.471 | 0.813 | 0.596 |
| COLO829_NYGC | 0.4 | DeepSomatic-TO+LongPhase-TO | 0.815 | 0.621 | 0.704 |
| COLO829_NYGC | 0.2 | ClairS-TO | 0.405 | 0.339 | 0.369 |
| COLO829_NYGC | 0.2 | ClairS-TO+LongPhase-TO | 0.691 | 0.321 | 0.438 |
| COLO829_NYGC | 0.2 | DeepSomatic-TO | 0.394 | 0.620 | 0.482 |
| COLO829_NYGC | 0.2 | DeepSomatic-TO+LongPhase-TO | 0.741 | 0.463 | 0.570 |
| H1437_UCSC | 1 | ClairS-TO | 0.770 | 0.588 | 0.667 |
| H1437_UCSC | 1 | ClairS-TO+LongPhase-TO | 0.753 | 0.611 | 0.675 |
| H1437_UCSC | 1 | DeepSomatic-TO | 0.680 | 0.620 | 0.649 |
| H1437_UCSC | 1 | DeepSomatic-TO+LongPhase-TO | 0.726 | 0.609 | 0.662 |
| H1437_UCSC | 0.8 | ClairS-TO | 0.728 | 0.558 | 0.632 |
| H1437_UCSC | 0.8 | ClairS-TO+LongPhase-TO | 0.871 | 0.542 | 0.668 |
| H1437_UCSC | 0.8 | DeepSomatic-TO | 0.624 | 0.620 | 0.622 |
| H1437_UCSC | 0.8 | DeepSomatic-TO+LongPhase-TO | 0.847 | 0.535 | 0.656 |
| H1437_UCSC | 0.6 | ClairS-TO | 0.721 | 0.533 | 0.613 |
| H1437_UCSC | 0.6 | ClairS-TO+LongPhase-TO | 0.831 | 0.562 | 0.671 |
| H1437_UCSC | 0.6 | DeepSomatic-TO | 0.603 | 0.607 | 0.605 |
| H1437_UCSC | 0.6 | DeepSomatic-TO+LongPhase-TO | 0.837 | 0.575 | 0.682 |
| H1437_UCSC | 0.4 | ClairS-TO | 0.730 | 0.483 | 0.581 |
| H1437_UCSC | 0.4 | ClairS-TO+LongPhase-TO | 0.805 | 0.503 | 0.619 |
| H1437_UCSC | 0.4 | DeepSomatic-TO | 0.584 | 0.570 | 0.577 |
| H1437_UCSC | 0.4 | DeepSomatic-TO+LongPhase-TO | 0.825 | 0.540 | 0.653 |
| H1437_UCSC | 0.2 | ClairS-TO | 0.836 | 0.278 | 0.417 |
| H1437_UCSC | 0.2 | ClairS-TO+LongPhase-TO | 0.718 | 0.309 | 0.432 |
| H1437_UCSC | 0.2 | DeepSomatic-TO | 0.515 | 0.436 | 0.472 |
| H1437_UCSC | 0.2 | DeepSomatic-TO+LongPhase-TO | 0.784 | 0.407 | 0.536 |
| H2009_UCSC | 1 | ClairS-TO | 0.912 | 0.913 | 0.912 |
| H2009_UCSC | 1 | ClairS-TO+LongPhase-TO | 0.904 | 0.937 | 0.920 |
| H2009_UCSC | 1 | DeepSomatic-TO | 0.861 | 0.938 | 0.898 |
| H2009_UCSC | 1 | DeepSomatic-TO+LongPhase-TO | 0.884 | 0.916 | 0.900 |
| H2009_UCSC | 0.8 | ClairS-TO | 0.884 | 0.905 | 0.895 |
| H2009_UCSC | 0.8 | ClairS-TO+LongPhase-TO | 0.942 | 0.848 | 0.893 |
| H2009_UCSC | 0.8 | DeepSomatic-TO | 0.824 | 0.946 | 0.881 |
| H2009_UCSC | 0.8 | DeepSomatic-TO+LongPhase-TO | 0.923 | 0.825 | 0.871 |
| H2009_UCSC | 0.6 | ClairS-TO | 0.855 | 0.885 | 0.870 |
| H2009_UCSC | 0.6 | ClairS-TO+LongPhase-TO | 0.920 | 0.889 | 0.904 |
| H2009_UCSC | 0.6 | DeepSomatic-TO | 0.808 | 0.938 | 0.868 |
| H2009_UCSC | 0.6 | DeepSomatic-TO+LongPhase-TO | 0.912 | 0.888 | 0.900 |
| H2009_UCSC | 0.4 | ClairS-TO | 0.830 | 0.783 | 0.805 |
| H2009_UCSC | 0.4 | ClairS-TO+LongPhase-TO | 0.903 | 0.800 | 0.848 |
| H2009_UCSC | 0.4 | DeepSomatic-TO | 0.797 | 0.899 | 0.845 |
| H2009_UCSC | 0.4 | DeepSomatic-TO+LongPhase-TO | 0.905 | 0.847 | 0.875 |
| H2009_UCSC | 0.2 | ClairS-TO | 0.696 | 0.376 | 0.488 |
| H2009_UCSC | 0.2 | ClairS-TO+LongPhase-TO | 0.837 | 0.416 | 0.555 |
| H2009_UCSC | 0.2 | DeepSomatic-TO | 0.720 | 0.616 | 0.664 |
| H2009_UCSC | 0.2 | DeepSomatic-TO+LongPhase-TO | 0.865 | 0.555 | 0.676 |
| HCC1395_HKU | 1 | ClairS-TO | 0.708 | 0.719 | 0.714 |
| HCC1395_HKU | 1 | ClairS-TO+LongPhase-TO | 0.699 | 0.729 | 0.714 |
| HCC1395_HKU | 1 | DeepSomatic-TO | 0.567 | 0.763 | 0.650 |
| HCC1395_HKU | 1 | DeepSomatic-TO+LongPhase-TO | 0.650 | 0.743 | 0.693 |
| HCC1395_HKU | 0.8 | ClairS-TO | 0.640 | 0.631 | 0.635 |
| HCC1395_HKU | 0.8 | ClairS-TO+LongPhase-TO | 0.789 | 0.664 | 0.721 |
| HCC1395_HKU | 0.8 | DeepSomatic-TO | 0.507 | 0.753 | 0.606 |
| HCC1395_HKU | 0.8 | DeepSomatic-TO+LongPhase-TO | 0.751 | 0.656 | 0.701 |
| HCC1395_HKU | 0.6 | ClairS-TO | 0.642 | 0.613 | 0.627 |
| HCC1395_HKU | 0.6 | ClairS-TO+LongPhase-TO | 0.747 | 0.651 | 0.696 |
| HCC1395_HKU | 0.6 | DeepSomatic-TO | 0.483 | 0.733 | 0.582 |
| HCC1395_HKU | 0.6 | DeepSomatic-TO+LongPhase-TO | 0.738 | 0.672 | 0.704 |

| Sample | Fraction | Variant caller | Precision | Recall | F1-score |
|---|---|---|---|---|---|
| HCC1395_HKU | 0.4 | ClairS-TO | 0.568 | 0.521 | 0.543 |
| HCC1395_HKU | 0.4 | ClairS-TO+LongPhase-TO | 0.702 | 0.538 | 0.609 |
| HCC1395_HKU | 0.4 | DeepSomatic-TO | 0.459 | 0.673 | 0.546 |
| HCC1395_HKU | 0.4 | DeepSomatic-TO+LongPhase-TO | 0.723 | 0.611 | 0.663 |
| HCC1395_HKU | 0.2 | ClairS-TO | 0.332 | 0.243 | 0.280 |
| HCC1395_HKU | 0.2 | ClairS-TO+LongPhase-TO | 0.544 | 0.268 | 0.359 |
| HCC1395_HKU | 0.2 | DeepSomatic-TO | 0.371 | 0.465 | 0.413 |
| HCC1395_HKU | 0.2 | DeepSomatic-TO+LongPhase-TO | 0.642 | 0.407 | 0.498 |
| HCC1395_NYGC | 1 | ClairS-TO | 0.712 | 0.728 | 0.720 |
| HCC1395_NYGC | 1 | ClairS-TO+LongPhase-TO | 0.695 | 0.743 | 0.718 |
| HCC1395_NYGC | 1 | DeepSomatic-TO | 0.605 | 0.786 | 0.684 |
| HCC1395_NYGC | 1 | DeepSomatic-TO+LongPhase-TO | 0.672 | 0.772 | 0.719 |
| HCC1395_NYGC | 0.8 | ClairS-TO | 0.647 | 0.710 | 0.677 |
| HCC1395_NYGC | 0.8 | ClairS-TO+LongPhase-TO | 0.830 | 0.610 | 0.703 |
| HCC1395_NYGC | 0.8 | DeepSomatic-TO | 0.536 | 0.773 | 0.633 |
| HCC1395_NYGC | 0.8 | DeepSomatic-TO+LongPhase-TO | 0.819 | 0.567 | 0.670 |
| HCC1395_NYGC | 0.6 | ClairS-TO | 0.588 | 0.666 | 0.625 |
| HCC1395_NYGC | 0.6 | ClairS-TO+LongPhase-TO | 0.794 | 0.607 | 0.688 |
| HCC1395_NYGC | 0.6 | DeepSomatic-TO | 0.508 | 0.749 | 0.606 |
| HCC1395_NYGC | 0.6 | DeepSomatic-TO+LongPhase-TO | 0.808 | 0.593 | 0.684 |
| HCC1395_NYGC | 0.4 | ClairS-TO | 0.529 | 0.544 | 0.536 |
| HCC1395_NYGC | 0.4 | ClairS-TO+LongPhase-TO | 0.756 | 0.507 | 0.607 |
| HCC1395_NYGC | 0.4 | DeepSomatic-TO | 0.484 | 0.689 | 0.568 |
| HCC1395_NYGC | 0.4 | DeepSomatic-TO+LongPhase-TO | 0.794 | 0.539 | 0.643 |
| HCC1395_NYGC | 0.2 | ClairS-TO | 0.324 | 0.234 | 0.272 |
| HCC1395_NYGC | 0.2 | ClairS-TO+LongPhase-TO | 0.610 | 0.231 | 0.335 |
| HCC1395_NYGC | 0.2 | DeepSomatic-TO | 0.362 | 0.437 | 0.396 |
| HCC1395_NYGC | 0.2 | DeepSomatic-TO+LongPhase-TO | 0.694 | 0.325 | 0.443 |
| HCC1937_UCSC | 1 | ClairS-TO | 0.483 | 0.748 | 0.587 |
| HCC1937_UCSC | 1 | ClairS-TO+LongPhase-TO | 0.462 | 0.774 | 0.579 |
| HCC1937_UCSC | 1 | DeepSomatic-TO | 0.379 | 0.834 | 0.521 |
| HCC1937_UCSC | 1 | DeepSomatic-TO+LongPhase-TO | 0.437 | 0.816 | 0.569 |
| HCC1937_UCSC | 0.8 | ClairS-TO | 0.409 | 0.692 | 0.514 |
| HCC1937_UCSC | 0.8 | ClairS-TO+LongPhase-TO | 0.685 | 0.687 | 0.686 |
| HCC1937_UCSC | 0.8 | DeepSomatic-TO | 0.312 | 0.791 | 0.448 |
| HCC1937_UCSC | 0.8 | DeepSomatic-TO+LongPhase-TO | 0.636 | 0.717 | 0.674 |
| HCC1937_UCSC | 0.6 | ClairS-TO | 0.338 | 0.608 | 0.434 |
| HCC1937_UCSC | 0.6 | ClairS-TO+LongPhase-TO | 0.620 | 0.635 | 0.627 |
| HCC1937_UCSC | 0.6 | DeepSomatic-TO | 0.275 | 0.724 | 0.399 |
| HCC1937_UCSC | 0.6 | DeepSomatic-TO+LongPhase-TO | 0.604 | 0.685 | 0.642 |
| HCC1937_UCSC | 0.4 | ClairS-TO | 0.268 | 0.454 | 0.337 |
| HCC1937_UCSC | 0.4 | ClairS-TO+LongPhase-TO | 0.538 | 0.487 | 0.511 |
| HCC1937_UCSC | 0.4 | DeepSomatic-TO | 0.228 | 0.589 | 0.329 |
| HCC1937_UCSC | 0.4 | DeepSomatic-TO+LongPhase-TO | 0.529 | 0.552 | 0.540 |
| HCC1937_UCSC | 0.2 | ClairS-TO | 0.128 | 0.184 | 0.151 |
| HCC1937_UCSC | 0.2 | ClairS-TO+LongPhase-TO | 0.348 | 0.220 | 0.270 |
| HCC1937_UCSC | 0.2 | DeepSomatic-TO | 0.131 | 0.309 | 0.184 |
| HCC1937_UCSC | 0.2 | DeepSomatic-TO+LongPhase-TO | 0.360 | 0.276 | 0.312 |
| HCC1954_UCSC | 1 | ClairS-TO | 0.248 | 0.799 | 0.379 |
| HCC1954_UCSC | 1 | ClairS-TO+LongPhase-TO | 0.254 | 0.834 | 0.389 |
| HCC1954_UCSC | 1 | DeepSomatic-TO | 0.214 | 0.888 | 0.345 |
| HCC1954_UCSC | 1 | DeepSomatic-TO+LongPhase-TO | 0.260 | 0.872 | 0.401 |
| HCC1954_UCSC | 0.8 | ClairS-TO | 0.218 | 0.718 | 0.334 |
| HCC1954_UCSC | 0.8 | ClairS-TO+LongPhase-TO | 0.521 | 0.733 | 0.609 |
| HCC1954_UCSC | 0.8 | DeepSomatic-TO | 0.196 | 0.854 | 0.319 |
| HCC1954_UCSC | 0.8 | DeepSomatic-TO+LongPhase-TO | 0.616 | 0.774 | 0.686 |
| HCC1954_UCSC | 0.6 | ClairS-TO | 0.175 | 0.577 | 0.269 |
| HCC1954_UCSC | 0.6 | ClairS-TO+LongPhase-TO | 0.448 | 0.617 | 0.519 |
| HCC1954_UCSC | 0.6 | DeepSomatic-TO | 0.179 | 0.782 | 0.292 |
| HCC1954_UCSC | 0.6 | DeepSomatic-TO+LongPhase-TO | 0.579 | 0.723 | 0.643 |
| HCC1954_UCSC | 0.4 | ClairS-TO | 0.117 | 0.364 | 0.177 |
| HCC1954_UCSC | 0.4 | ClairS-TO+LongPhase-TO | 0.340 | 0.406 | 0.370 |
| HCC1954_UCSC | 0.4 | DeepSomatic-TO | 0.147 | 0.619 | 0.237 |
| HCC1954_UCSC | 0.4 | DeepSomatic-TO+LongPhase-TO | 0.514 | 0.565 | 0.538 |
| HCC1954_UCSC | 0.2 | ClairS-TO | 0.048 | 0.140 | 0.072 |

| Sample | Fraction | Variant caller | Precision | Recall | F1-score |
|---|---|---|---|---|---|
| HCC1954_UCSC | 0.2 | ClairS-TO+LongPhase-TO | 0.180 | 0.168 | 0.174 |
| HCC1954_UCSC | 0.2 | DeepSomatic-TO | 0.087 | 0.347 | 0.139 |
| HCC1954_UCSC | 0.2 | DeepSomatic-TO+LongPhase-TO | 0.358 | 0.309 | 0.332 |

**Supplementary Table 5**: Somatic indel calling performance across DNA fraction levels for eight tumor cell line datasets. Fraction is the tumor DNA fraction.

| Sample | Fraction | Variant caller | Precision | Recall | F1-score |
|---|---|---|---|---|---|
| COLO829_ONT | 1 | ClairS-TO | 0.154 | 0.226 | 0.183 |
| COLO829_ONT | 1 | ClairS-TO+LongPhase-TO | 0.127 | 0.301 | 0.179 |
| COLO829_ONT | 1 | DeepSomatic-TO | 0.072 | 0.115 | 0.088 |
| COLO829_ONT | 1 | DeepSomatic-TO+LongPhase-TO | 0.165 | 0.108 | 0.130 |
| COLO829_ONT | 0.8 | ClairS-TO | 0.154 | 0.210 | 0.178 |
| COLO829_ONT | 0.8 | ClairS-TO+LongPhase-TO | 0.225 | 0.190 | 0.206 |
| COLO829_ONT | 0.8 | DeepSomatic-TO | 0.080 | 0.135 | 0.100 |
| COLO829_ONT | 0.8 | DeepSomatic-TO+LongPhase-TO | 0.282 | 0.084 | 0.130 |
| COLO829_ONT | 0.6 | ClairS-TO | 0.111 | 0.148 | 0.127 |
| COLO829_ONT | 0.6 | ClairS-TO+LongPhase-TO | 0.222 | 0.181 | 0.199 |
| COLO829_ONT | 0.6 | DeepSomatic-TO | 0.074 | 0.131 | 0.095 |
| COLO829_ONT | 0.6 | DeepSomatic-TO+LongPhase-TO | 0.356 | 0.106 | 0.163 |
| COLO829_ONT | 0.4 | ClairS-TO | 0.056 | 0.075 | 0.064 |
| COLO829_ONT | 0.4 | ClairS-TO+LongPhase-TO | 0.125 | 0.099 | 0.110 |
| COLO829_ONT | 0.4 | DeepSomatic-TO | 0.056 | 0.101 | 0.072 |
| COLO829_ONT | 0.4 | DeepSomatic-TO+LongPhase-TO | 0.310 | 0.091 | 0.141 |
| COLO829_ONT | 0.2 | ClairS-TO | 0.005 | 0.006 | 0.005 |
| COLO829_ONT | 0.2 | ClairS-TO+LongPhase-TO | 0.016 | 0.011 | 0.013 |
| COLO829_ONT | 0.2 | DeepSomatic-TO | 0.025 | 0.044 | 0.032 |
| COLO829_ONT | 0.2 | DeepSomatic-TO+LongPhase-TO | 0.174 | 0.041 | 0.066 |
| COLO829_NYGC | 1 | ClairS-TO | 0.177 | 0.243 | 0.205 |
| COLO829_NYGC | 1 | ClairS-TO+LongPhase-TO | 0.141 | 0.324 | 0.197 |
| COLO829_NYGC | 1 | DeepSomatic-TO | 0.044 | 0.114 | 0.063 |
| COLO829_NYGC | 1 | DeepSomatic-TO+LongPhase-TO | 0.140 | 0.108 | 0.122 |
| COLO829_NYGC | 0.8 | ClairS-TO | 0.149 | 0.206 | 0.173 |
| COLO829_NYGC | 0.8 | ClairS-TO+LongPhase-TO | 0.213 | 0.207 | 0.210 |
| COLO829_NYGC | 0.8 | DeepSomatic-TO | 0.052 | 0.145 | 0.076 |
| COLO829_NYGC | 0.8 | DeepSomatic-TO+LongPhase-TO | 0.269 | 0.092 | 0.138 |
| COLO829_NYGC | 0.6 | ClairS-TO | 0.113 | 0.162 | 0.133 |
| COLO829_NYGC | 0.6 | ClairS-TO+LongPhase-TO | 0.190 | 0.183 | 0.186 |
| COLO829_NYGC | 0.6 | DeepSomatic-TO | 0.045 | 0.137 | 0.068 |
| COLO829_NYGC | 0.6 | DeepSomatic-TO+LongPhase-TO | 0.288 | 0.107 | 0.156 |
| COLO829_NYGC | 0.4 | ClairS-TO | 0.051 | 0.073 | 0.060 |
| COLO829_NYGC | 0.4 | ClairS-TO+LongPhase-TO | 0.104 | 0.098 | 0.101 |
| COLO829_NYGC | 0.4 | DeepSomatic-TO | 0.035 | 0.110 | 0.053 |
| COLO829_NYGC | 0.4 | DeepSomatic-TO+LongPhase-TO | 0.259 | 0.085 | 0.128 |
| COLO829_NYGC | 0.2 | ClairS-TO | 0.004 | 0.006 | 0.005 |
| COLO829_NYGC | 0.2 | ClairS-TO+LongPhase-TO | 0.011 | 0.009 | 0.010 |
| COLO829_NYGC | 0.2 | DeepSomatic-TO | 0.013 | 0.041 | 0.019 |
| COLO829_NYGC | 0.2 | DeepSomatic-TO+LongPhase-TO | 0.106 | 0.030 | 0.047 |
| H1437_UCSC | 1 | ClairS-TO | 0.569 | 0.279 | 0.375 |
| H1437_UCSC | 1 | ClairS-TO+LongPhase-TO | 0.511 | 0.343 | 0.410 |
| H1437_UCSC | 1 | DeepSomatic-TO | 0.304 | 0.110 | 0.162 |
| H1437_UCSC | 1 | DeepSomatic-TO+LongPhase-TO | 0.443 | 0.106 | 0.171 |
| H1437_UCSC | 0.8 | ClairS-TO | 0.527 | 0.233 | 0.323 |
| H1437_UCSC | 0.8 | ClairS-TO+LongPhase-TO | 0.628 | 0.217 | 0.323 |
| H1437_UCSC | 0.8 | DeepSomatic-TO | 0.319 | 0.135 | 0.190 |
| H1437_UCSC | 0.8 | DeepSomatic-TO+LongPhase-TO | 0.635 | 0.096 | 0.167 |
| H1437_UCSC | 0.6 | ClairS-TO | 0.504 | 0.170 | 0.254 |
| H1437_UCSC | 0.6 | ClairS-TO+LongPhase-TO | 0.601 | 0.203 | 0.304 |
| H1437_UCSC | 0.6 | DeepSomatic-TO | 0.288 | 0.132 | 0.181 |
| H1437_UCSC | 0.6 | DeepSomatic-TO+LongPhase-TO | 0.623 | 0.116 | 0.196 |
| H1437_UCSC | 0.4 | ClairS-TO | 0.307 | 0.073 | 0.117 |
| H1437_UCSC | 0.4 | ClairS-TO+LongPhase-TO | 0.369 | 0.105 | 0.163 |
| H1437_UCSC | 0.4 | DeepSomatic-TO | 0.218 | 0.095 | 0.132 |

| Sample | Fraction | Variant caller | Precision | Recall | F1-score |
|---|---|---|---|---|---|
| H1437_UCSC | 0.4 | DeepSomatic-TO+LongPhase-TO | 0.552 | 0.088 | 0.152 |
| H1437_UCSC | 0.2 | ClairS-TO | 0.146 | 0.008 | 0.014 |
| H1437_UCSC | 0.2 | ClairS-TO+LongPhase-TO | 0.053 | 0.012 | 0.019 |
| H1437_UCSC | 0.2 | DeepSomatic-TO | 0.061 | 0.023 | 0.033 |
| H1437_UCSC | 0.2 | DeepSomatic-TO+LongPhase-TO | 0.230 | 0.022 | 0.039 |
| H2009_UCSC | 1 | ClairS-TO | 0.764 | 0.570 | 0.653 |
| H2009_UCSC | 1 | ClairS-TO+LongPhase-TO | 0.721 | 0.672 | 0.696 |
| H2009_UCSC | 1 | DeepSomatic-TO | 0.477 | 0.185 | 0.267 |
| H2009_UCSC | 1 | DeepSomatic-TO+LongPhase-TO | 0.630 | 0.180 | 0.279 |
| H2009_UCSC | 0.8 | ClairS-TO | 0.721 | 0.467 | 0.567 |
| H2009_UCSC | 0.8 | ClairS-TO+LongPhase-TO | 0.806 | 0.429 | 0.560 |
| H2009_UCSC | 0.8 | DeepSomatic-TO | 0.500 | 0.227 | 0.312 |
| H2009_UCSC | 0.8 | DeepSomatic-TO+LongPhase-TO | 0.777 | 0.173 | 0.283 |
| H2009_UCSC | 0.6 | ClairS-TO | 0.613 | 0.319 | 0.419 |
| H2009_UCSC | 0.6 | ClairS-TO+LongPhase-TO | 0.753 | 0.362 | 0.489 |
| H2009_UCSC | 0.6 | DeepSomatic-TO | 0.454 | 0.218 | 0.295 |
| H2009_UCSC | 0.6 | DeepSomatic-TO+LongPhase-TO | 0.765 | 0.191 | 0.306 |
| H2009_UCSC | 0.4 | ClairS-TO | 0.400 | 0.152 | 0.221 |
| H2009_UCSC | 0.4 | ClairS-TO+LongPhase-TO | 0.571 | 0.197 | 0.293 |
| H2009_UCSC | 0.4 | DeepSomatic-TO | 0.356 | 0.160 | 0.221 |
| H2009_UCSC | 0.4 | DeepSomatic-TO+LongPhase-TO | 0.681 | 0.148 | 0.244 |
| H2009_UCSC | 0.2 | ClairS-TO | 0.053 | 0.015 | 0.023 |
| H2009_UCSC | 0.2 | ClairS-TO+LongPhase-TO | 0.130 | 0.024 | 0.041 |
| H2009_UCSC | 0.2 | DeepSomatic-TO | 0.092 | 0.034 | 0.050 |
| H2009_UCSC | 0.2 | DeepSomatic-TO+LongPhase-TO | 0.311 | 0.031 | 0.056 |
| HCC1395_HKU | 1 | ClairS-TO | 0.376 | 0.357 | 0.366 |
| HCC1395_HKU | 1 | ClairS-TO+LongPhase-TO | 0.293 | 0.445 | 0.353 |
| HCC1395_HKU | 1 | DeepSomatic-TO | 0.376 | 0.536 | 0.442 |
| HCC1395_HKU | 1 | DeepSomatic-TO+LongPhase-TO | 0.499 | 0.513 | 0.506 |
| HCC1395_HKU | 0.8 | ClairS-TO | 0.326 | 0.256 | 0.287 |
| HCC1395_HKU | 0.8 | ClairS-TO+LongPhase-TO | 0.397 | 0.329 | 0.360 |
| HCC1395_HKU | 0.8 | DeepSomatic-TO | 0.341 | 0.515 | 0.410 |
| HCC1395_HKU | 0.8 | DeepSomatic-TO+LongPhase-TO | 0.582 | 0.422 | 0.489 |
| HCC1395_HKU | 0.6 | ClairS-TO | 0.294 | 0.175 | 0.220 |
| HCC1395_HKU | 0.6 | ClairS-TO+LongPhase-TO | 0.216 | 0.255 | 0.234 |
| HCC1395_HKU | 0.6 | DeepSomatic-TO | 0.293 | 0.453 | 0.355 |
| HCC1395_HKU | 0.6 | DeepSomatic-TO+LongPhase-TO | 0.584 | 0.384 | 0.464 |
| HCC1395_HKU | 0.4 | ClairS-TO | 0.109 | 0.066 | 0.082 |
| HCC1395_HKU | 0.4 | ClairS-TO+LongPhase-TO | 0.179 | 0.107 | 0.134 |
| HCC1395_HKU | 0.4 | DeepSomatic-TO | 0.207 | 0.317 | 0.251 |
| HCC1395_HKU | 0.4 | DeepSomatic-TO+LongPhase-TO | 0.520 | 0.277 | 0.362 |
| HCC1395_HKU | 0.2 | ClairS-TO | 0.007 | 0.005 | 0.006 |
| HCC1395_HKU | 0.2 | ClairS-TO+LongPhase-TO | 0.023 | 0.012 | 0.016 |
| HCC1395_HKU | 0.2 | DeepSomatic-TO | 0.077 | 0.100 | 0.087 |
| HCC1395_HKU | 0.2 | DeepSomatic-TO+LongPhase-TO | 0.283 | 0.085 | 0.131 |
| HCC1395_NYGC | 1 | ClairS-TO | 0.434 | 0.401 | 0.417 |
| HCC1395_NYGC | 1 | ClairS-TO+LongPhase-TO | 0.342 | 0.488 | 0.403 |
| HCC1395_NYGC | 1 | DeepSomatic-TO | 0.464 | 0.630 | 0.534 |
| HCC1395_NYGC | 1 | DeepSomatic-TO+LongPhase-TO | 0.587 | 0.598 | 0.593 |
| HCC1395_NYGC | 0.8 | ClairS-TO | 0.403 | 0.328 | 0.362 |
| HCC1395_NYGC | 0.8 | ClairS-TO+LongPhase-TO | 0.467 | 0.344 | 0.396 |
| HCC1395_NYGC | 0.8 | DeepSomatic-TO | 0.426 | 0.578 | 0.491 |
| HCC1395_NYGC | 0.8 | DeepSomatic-TO+LongPhase-TO | 0.685 | 0.438 | 0.534 |
| HCC1395_NYGC | 0.6 | ClairS-TO | 0.299 | 0.220 | 0.253 |
| HCC1395_NYGC | 0.6 | ClairS-TO+LongPhase-TO | 0.428 | 0.255 | 0.320 |
| HCC1395_NYGC | 0.6 | DeepSomatic-TO | 0.348 | 0.496 | 0.409 |
| HCC1395_NYGC | 0.6 | DeepSomatic-TO+LongPhase-TO | 0.631 | 0.391 | 0.483 |
| HCC1395_NYGC | 0.4 | ClairS-TO | 0.134 | 0.093 | 0.110 |
| HCC1395_NYGC | 0.4 | ClairS-TO+LongPhase-TO | 0.251 | 0.120 | 0.162 |
| HCC1395_NYGC | 0.4 | DeepSomatic-TO | 0.258 | 0.341 | 0.294 |
| HCC1395_NYGC | 0.4 | DeepSomatic-TO+LongPhase-TO | 0.572 | 0.267 | 0.364 |
| HCC1395_NYGC | 0.2 | ClairS-TO | 0.011 | 0.007 | 0.009 |
| HCC1395_NYGC | 0.2 | ClairS-TO+LongPhase-TO | 0.035 | 0.015 | 0.021 |
| HCC1395_NYGC | 0.2 | DeepSomatic-TO | 0.092 | 0.103 | 0.097 |
| HCC1395_NYGC | 0.2 | DeepSomatic-TO+LongPhase-TO | 0.315 | 0.078 | 0.125 |

| Sample | Fraction | Variant caller | Precision | Recall | F1-score |
|---|---|---|---|---|---|
| HCC1937_UCSC | 1 | ClairS-TO | 0.272 | 0.413 | 0.328 |
| HCC1937_UCSC | 1 | ClairS-TO+LongPhase-TO | 0.203 | 0.535 | 0.294 |
| HCC1937_UCSC | 1 | DeepSomatic-TO | 0.219 | 0.378 | 0.277 |
| HCC1937_UCSC | 1 | DeepSomatic-TO+LongPhase-TO | 0.297 | 0.363 | 0.327 |
| HCC1937_UCSC | 0.8 | ClairS-TO | 0.285 | 0.391 | 0.330 |
| HCC1937_UCSC | 0.8 | ClairS-TO+LongPhase-TO | 0.394 | 0.401 | 0.397 |
| HCC1937_UCSC | 0.8 | DeepSomatic-TO | 0.246 | 0.432 | 0.313 |
| HCC1937_UCSC | 0.8 | DeepSomatic-TO+LongPhase-TO | 0.457 | 0.337 | 0.388 |
| HCC1937_UCSC | 0.6 | ClairS-TO | 0.240 | 0.320 | 0.275 |
| HCC1937_UCSC | 0.6 | ClairS-TO+LongPhase-TO | 0.424 | 0.368 | 0.394 |
| HCC1937_UCSC | 0.6 | DeepSomatic-TO | 0.224 | 0.417 | 0.292 |
| HCC1937_UCSC | 0.6 | DeepSomatic-TO+LongPhase-TO | 0.524 | 0.384 | 0.443 |
| HCC1937_UCSC | 0.4 | ClairS-TO | 0.117 | 0.160 | 0.135 |
| HCC1937_UCSC | 0.4 | ClairS-TO+LongPhase-TO | 0.247 | 0.207 | 0.225 |
| HCC1937_UCSC | 0.4 | DeepSomatic-TO | 0.162 | 0.309 | 0.212 |
| HCC1937_UCSC | 0.4 | DeepSomatic-TO+LongPhase-TO | 0.458 | 0.285 | 0.351 |
| HCC1937_UCSC | 0.2 | ClairS-TO | 0.017 | 0.024 | 0.020 |
| HCC1937_UCSC | 0.2 | ClairS-TO+LongPhase-TO | 0.054 | 0.047 | 0.050 |
| HCC1937_UCSC | 0.2 | DeepSomatic-TO | 0.063 | 0.113 | 0.081 |
| HCC1937_UCSC | 0.2 | DeepSomatic-TO+LongPhase-TO | 0.262 | 0.095 | 0.139 |
| HCC1954_UCSC | 1 | ClairS-TO | 0.165 | 0.212 | 0.186 |
| HCC1954_UCSC | 1 | ClairS-TO+LongPhase-TO | 0.169 | 0.292 | 0.214 |
| HCC1954_UCSC | 1 | DeepSomatic-TO | 0.145 | 0.135 | 0.140 |
| HCC1954_UCSC | 1 | DeepSomatic-TO+LongPhase-TO | 0.216 | 0.130 | 0.162 |
| HCC1954_UCSC | 0.8 | ClairS-TO | 0.100 | 0.128 | 0.112 |
| HCC1954_UCSC | 0.8 | ClairS-TO+LongPhase-TO | 0.249 | 0.171 | 0.203 |
| HCC1954_UCSC | 0.8 | DeepSomatic-TO | 0.113 | 0.104 | 0.109 |
| HCC1954_UCSC | 0.8 | DeepSomatic-TO+LongPhase-TO | 0.389 | 0.090 | 0.146 |
| HCC1954_UCSC | 0.6 | ClairS-TO | 0.052 | 0.067 | 0.058 |
| HCC1954_UCSC | 0.6 | ClairS-TO+LongPhase-TO | 0.134 | 0.090 | 0.108 |
| HCC1954_UCSC | 0.6 | DeepSomatic-TO | 0.080 | 0.075 | 0.077 |
| HCC1954_UCSC | 0.6 | DeepSomatic-TO+LongPhase-TO | 0.308 | 0.067 | 0.110 |
| HCC1954_UCSC | 0.4 | ClairS-TO | 0.014 | 0.019 | 0.016 |
| HCC1954_UCSC | 0.4 | ClairS-TO+LongPhase-TO | 0.040 | 0.028 | 0.033 |
| HCC1954_UCSC | 0.4 | DeepSomatic-TO | 0.040 | 0.037 | 0.038 |
| HCC1954_UCSC | 0.4 | DeepSomatic-TO+LongPhase-TO | 0.167 | 0.033 | 0.055 |
| HCC1954_UCSC | 0.2 | ClairS-TO | 0.002 | 0.003 | 0.002 |
| HCC1954_UCSC | 0.2 | ClairS-TO+LongPhase-TO | 0.005 | 0.004 | 0.004 |
| HCC1954_UCSC | 0.2 | DeepSomatic-TO | 0.009 | 0.009 | 0.009 |
| HCC1954_UCSC | 0.2 | DeepSomatic-TO+LongPhase-TO | 0.045 | 0.008 | 0.014 |

**Supplementary Table 6**: Incremental effect of methylation-aware recalibration on somatic SNV and indel calling, for the five datasets with 5-methylcytosine calls available. Fraction is the tumor DNA fraction. LP-TO denotes LongPhase-TO; (hap) is haplotype-aware recalibration alone and (hap+5mC) adds the methylation-aware step.

| Sample | Type | Fraction | Variant caller | Precision | Recall | F1-score |
|---|---|---|---|---|---|---|
| H1437_UCSC | SNV | 1 | ClairS-TO | 0.770 | 0.588 | 0.667 |
| H1437_UCSC | SNV | 1 | ClairS-TO+LP (hap) | 0.753 | 0.611 | 0.675 |
| H1437_UCSC | SNV | 1 | ClairS-TO+LP (hap+5mC) | 0.753 | 0.611 | 0.675 |
| H1437_UCSC | SNV | 1 | DeepSomatic-TO | 0.680 | 0.620 | 0.649 |
| H1437_UCSC | SNV | 1 | DeepSomatic-TO+LP (hap) | 0.726 | 0.609 | 0.662 |
| H1437_UCSC | SNV | 1 | DeepSomatic-TO+LP (hap+5mC) | 0.726 | 0.609 | 0.662 |
| H1437_UCSC | SNV | 0.8 | ClairS-TO | 0.728 | 0.558 | 0.632 |
| H1437_UCSC | SNV | 0.8 | ClairS-TO+LP (hap) | 0.871 | 0.542 | 0.668 |
| H1437_UCSC | SNV | 0.8 | ClairS-TO+LP (hap+5mC) | 0.871 | 0.541 | 0.667 |
| H1437_UCSC | SNV | 0.8 | DeepSomatic-TO | 0.624 | 0.620 | 0.622 |
| H1437_UCSC | SNV | 0.8 | DeepSomatic-TO+LP (hap) | 0.847 | 0.535 | 0.656 |
| H1437_UCSC | SNV | 0.8 | DeepSomatic-TO+LP (hap+5mC) | 0.848 | 0.534 | 0.655 |
| H1437_UCSC | SNV | 0.6 | ClairS-TO | 0.721 | 0.533 | 0.613 |
| H1437_UCSC | SNV | 0.6 | ClairS-TO+LP (hap) | 0.831 | 0.562 | 0.671 |
| H1437_UCSC | SNV | 0.6 | ClairS-TO+LP (hap+5mC) | 0.856 | 0.522 | 0.649 |
| H1437_UCSC | SNV | 0.6 | DeepSomatic-TO | 0.603 | 0.607 | 0.605 |
| H1437_UCSC | SNV | 0.6 | DeepSomatic-TO+LP (hap) | 0.837 | 0.575 | 0.682 |
| H1437_UCSC | SNV | 0.6 | DeepSomatic-TO+LP (hap+5mC) | 0.847 | 0.536 | 0.656 |
| H1437_UCSC | SNV | 0.4 | ClairS-TO | 0.730 | 0.483 | 0.581 |
| H1437_UCSC | SNV | 0.4 | ClairS-TO+LP (hap) | 0.805 | 0.503 | 0.619 |
| H1437_UCSC | SNV | 0.4 | ClairS-TO+LP (hap+5mC) | 0.844 | 0.495 | 0.624 |
| H1437_UCSC | SNV | 0.4 | DeepSomatic-TO | 0.584 | 0.570 | 0.577 |

| Sample | Type | Fraction | Variant caller | Precision | Recall | F1-score |
|---|---|---|---|---|---|---|
| H1437_UCSC | SNV | 0.4 | DeepSomatic-TO+LP (hap) | 0.825 | 0.540 | 0.653 |
| H1437_UCSC | SNV | 0.4 | DeepSomatic-TO+LP (hap+5mC) | 0.847 | 0.532 | 0.653 |
| H1437_UCSC | SNV | 0.2 | ClairS-TO | 0.836 | 0.278 | 0.417 |
| H1437_UCSC | SNV | 0.2 | ClairS-TO+LP (hap) | 0.718 | 0.309 | 0.432 |
| H1437_UCSC | SNV | 0.2 | ClairS-TO+LP (hap+5mC) | 0.790 | 0.308 | 0.444 |
| H1437_UCSC | SNV | 0.2 | DeepSomatic-TO | 0.515 | 0.436 | 0.472 |
| H1437_UCSC | SNV | 0.2 | DeepSomatic-TO+LP (hap) | 0.784 | 0.407 | 0.536 |
| H1437_UCSC | SNV | 0.2 | DeepSomatic-TO+LP (hap+5mC) | 0.823 | 0.406 | 0.543 |
| H2009_UCSC | SNV | 1 | ClairS-TO | 0.912 | 0.913 | 0.912 |
| H2009_UCSC | SNV | 1 | ClairS-TO+LP (hap) | 0.904 | 0.937 | 0.920 |
| H2009_UCSC | SNV | 1 | ClairS-TO+LP (hap+5mC) | 0.904 | 0.937 | 0.920 |
| H2009_UCSC | SNV | 1 | DeepSomatic-TO | 0.861 | 0.938 | 0.898 |
| H2009_UCSC | SNV | 1 | DeepSomatic-TO+LP (hap) | 0.884 | 0.916 | 0.900 |
| H2009_UCSC | SNV | 1 | DeepSomatic-TO+LP (hap+5mC) | 0.884 | 0.916 | 0.900 |
| H2009_UCSC | SNV | 0.8 | ClairS-TO | 0.884 | 0.905 | 0.895 |
| H2009_UCSC | SNV | 0.8 | ClairS-TO+LP (hap) | 0.942 | 0.848 | 0.893 |
| H2009_UCSC | SNV | 0.8 | ClairS-TO+LP (hap+5mC) | 0.942 | 0.847 | 0.892 |
| H2009_UCSC | SNV | 0.8 | DeepSomatic-TO | 0.824 | 0.946 | 0.881 |
| H2009_UCSC | SNV | 0.8 | DeepSomatic-TO+LP (hap) | 0.923 | 0.825 | 0.871 |
| H2009_UCSC | SNV | 0.8 | DeepSomatic-TO+LP (hap+5mC) | 0.923 | 0.824 | 0.871 |
| H2009_UCSC | SNV | 0.6 | ClairS-TO | 0.855 | 0.885 | 0.870 |
| H2009_UCSC | SNV | 0.6 | ClairS-TO+LP (hap) | 0.920 | 0.889 | 0.904 |
| H2009_UCSC | SNV | 0.6 | ClairS-TO+LP (hap+5mC) | 0.939 | 0.856 | 0.895 |
| H2009_UCSC | SNV | 0.6 | DeepSomatic-TO | 0.808 | 0.938 | 0.868 |
| H2009_UCSC | SNV | 0.6 | DeepSomatic-TO+LP (hap) | 0.912 | 0.888 | 0.900 |
| H2009_UCSC | SNV | 0.6 | DeepSomatic-TO+LP (hap+5mC) | 0.926 | 0.857 | 0.890 |
| H2009_UCSC | SNV | 0.4 | ClairS-TO | 0.830 | 0.783 | 0.805 |
| H2009_UCSC | SNV | 0.4 | ClairS-TO+LP (hap) | 0.903 | 0.800 | 0.848 |
| H2009_UCSC | SNV | 0.4 | ClairS-TO+LP (hap+5mC) | 0.925 | 0.794 | 0.854 |
| H2009_UCSC | SNV | 0.4 | DeepSomatic-TO | 0.797 | 0.899 | 0.845 |
| H2009_UCSC | SNV | 0.4 | DeepSomatic-TO+LP (hap) | 0.905 | 0.847 | 0.875 |
| H2009_UCSC | SNV | 0.4 | DeepSomatic-TO+LP (hap+5mC) | 0.921 | 0.840 | 0.879 |
| H2009_UCSC | SNV | 0.2 | ClairS-TO | 0.696 | 0.376 | 0.488 |
| H2009_UCSC | SNV | 0.2 | ClairS-TO+LP (hap) | 0.837 | 0.416 | 0.555 |
| H2009_UCSC | SNV | 0.2 | ClairS-TO+LP (hap+5mC) | 0.890 | 0.414 | 0.565 |
| H2009_UCSC | SNV | 0.2 | DeepSomatic-TO | 0.720 | 0.616 | 0.664 |
| H2009_UCSC | SNV | 0.2 | DeepSomatic-TO+LP (hap) | 0.865 | 0.555 | 0.676 |
| H2009_UCSC | SNV | 0.2 | DeepSomatic-TO+LP (hap+5mC) | 0.899 | 0.552 | 0.684 |
| HCC1395_NYGC | SNV | 1 | ClairS-TO | 0.712 | 0.728 | 0.720 |
| HCC1395_NYGC | SNV | 1 | ClairS-TO+LP (hap) | 0.695 | 0.743 | 0.718 |
| HCC1395_NYGC | SNV | 1 | ClairS-TO+LP (hap+5mC) | 0.695 | 0.743 | 0.718 |
| HCC1395_NYGC | SNV | 1 | DeepSomatic-TO | 0.605 | 0.786 | 0.684 |
| HCC1395_NYGC | SNV | 1 | DeepSomatic-TO+LP (hap) | 0.672 | 0.772 | 0.719 |
| HCC1395_NYGC | SNV | 1 | DeepSomatic-TO+LP (hap+5mC) | 0.672 | 0.772 | 0.719 |
| HCC1395_NYGC | SNV | 0.8 | ClairS-TO | 0.647 | 0.710 | 0.677 |
| HCC1395_NYGC | SNV | 0.8 | ClairS-TO+LP (hap) | 0.830 | 0.610 | 0.703 |
| HCC1395_NYGC | SNV | 0.8 | ClairS-TO+LP (hap+5mC) | 0.831 | 0.604 | 0.699 |
| HCC1395_NYGC | SNV | 0.8 | DeepSomatic-TO | 0.536 | 0.773 | 0.633 |
| HCC1395_NYGC | SNV | 0.8 | DeepSomatic-TO+LP (hap) | 0.819 | 0.567 | 0.670 |
| HCC1395_NYGC | SNV | 0.8 | DeepSomatic-TO+LP (hap+5mC) | 0.820 | 0.566 | 0.670 |
| HCC1395_NYGC | SNV | 0.6 | ClairS-TO | 0.588 | 0.666 | 0.625 |
| HCC1395_NYGC | SNV | 0.6 | ClairS-TO+LP (hap) | 0.794 | 0.607 | 0.688 |
| HCC1395_NYGC | SNV | 0.6 | ClairS-TO+LP (hap+5mC) | 0.866 | 0.566 | 0.685 |
| HCC1395_NYGC | SNV | 0.6 | DeepSomatic-TO | 0.508 | 0.749 | 0.606 |
| HCC1395_NYGC | SNV | 0.6 | DeepSomatic-TO+LP (hap) | 0.808 | 0.593 | 0.684 |
| HCC1395_NYGC | SNV | 0.6 | DeepSomatic-TO+LP (hap+5mC) | 0.848 | 0.555 | 0.671 |
| HCC1395_NYGC | SNV | 0.4 | ClairS-TO | 0.529 | 0.544 | 0.536 |
| HCC1395_NYGC | SNV | 0.4 | ClairS-TO+LP (hap) | 0.756 | 0.507 | 0.607 |
| HCC1395_NYGC | SNV | 0.4 | ClairS-TO+LP (hap+5mC) | 0.854 | 0.497 | 0.628 |
| HCC1395_NYGC | SNV | 0.4 | DeepSomatic-TO | 0.484 | 0.689 | 0.568 |
| HCC1395_NYGC | SNV | 0.4 | DeepSomatic-TO+LP (hap) | 0.794 | 0.539 | 0.643 |
| HCC1395_NYGC | SNV | 0.4 | DeepSomatic-TO+LP (hap+5mC) | 0.852 | 0.530 | 0.653 |
| HCC1395_NYGC | SNV | 0.2 | ClairS-TO | 0.324 | 0.234 | 0.272 |
| HCC1395_NYGC | SNV | 0.2 | ClairS-TO+LP (hap) | 0.610 | 0.231 | 0.335 |
| HCC1395_NYGC | SNV | 0.2 | ClairS-TO+LP (hap+5mC) | 0.804 | 0.229 | 0.356 |
| HCC1395_NYGC | SNV | 0.2 | DeepSomatic-TO | 0.362 | 0.437 | 0.396 |
| HCC1395_NYGC | SNV | 0.2 | DeepSomatic-TO+LP (hap) | 0.694 | 0.325 | 0.443 |
| HCC1395_NYGC | SNV | 0.2 | DeepSomatic-TO+LP (hap+5mC) | 0.799 | 0.322 | 0.459 |
| HCC1937_UCSC | SNV | 1 | ClairS-TO | 0.483 | 0.748 | 0.587 |
| HCC1937_UCSC | SNV | 1 | ClairS-TO+LP (hap) | 0.462 | 0.774 | 0.579 |
| HCC1937_UCSC | SNV | 1 | ClairS-TO+LP (hap+5mC) | 0.462 | 0.774 | 0.579 |
| HCC1937_UCSC | SNV | 1 | DeepSomatic-TO | 0.379 | 0.834 | 0.521 |
| HCC1937_UCSC | SNV | 1 | DeepSomatic-TO+LP (hap) | 0.437 | 0.816 | 0.569 |
| HCC1937_UCSC | SNV | 1 | DeepSomatic-TO+LP (hap+5mC) | 0.437 | 0.816 | 0.569 |
| HCC1937_UCSC | SNV | 0.8 | ClairS-TO | 0.409 | 0.692 | 0.514 |
| HCC1937_UCSC | SNV | 0.8 | ClairS-TO+LP (hap) | 0.685 | 0.687 | 0.686 |
| HCC1937_UCSC | SNV | 0.8 | ClairS-TO+LP (hap+5mC) | 0.686 | 0.682 | 0.684 |
| HCC1937_UCSC | SNV | 0.8 | DeepSomatic-TO | 0.312 | 0.791 | 0.448 |
| HCC1937_UCSC | SNV | 0.8 | DeepSomatic-TO+LP (hap) | 0.636 | 0.717 | 0.674 |
| HCC1937_UCSC | SNV | 0.8 | DeepSomatic-TO+LP (hap+5mC) | 0.636 | 0.715 | 0.673 |
| HCC1937_UCSC | SNV | 0.6 | ClairS-TO | 0.338 | 0.608 | 0.434 |
| HCC1937_UCSC | SNV | 0.6 | ClairS-TO+LP (hap) | 0.620 | 0.635 | 0.627 |
| HCC1937_UCSC | SNV | 0.6 | ClairS-TO+LP (hap+5mC) | 0.727 | 0.589 | 0.651 |
| HCC1937_UCSC | SNV | 0.6 | DeepSomatic-TO | 0.275 | 0.724 | 0.399 |
| HCC1937_UCSC | SNV | 0.6 | DeepSomatic-TO+LP (hap) | 0.604 | 0.685 | 0.642 |
| HCC1937_UCSC | SNV | 0.6 | DeepSomatic-TO+LP (hap+5mC) | 0.658 | 0.640 | 0.649 |
| HCC1937_UCSC | SNV | 0.4 | ClairS-TO | 0.268 | 0.454 | 0.337 |
| HCC1937_UCSC | SNV | 0.4 | ClairS-TO+LP (hap) | 0.538 | 0.487 | 0.511 |
| HCC1937_UCSC | SNV | 0.4 | ClairS-TO+LP (hap+5mC) | 0.706 | 0.471 | 0.565 |
| HCC1937_UCSC | SNV | 0.4 | DeepSomatic-TO | 0.228 | 0.589 | 0.329 |

| Sample | Type | Fraction | Variant caller | Precision | Recall | F1-score |
|---|---|---|---|---|---|---|
| HCC1937_UCSC | SNV | 0.4 | DeepSomatic-TO+LP (hap) | 0.529 | 0.552 | 0.540 |
| HCC1937_UCSC | SNV | 0.4 | DeepSomatic-TO+LP (hap+5mC) | 0.623 | 0.536 | 0.576 |
| HCC1937_UCSC | SNV | 0.2 | ClairS-TO | 0.128 | 0.184 | 0.151 |
| HCC1937_UCSC | SNV | 0.2 | ClairS-TO+LP (hap) | 0.348 | 0.220 | 0.270 |
| HCC1937_UCSC | SNV | 0.2 | ClairS-TO+LP (hap+5mC) | 0.619 | 0.213 | 0.317 |
| HCC1937_UCSC | SNV | 0.2 | DeepSomatic-TO | 0.131 | 0.309 | 0.184 |
| HCC1937_UCSC | SNV | 0.2 | DeepSomatic-TO+LP (hap) | 0.360 | 0.276 | 0.312 |
| HCC1937_UCSC | SNV | 0.2 | DeepSomatic-TO+LP (hap+5mC) | 0.502 | 0.267 | 0.349 |
| HCC1954_UCSC | SNV | 1 | ClairS-TO | 0.248 | 0.799 | 0.379 |
| HCC1954_UCSC | SNV | 1 | ClairS-TO+LP (hap) | 0.254 | 0.834 | 0.389 |
| HCC1954_UCSC | SNV | 1 | ClairS-TO+LP (hap+5mC) | 0.248 | 0.839 | 0.382 |
| HCC1954_UCSC | SNV | 1 | DeepSomatic-TO | 0.214 | 0.888 | 0.345 |
| HCC1954_UCSC | SNV | 1 | DeepSomatic-TO+LP (hap) | 0.260 | 0.872 | 0.401 |
| HCC1954_UCSC | SNV | 1 | DeepSomatic-TO+LP (hap+5mC) | 0.253 | 0.877 | 0.393 |
| HCC1954_UCSC | SNV | 0.8 | ClairS-TO | 0.218 | 0.718 | 0.334 |
| HCC1954_UCSC | SNV | 0.8 | ClairS-TO+LP (hap) | 0.521 | 0.733 | 0.609 |
| HCC1954_UCSC | SNV | 0.8 | ClairS-TO+LP (hap+5mC) | 0.521 | 0.731 | 0.609 |
| HCC1954_UCSC | SNV | 0.8 | DeepSomatic-TO | 0.196 | 0.854 | 0.319 |
| HCC1954_UCSC | SNV | 0.8 | DeepSomatic-TO+LP (hap) | 0.616 | 0.774 | 0.686 |
| HCC1954_UCSC | SNV | 0.8 | DeepSomatic-TO+LP (hap+5mC) | 0.616 | 0.773 | 0.686 |
| HCC1954_UCSC | SNV | 0.6 | ClairS-TO | 0.175 | 0.577 | 0.269 |
| HCC1954_UCSC | SNV | 0.6 | ClairS-TO+LP (hap) | 0.448 | 0.617 | 0.519 |
| HCC1954_UCSC | SNV | 0.6 | ClairS-TO+LP (hap+5mC) | 0.568 | 0.608 | 0.587 |
| HCC1954_UCSC | SNV | 0.6 | DeepSomatic-TO | 0.179 | 0.782 | 0.292 |
| HCC1954_UCSC | SNV | 0.6 | DeepSomatic-TO+LP (hap) | 0.579 | 0.723 | 0.643 |
| HCC1954_UCSC | SNV | 0.6 | DeepSomatic-TO+LP (hap+5mC) | 0.661 | 0.713 | 0.686 |
| HCC1954_UCSC | SNV | 0.4 | ClairS-TO | 0.117 | 0.364 | 0.177 |
| HCC1954_UCSC | SNV | 0.4 | ClairS-TO+LP (hap) | 0.340 | 0.406 | 0.370 |
| HCC1954_UCSC | SNV | 0.4 | ClairS-TO+LP (hap+5mC) | 0.467 | 0.404 | 0.433 |
| HCC1954_UCSC | SNV | 0.4 | DeepSomatic-TO | 0.147 | 0.619 | 0.237 |
| HCC1954_UCSC | SNV | 0.4 | DeepSomatic-TO+LP (hap) | 0.514 | 0.565 | 0.538 |
| HCC1954_UCSC | SNV | 0.4 | DeepSomatic-TO+LP (hap+5mC) | 0.614 | 0.562 | 0.587 |
| HCC1954_UCSC | SNV | 0.2 | ClairS-TO | 0.048 | 0.140 | 0.072 |
| HCC1954_UCSC | SNV | 0.2 | ClairS-TO+LP (hap) | 0.180 | 0.168 | 0.174 |
| HCC1954_UCSC | SNV | 0.2 | ClairS-TO+LP (hap+5mC) | 0.303 | 0.167 | 0.216 |
| HCC1954_UCSC | SNV | 0.2 | DeepSomatic-TO | 0.087 | 0.347 | 0.139 |
| HCC1954_UCSC | SNV | 0.2 | DeepSomatic-TO+LP (hap) | 0.358 | 0.309 | 0.332 |
| HCC1954_UCSC | SNV | 0.2 | DeepSomatic-TO+LP (hap+5mC) | 0.482 | 0.308 | 0.376 |
| H1437_UCSC | Indel | 1 | ClairS-TO | 0.569 | 0.279 | 0.375 |
| H1437_UCSC | Indel | 1 | ClairS-TO+LP (hap) | 0.511 | 0.343 | 0.410 |
| H1437_UCSC | Indel | 1 | ClairS-TO+LP (hap+5mC) | 0.509 | 0.306 | 0.382 |
| H1437_UCSC | Indel | 1 | DeepSomatic-TO | 0.304 | 0.110 | 0.162 |
| H1437_UCSC | Indel | 1 | DeepSomatic-TO+LP (hap) | 0.443 | 0.106 | 0.171 |
| H1437_UCSC | Indel | 1 | DeepSomatic-TO+LP (hap+5mC) | 0.429 | 0.098 | 0.159 |
| H1437_UCSC | Indel | 0.8 | ClairS-TO | 0.527 | 0.233 | 0.323 |
| H1437_UCSC | Indel | 0.8 | ClairS-TO+LP (hap) | 0.628 | 0.217 | 0.323 |
| H1437_UCSC | Indel | 0.8 | ClairS-TO+LP (hap+5mC) | 0.628 | 0.217 | 0.323 |
| H1437_UCSC | Indel | 0.8 | DeepSomatic-TO | 0.319 | 0.135 | 0.190 |
| H1437_UCSC | Indel | 0.8 | DeepSomatic-TO+LP (hap) | 0.635 | 0.096 | 0.167 |
| H1437_UCSC | Indel | 0.8 | DeepSomatic-TO+LP (hap+5mC) | 0.635 | 0.096 | 0.167 |
| H1437_UCSC | Indel | 0.6 | ClairS-TO | 0.504 | 0.170 | 0.254 |
| H1437_UCSC | Indel | 0.6 | ClairS-TO+LP (hap) | 0.601 | 0.203 | 0.304 |
| H1437_UCSC | Indel | 0.6 | ClairS-TO+LP (hap+5mC) | 0.752 | 0.171 | 0.279 |
| H1437_UCSC | Indel | 0.6 | DeepSomatic-TO | 0.288 | 0.132 | 0.181 |
| H1437_UCSC | Indel | 0.6 | DeepSomatic-TO+LP (hap) | 0.623 | 0.116 | 0.196 |
| H1437_UCSC | Indel | 0.6 | DeepSomatic-TO+LP (hap+5mC) | 0.683 | 0.105 | 0.182 |
| H1437_UCSC | Indel | 0.4 | ClairS-TO | 0.307 | 0.073 | 0.117 |
| H1437_UCSC | Indel | 0.4 | ClairS-TO+LP (hap) | 0.369 | 0.105 | 0.163 |
| H1437_UCSC | Indel | 0.4 | ClairS-TO+LP (hap+5mC) | 0.540 | 0.098 | 0.166 |
| H1437_UCSC | Indel | 0.4 | DeepSomatic-TO | 0.218 | 0.095 | 0.132 |
| H1437_UCSC | Indel | 0.4 | DeepSomatic-TO+LP (hap) | 0.552 | 0.088 | 0.152 |
| H1437_UCSC | Indel | 0.4 | DeepSomatic-TO+LP (hap+5mC) | 0.633 | 0.084 | 0.149 |
| H1437_UCSC | Indel | 0.2 | ClairS-TO | 0.146 | 0.008 | 0.014 |
| H1437_UCSC | Indel | 0.2 | ClairS-TO+LP (hap) | 0.053 | 0.012 | 0.019 |
| H1437_UCSC | Indel | 0.2 | ClairS-TO+LP (hap+5mC) | 0.116 | 0.011 | 0.021 |
| H1437_UCSC | Indel | 0.2 | DeepSomatic-TO | 0.061 | 0.023 | 0.033 |
| H1437_UCSC | Indel | 0.2 | DeepSomatic-TO+LP (hap) | 0.230 | 0.022 | 0.039 |
| H1437_UCSC | Indel | 0.2 | DeepSomatic-TO+LP (hap+5mC) | 0.348 | 0.021 | 0.040 |
| H2009_UCSC | Indel | 1 | ClairS-TO | 0.764 | 0.570 | 0.653 |
| H2009_UCSC | Indel | 1 | ClairS-TO+LP (hap) | 0.721 | 0.672 | 0.696 |
| H2009_UCSC | Indel | 1 | ClairS-TO+LP (hap+5mC) | 0.720 | 0.619 | 0.666 |
| H2009_UCSC | Indel | 1 | DeepSomatic-TO | 0.477 | 0.185 | 0.267 |
| H2009_UCSC | Indel | 1 | DeepSomatic-TO+LP (hap) | 0.630 | 0.180 | 0.279 |
| H2009_UCSC | Indel | 1 | DeepSomatic-TO+LP (hap+5mC) | 0.626 | 0.171 | 0.269 |
| H2009_UCSC | Indel | 0.8 | ClairS-TO | 0.721 | 0.467 | 0.567 |
| H2009_UCSC | Indel | 0.8 | ClairS-TO+LP (hap) | 0.806 | 0.429 | 0.560 |
| H2009_UCSC | Indel | 0.8 | ClairS-TO+LP (hap+5mC) | 0.806 | 0.429 | 0.560 |
| H2009_UCSC | Indel | 0.8 | DeepSomatic-TO | 0.500 | 0.227 | 0.312 |
| H2009_UCSC | Indel | 0.8 | DeepSomatic-TO+LP (hap) | 0.777 | 0.173 | 0.283 |
| H2009_UCSC | Indel | 0.8 | DeepSomatic-TO+LP (hap+5mC) | 0.777 | 0.173 | 0.283 |
| H2009_UCSC | Indel | 0.6 | ClairS-TO | 0.613 | 0.319 | 0.419 |
| H2009_UCSC | Indel | 0.6 | ClairS-TO+LP (hap) | 0.753 | 0.362 | 0.489 |
| H2009_UCSC | Indel | 0.6 | ClairS-TO+LP (hap+5mC) | 0.874 | 0.326 | 0.475 |
| H2009_UCSC | Indel | 0.6 | DeepSomatic-TO | 0.454 | 0.218 | 0.295 |
| H2009_UCSC | Indel | 0.6 | DeepSomatic-TO+LP (hap) | 0.765 | 0.191 | 0.306 |
| H2009_UCSC | Indel | 0.6 | DeepSomatic-TO+LP (hap+5mC) | 0.824 | 0.177 | 0.292 |
| H2009_UCSC | Indel | 0.4 | ClairS-TO | 0.400 | 0.152 | 0.221 |
| H2009_UCSC | Indel | 0.4 | ClairS-TO+LP (hap) | 0.571 | 0.197 | 0.293 |
| H2009_UCSC | Indel | 0.4 | ClairS-TO+LP (hap+5mC) | 0.734 | 0.190 | 0.302 |
| H2009_UCSC | Indel | 0.4 | DeepSomatic-TO | 0.356 | 0.160 | 0.221 |

| Sample | Type | Fraction | Variant caller | Precision | Recall | F1-score |
|---|---|---|---|---|---|---|
| H2009_UCSC | Indel | 0.4 | DeepSomatic-TO+LP (hap) | 0.681 | 0.148 | 0.244 |
| H2009_UCSC | Indel | 0.4 | DeepSomatic-TO+LP (hap+5mC) | 0.762 | 0.144 | 0.242 |
| H2009_UCSC | Indel | 0.2 | ClairS-TO | 0.053 | 0.015 | 0.023 |
| H2009_UCSC | Indel | 0.2 | ClairS-TO+LP (hap) | 0.130 | 0.024 | 0.041 |
| H2009_UCSC | Indel | 0.2 | ClairS-TO+LP (hap+5mC) | 0.280 | 0.023 | 0.043 |
| H2009_UCSC | Indel | 0.2 | DeepSomatic-TO | 0.092 | 0.034 | 0.050 |
| H2009_UCSC | Indel | 0.2 | DeepSomatic-TO+LP (hap) | 0.311 | 0.031 | 0.056 |
| H2009_UCSC | Indel | 0.2 | DeepSomatic-TO+LP (hap+5mC) | 0.483 | 0.030 | 0.057 |
| HCC1395_NYGC | Indel | 1 | ClairS-TO | 0.434 | 0.401 | 0.417 |
| HCC1395_NYGC | Indel | 1 | ClairS-TO+LP (hap) | 0.342 | 0.488 | 0.403 |
| HCC1395_NYGC | Indel | 1 | ClairS-TO+LP (hap+5mC) | 0.350 | 0.479 | 0.404 |
| HCC1395_NYGC | Indel | 1 | DeepSomatic-TO | 0.464 | 0.630 | 0.534 |
| HCC1395_NYGC | Indel | 1 | DeepSomatic-TO+LP (hap) | 0.587 | 0.598 | 0.593 |
| HCC1395_NYGC | Indel | 1 | DeepSomatic-TO+LP (hap+5mC) | 0.585 | 0.586 | 0.586 |
| HCC1395_NYGC | Indel | 0.8 | ClairS-TO | 0.403 | 0.328 | 0.362 |
| HCC1395_NYGC | Indel | 0.8 | ClairS-TO+LP (hap) | 0.467 | 0.344 | 0.396 |
| HCC1395_NYGC | Indel | 0.8 | ClairS-TO+LP (hap+5mC) | 0.467 | 0.344 | 0.396 |
| HCC1395_NYGC | Indel | 0.8 | DeepSomatic-TO | 0.426 | 0.578 | 0.491 |
| HCC1395_NYGC | Indel | 0.8 | DeepSomatic-TO+LP (hap) | 0.685 | 0.438 | 0.534 |
| HCC1395_NYGC | Indel | 0.8 | DeepSomatic-TO+LP (hap+5mC) | 0.685 | 0.438 | 0.534 |
| HCC1395_NYGC | Indel | 0.6 | ClairS-TO | 0.299 | 0.220 | 0.253 |
| HCC1395_NYGC | Indel | 0.6 | ClairS-TO+LP (hap) | 0.428 | 0.255 | 0.320 |
| HCC1395_NYGC | Indel | 0.6 | ClairS-TO+LP (hap+5mC) | 0.676 | 0.230 | 0.343 |
| HCC1395_NYGC | Indel | 0.6 | DeepSomatic-TO | 0.348 | 0.496 | 0.409 |
| HCC1395_NYGC | Indel | 0.6 | DeepSomatic-TO+LP (hap) | 0.631 | 0.391 | 0.483 |
| HCC1395_NYGC | Indel | 0.6 | DeepSomatic-TO+LP (hap+5mC) | 0.697 | 0.361 | 0.476 |
| HCC1395_NYGC | Indel | 0.4 | ClairS-TO | 0.134 | 0.093 | 0.110 |
| HCC1395_NYGC | Indel | 0.4 | ClairS-TO+LP (hap) | 0.251 | 0.120 | 0.162 |
| HCC1395_NYGC | Indel | 0.4 | ClairS-TO+LP (hap+5mC) | 0.484 | 0.115 | 0.186 |
| HCC1395_NYGC | Indel | 0.4 | DeepSomatic-TO | 0.258 | 0.341 | 0.294 |
| HCC1395_NYGC | Indel | 0.4 | DeepSomatic-TO+LP (hap) | 0.572 | 0.267 | 0.364 |
| HCC1395_NYGC | Indel | 0.4 | DeepSomatic-TO+LP (hap+5mC) | 0.659 | 0.254 | 0.367 |
| HCC1395_NYGC | Indel | 0.2 | ClairS-TO | 0.011 | 0.007 | 0.009 |
| HCC1395_NYGC | Indel | 0.2 | ClairS-TO+LP (hap) | 0.035 | 0.015 | 0.021 |
| HCC1395_NYGC | Indel | 0.2 | ClairS-TO+LP (hap+5mC) | 0.120 | 0.012 | 0.021 |
| HCC1395_NYGC | Indel | 0.2 | DeepSomatic-TO | 0.092 | 0.103 | 0.097 |
| HCC1395_NYGC | Indel | 0.2 | DeepSomatic-TO+LP (hap) | 0.315 | 0.078 | 0.125 |
| HCC1395_NYGC | Indel | 0.2 | DeepSomatic-TO+LP (hap+5mC) | 0.531 | 0.069 | 0.122 |
| HCC1937_UCSC | Indel | 1 | ClairS-TO | 0.272 | 0.413 | 0.328 |
| HCC1937_UCSC | Indel | 1 | ClairS-TO+LP (hap) | 0.203 | 0.535 | 0.294 |
| HCC1937_UCSC | Indel | 1 | ClairS-TO+LP (hap+5mC) | 0.196 | 0.464 | 0.275 |
| HCC1937_UCSC | Indel | 1 | DeepSomatic-TO | 0.219 | 0.378 | 0.277 |
| HCC1937_UCSC | Indel | 1 | DeepSomatic-TO+LP (hap) | 0.297 | 0.363 | 0.327 |
| HCC1937_UCSC | Indel | 1 | DeepSomatic-TO+LP (hap+5mC) | 0.293 | 0.347 | 0.317 |
| HCC1937_UCSC | Indel | 0.8 | ClairS-TO | 0.285 | 0.391 | 0.330 |
| HCC1937_UCSC | Indel | 0.8 | ClairS-TO+LP (hap) | 0.394 | 0.401 | 0.397 |
| HCC1937_UCSC | Indel | 0.8 | ClairS-TO+LP (hap+5mC) | 0.394 | 0.401 | 0.397 |
| HCC1937_UCSC | Indel | 0.8 | DeepSomatic-TO | 0.246 | 0.432 | 0.313 |
| HCC1937_UCSC | Indel | 0.8 | DeepSomatic-TO+LP (hap) | 0.457 | 0.337 | 0.388 |
| HCC1937_UCSC | Indel | 0.8 | DeepSomatic-TO+LP (hap+5mC) | 0.457 | 0.337 | 0.388 |
| HCC1937_UCSC | Indel | 0.6 | ClairS-TO | 0.240 | 0.320 | 0.275 |
| HCC1937_UCSC | Indel | 0.6 | ClairS-TO+LP (hap) | 0.424 | 0.368 | 0.394 |
| HCC1937_UCSC | Indel | 0.6 | ClairS-TO+LP (hap+5mC) | 0.706 | 0.296 | 0.417 |
| HCC1937_UCSC | Indel | 0.6 | DeepSomatic-TO | 0.224 | 0.417 | 0.292 |
| HCC1937_UCSC | Indel | 0.6 | DeepSomatic-TO+LP (hap) | 0.524 | 0.384 | 0.443 |
| HCC1937_UCSC | Indel | 0.6 | DeepSomatic-TO+LP (hap+5mC) | 0.657 | 0.334 | 0.443 |
| HCC1937_UCSC | Indel | 0.4 | ClairS-TO | 0.117 | 0.160 | 0.135 |
| HCC1937_UCSC | Indel | 0.4 | ClairS-TO+LP (hap) | 0.247 | 0.207 | 0.225 |
| HCC1937_UCSC | Indel | 0.4 | ClairS-TO+LP (hap+5mC) | 0.567 | 0.187 | 0.281 |
| HCC1937_UCSC | Indel | 0.4 | DeepSomatic-TO | 0.162 | 0.309 | 0.212 |
| HCC1937_UCSC | Indel | 0.4 | DeepSomatic-TO+LP (hap) | 0.458 | 0.285 | 0.351 |
| HCC1937_UCSC | Indel | 0.4 | DeepSomatic-TO+LP (hap+5mC) | 0.621 | 0.266 | 0.373 |
| HCC1937_UCSC | Indel | 0.2 | ClairS-TO | 0.017 | 0.024 | 0.020 |
| HCC1937_UCSC | Indel | 0.2 | ClairS-TO+LP (hap) | 0.054 | 0.047 | 0.050 |
| HCC1937_UCSC | Indel | 0.2 | ClairS-TO+LP (hap+5mC) | 0.28 | 0.041 | 0.072 |
| HCC1937_UCSC | Indel | 0.2 | DeepSomatic-TO | 0.063 | 0.113 | 0.081 |
| HCC1937_UCSC | Indel | 0.2 | DeepSomatic-TO+LP (hap) | 0.262 | 0.095 | 0.139 |
| HCC1937_UCSC | Indel | 0.2 | DeepSomatic-TO+LP (hap+5mC) | 0.535 | 0.083 | 0.144 |
| HCC1954_UCSC | Indel | 1 | ClairS-TO | 0.165 | 0.212 | 0.186 |
| HCC1954_UCSC | Indel | 1 | ClairS-TO+LP (hap) | 0.169 | 0.292 | 0.214 |
| HCC1954_UCSC | Indel | 1 | ClairS-TO+LP (hap+5mC) | 0.169 | 0.292 | 0.214 |
| HCC1954_UCSC | Indel | 1 | DeepSomatic-TO | 0.145 | 0.135 | 0.140 |
| HCC1954_UCSC | Indel | 1 | DeepSomatic-TO+LP (hap) | 0.216 | 0.130 | 0.162 |
| HCC1954_UCSC | Indel | 1 | DeepSomatic-TO+LP (hap+5mC) | 0.216 | 0.130 | 0.162 |
| HCC1954_UCSC | Indel | 0.8 | ClairS-TO | 0.100 | 0.128 | 0.112 |
| HCC1954_UCSC | Indel | 0.8 | ClairS-TO+LP (hap) | 0.249 | 0.171 | 0.203 |
| HCC1954_UCSC | Indel | 0.8 | ClairS-TO+LP (hap+5mC) | 0.249 | 0.171 | 0.203 |
| HCC1954_UCSC | Indel | 0.8 | DeepSomatic-TO | 0.113 | 0.104 | 0.109 |
| HCC1954_UCSC | Indel | 0.8 | DeepSomatic-TO+LP (hap) | 0.389 | 0.090 | 0.146 |
| HCC1954_UCSC | Indel | 0.8 | DeepSomatic-TO+LP (hap+5mC) | 0.389 | 0.090 | 0.146 |
| HCC1954_UCSC | Indel | 0.6 | ClairS-TO | 0.052 | 0.067 | 0.058 |
| HCC1954_UCSC | Indel | 0.6 | ClairS-TO+LP (hap) | 0.134 | 0.090 | 0.108 |
| HCC1954_UCSC | Indel | 0.6 | ClairS-TO+LP (hap+5mC) | 0.304 | 0.079 | 0.126 |
| HCC1954_UCSC | Indel | 0.6 | DeepSomatic-TO | 0.080 | 0.075 | 0.077 |
| HCC1954_UCSC | Indel | 0.6 | DeepSomatic-TO+LP (hap) | 0.308 | 0.067 | 0.110 |
| HCC1954_UCSC | Indel | 0.6 | DeepSomatic-TO+LP (hap+5mC) | 0.458 | 0.062 | 0.109 |
| HCC1954_UCSC | Indel | 0.4 | ClairS-TO | 0.014 | 0.019 | 0.016 |
| HCC1954_UCSC | Indel | 0.4 | ClairS-TO+LP (hap) | 0.040 | 0.028 | 0.033 |
| HCC1954_UCSC | Indel | 0.4 | ClairS-TO+LP (hap+5mC) | 0.114 | 0.027 | 0.044 |
| HCC1954_UCSC | Indel | 0.4 | DeepSomatic-TO | 0.040 | 0.037 | 0.038 |

| Sample | Type | Fraction | Variant caller | Precision | Recall | F1-score |
|---|---|---|---|---|---|---|
| HCC1954_UCSC | Indel | 0.4 | DeepSomatic-TO+LP (hap) | 0.167 | 0.033 | 0.055 |
| HCC1954_UCSC | Indel | 0.4 | DeepSomatic-TO+LP (hap+5mC) | 0.310 | 0.033 | 0.059 |
| HCC1954_UCSC | Indel | 0.2 | ClairS-TO | 0.002 | 0.003 | 0.002 |
| HCC1954_UCSC | Indel | 0.2 | ClairS-TO+LP (hap) | 0.005 | 0.004 | 0.004 |
| HCC1954_UCSC | Indel | 0.2 | ClairS-TO+LP (hap+5mC) | 0.019 | 0.004 | 0.006 |
| HCC1954_UCSC | Indel | 0.2 | DeepSomatic-TO | 0.009 | 0.009 | 0.009 |
| HCC1954_UCSC | Indel | 0.2 | DeepSomatic-TO+LP (hap) | 0.045 | 0.008 | 0.014 |
| HCC1954_UCSC | Indel | 0.2 | DeepSomatic-TO+LP (hap+5mC) | 0.109 | 0.008 | 0.016 |

**Supplementary Table 7**: Inputs and modeling assumptions of the estimators compared in this study. The quantity in the final column differs between approaches: ASCAT and PURPLE infer cellular purity jointly with ploidy from allele-specific copy-number profiles, whereas LongPhase-TO infers the fraction of sequenced molecules of tumor origin from germline haplotype imbalance. The two quantities coincide in near-diploid genomes but diverge under whole-genome duplication. For all comparisons, each caller's purity was therefore converted to DNA fraction using its own fitted ploidy (see Methods).

| Method | Matched normal | Copy-number segmentation | Ploidy fit | Quantity estimated |
|---|---|---|---|---|
| ASCAT (T+N) | Required | Required | Required | Ploidy-adjusted cellular purity |
| ASCAT (TO) | No | Required | Required | Ploidy-adjusted cellular purity |
| PURPLE (TO) | No | Required | Required | Ploidy-adjusted cellular purity |
| LongPhase-TO | No | No | No | Read-level tumor DNA fraction |

**Supplementary Table 8**: Per-mixture tumor DNA fraction estimates for the nanopore panel, comparing ASCAT, PURPLE and LongPhase-TO across eight cell-line datasets and five DNA-fraction levels (40 mixtures). ASCAT and PURPLE values are cellular purities converted to DNA fraction with each tool's fitted ploidy (see Methods). LP-TO (C) and LP-TO (D) denote LongPhase-TO applied to somatic variants from ClairS-TO and DeepSomatic-TO, respectively. The expected value is the achieved read-level mixing fraction, which differs slightly from the nominal target where coverage was limiting. Entries marked n.a. indicate that the method returned no estimate.

| Dataset | Expected | ASCAT (T+N) | ASCAT (TO) | PURPLE (TO) | LP-TO (C) | LP-TO (D) |
|---|---|---|---|---|---|---|
| COLO829_ONT | 0.200 | 0.540 | 1.000 | 0.212 | 0.203 | 0.197 |
| COLO829_ONT | 0.400 | 0.420 | 0.851 | 0.410 | 0.396 | 0.395 |
| COLO829_ONT | 0.600 | 0.628 | 0.660 | 0.608 | 0.684 | 0.657 |
| COLO829_ONT | 0.805 | 0.820 | 0.835 | 0.794 | 0.830 | 0.800 |
| COLO829_ONT | 1.000 | 0.994 | 0.892 | 0.433 | 1.000 | 0.992 |
| COLO829_NYGC | 0.200 | 1.000 | 1.000 | 0.234 | 0.155 | 0.190 |
| COLO829_NYGC | 0.400 | 0.446 | 0.973 | 0.414 | 0.394 | 0.414 |
| COLO829_NYGC | 0.600 | 0.662 | 0.690 | 0.612 | 0.700 | 0.663 |
| COLO829_NYGC | 0.800 | 0.851 | 0.868 | 0.804 | 0.838 | 0.797 |
| COLO829_NYGC | 1.000 | 0.994 | 0.976 | 0.419 | 0.999 | 0.996 |
| HCC1395_HKU | 0.200 | 1.000 | 1.000 | 0.348 | 0.274 | 0.195 |
| HCC1395_HKU | 0.400 | 0.459 | 0.972 | 0.399 | 0.361 | 0.406 |
| HCC1395_HKU | 0.600 | 0.652 | 0.670 | 0.596 | 0.584 | 0.615 |
| HCC1395_HKU | 0.800 | 0.822 | 0.838 | 0.788 | 0.785 | 0.742 |
| HCC1395_HKU | 1.000 | 0.993 | 0.942 | 0.932 | 0.977 | 0.993 |
| HCC1395_NYGC | 0.194 | 1.000 | 0.904 | 0.370 | 0.221 | 0.222 |
| HCC1395_NYGC | 0.396 | 0.468 | 1.000 | 0.397 | 0.335 | 0.367 |
| HCC1395_NYGC | 0.600 | 0.661 | 0.679 | 0.598 | 0.572 | 0.549 |
| HCC1395_NYGC | 0.800 | 0.834 | 0.850 | 0.788 | 0.735 | 0.704 |
| HCC1395_NYGC | 1.000 | 0.989 | 0.939 | 0.959 | 0.993 | 1.000 |
| H1437_UCSC | 0.200 | 1.000 | 1.000 | 0.238 | 0.244 | 0.181 |
| H1437_UCSC | 0.400 | 0.465 | n.a. | 0.691 | 0.401 | 0.367 |
| H1437_UCSC | 0.600 | 0.661 | 0.708 | 0.606 | 0.625 | 0.594 |
| H1437_UCSC | 0.800 | 0.839 | 0.857 | 0.802 | 0.800 | 0.757 |
| H1437_UCSC | 1.000 | 1.000 | 0.956 | 0.994 | 1.000 | 1.000 |
| H2009_UCSC | 0.210 | 0.247 | 0.977 | 0.237 | 0.260 | 0.241 |
| H2009_UCSC | 0.400 | 0.443 | 0.934 | 0.407 | 0.385 | 0.362 |
| H2009_UCSC | 0.600 | 0.640 | 0.706 | 0.600 | 0.609 | 0.595 |

| Dataset | Expected | ASCAT (T+N) | ASCAT (TO) | PURPLE (TO) | LP-TO (C) | LP-TO (D) |
|---|---|---|---|---|---|---|
| H2009_UCSC | 0.800 | 0.826 | 0.874 | 0.792 | 0.806 | 0.767 |
| H2009_UCSC | 1.000 | 0.992 | 0.982 | 0.925 | 1.000 | 1.000 |
| HCC1937_UCSC | 0.188 | 1.000 | 1.000 | 0.399 | 0.288 | 0.319 |
| HCC1937_UCSC | 0.395 | 1.000 | 1.000 | 0.604 | 0.350 | 0.401 |
| HCC1937_UCSC | 0.600 | 0.575 | n.a. | 0.623 | 0.534 | 0.584 |
| HCC1937_UCSC | 0.800 | 0.784 | 0.795 | 0.799 | 0.745 | 0.748 |
| HCC1937_UCSC | 1.000 | 0.996 | 0.920 | 0.931 | 0.954 | 0.977 |
| HCC1954_UCSC | 0.200 | 1.000 | 1.000 | 0.318 | 0.840 | 0.373 |
| HCC1954_UCSC | 0.400 | 1.000 | 1.000 | 0.405 | 0.508 | 0.351 |
| HCC1954_UCSC | 0.600 | n.a. | n.a. | 0.600 | 0.491 | 0.472 |
| HCC1954_UCSC | 0.800 | n.a. | n.a. | 0.786 | 0.611 | 0.579 |
| HCC1954_UCSC | 1.000 | n.a. | 0.749 | 0.567 | 0.798 | 0.702 |

**Supplementary Table 9**: Accuracy of tumor DNA fraction estimation on both sequencing platforms. The unit of analysis is the cell-line dataset, not the individual mixture: the five DNA-fraction levels within a dataset are downsampled from one library and are therefore not independent. MAE is the mean absolute error against the achieved mixing fraction. The jackknife range is the smallest and largest overall MAE obtained when any single cell-line dataset is excluded. CCC is Lin's concordance correlation coefficient, which penalizes both bias and scale shift and is therefore preferred here over Pearson correlation. Slope and intercept are from ordinary least squares of the estimate on the achieved fraction. The two rightmost count columns give the number of mixtures whose estimate fell within 0.05 and within 0.10 of the achieved fraction; denominators exclude mixtures for which the method returned no estimate, counted in the final column.

| Method | MAE | Jackknife | CCC | Slope | Intercept | ≤0.05 | ≤0.10 | No est. |
|---|---|---|---|---|---|---|---|---|
| *Nanopore panel (8 cell-line datasets, 40 mixtures)* | | | | | | | | |
| ASCAT (T+N) | 0.20 | [0.17, 0.22] | 0.24 | 0.25 | +0.63 | 20/37 | 28/37 | 3 |
| ASCAT (TO) | 0.33 | [0.30, 0.33] | −0.11 | −0.12 | +0.96 | 7/36 | 17/36 | 4 |
| PURPLE (TO) | 0.08 | [0.07, 0.09] | 0.79 | 0.64 | +0.20 | 28/40 | 31/40 | 0 |
| LongPhase-TO (ClairS-TO) | 0.06 | [0.03, 0.07] | 0.90 | 0.84 | +0.10 | 27/40 | 34/40 | 0 |
| LongPhase-TO (DeepSomatic-TO) | 0.04 | [0.03, 0.05] | 0.96 | 0.89 | +0.05 | 29/40 | 35/40 | 0 |
| *PacBio HiFi panel (6 cell-line datasets, 30 mixtures)* | | | | | | | | |
| ASCAT (T+N) | 0.09 | [0.07, 0.10] | 0.59 | 0.58 | +0.33 | 27/30 | 27/30 | 0 |
| ASCAT (TO) | 0.24 | [0.22, 0.25] | −0.04 | −0.04 | +0.81 | 16/29 | 19/29 | 1 |
| PURPLE (TO) | 0.07 | [0.06, 0.08] | 0.87 | 0.71 | +0.17 | 21/30 | 23/30 | 0 |
| LongPhase-TO (ClairS-TO) | 0.08 | [0.04, 0.09] | 0.89 | 0.93 | 0.00 | 18/30 | 24/30 | 0 |
| LongPhase-TO (DeepSomatic-TO) | 0.06 | [0.03, 0.06] | 0.94 | 0.93 | 0.00 | 21/30 | 25/30 | 0 |

**Supplementary Table 10**: Mean absolute error of tumor DNA fraction estimation at each designed dilution level. Stratifying by level separates the regimes in which each approach degrades: the copy-number-based estimators lose accuracy at low tumor content, PURPLE loses accuracy in near-pure samples, and LongPhase-TO remains comparatively uniform across the range. Columns are the achieved mixing fraction, grouped to the nearest nominal level.

| Method | 0.2 | 0.4 | 0.6 | 0.8 | 1.0 | Overall |
|---|---|---|---|---|---|---|
| *Nanopore panel (8 cell-line datasets, 40 mixtures)* | | | | | | |
| ASCAT (T+N) | 0.65 | 0.19 | 0.05 | 0.03 | 0.01 | 0.20 |
| ASCAT (TO) | 0.79 | 0.56 | 0.09 | 0.05 | 0.08 | 0.33 |
| PURPLE (TO) | 0.10 | 0.07 | 0.01 | 0.01 | 0.23 | 0.08 |
| LongPhase-TO (ClairS-TO) | 0.12 | 0.03 | 0.05 | 0.05 | 0.03 | 0.06 |
| LongPhase-TO (DeepSomatic-TO) | 0.05 | 0.02 | 0.04 | 0.06 | 0.04 | 0.04 |
| *PacBio HiFi panel (6 cell-line datasets, 30 mixtures)* | | | | | | |
| ASCAT (T+N) | 0.41 | 0.01 | 0.01 | 0.01 | 0.01 | 0.09 |
| ASCAT (TO) | 0.78 | 0.26 | 0.01 | 0.01 | 0.12 | 0.24 |
| PURPLE (TO) | 0.11 | 0.05 | 0.00 | 0.00 | 0.16 | 0.07 |
| LongPhase-TO (ClairS-TO) | 0.13 | 0.05 | 0.08 | 0.07 | 0.06 | 0.08 |
| LongPhase-TO (DeepSomatic-TO) | 0.04 | 0.03 | 0.05 | 0.08 | 0.09 | 0.06 |

**Supplementary Table 11**: Somatic SNV calling performance across DNA fraction levels for the six PacBio HiFi cell line datasets, with and without LongPhase-TO recalibration. Fraction is the tumor DNA fraction. The triplet-graph coefficients and thresholds were those fitted on the nanopore panel and were applied without refitting.

| Sample | Fraction | Variant caller | Precision | Recall | F1-score |
|---|---|---|---|---|---|
| HCC1395_PacBio | 0.2 | ClairS-TO | 0.436 | 0.376 | 0.403 |
| HCC1395_PacBio | 0.2 | ClairS-TO+LongPhase-TO | 0.733 | 0.417 | 0.531 |
| HCC1395_PacBio | 0.2 | DeepSomatic-TO | 0.510 | 0.720 | 0.597 |
| HCC1395_PacBio | 0.2 | DeepSomatic-TO+LongPhase-TO | 0.783 | 0.630 | 0.698 |
| HCC1395_PacBio | 0.4 | ClairS-TO | 0.566 | 0.631 | 0.597 |
| HCC1395_PacBio | 0.4 | ClairS-TO+LongPhase-TO | 0.807 | 0.645 | 0.717 |
| HCC1395_PacBio | 0.4 | DeepSomatic-TO | 0.548 | 0.855 | 0.668 |
| HCC1395_PacBio | 0.4 | DeepSomatic-TO+LongPhase-TO | 0.800 | 0.764 | 0.782 |
| HCC1395_PacBio | 0.6 | ClairS-TO | 0.605 | 0.731 | 0.662 |
| HCC1395_PacBio | 0.6 | ClairS-TO+LongPhase-TO | 0.827 | 0.730 | 0.775 |
| HCC1395_PacBio | 0.6 | DeepSomatic-TO | 0.558 | 0.907 | 0.691 |
| HCC1395_PacBio | 0.6 | DeepSomatic-TO+LongPhase-TO | 0.802 | 0.814 | 0.808 |
| HCC1395_PacBio | 0.8 | ClairS-TO | 0.650 | 0.779 | 0.709 |
| HCC1395_PacBio | 0.8 | ClairS-TO+LongPhase-TO | 0.847 | 0.737 | 0.788 |
| HCC1395_PacBio | 0.8 | DeepSomatic-TO | 0.568 | 0.934 | 0.707 |
| HCC1395_PacBio | 0.8 | DeepSomatic-TO+LongPhase-TO | 0.799 | 0.780 | 0.789 |
| HCC1395_PacBio | 1 | ClairS-TO | 0.730 | 0.811 | 0.769 |
| HCC1395_PacBio | 1 | ClairS-TO+LongPhase-TO | 0.767 | 0.734 | 0.750 |
| HCC1395_PacBio | 1 | DeepSomatic-TO | 0.650 | 0.949 | 0.772 |
| HCC1395_PacBio | 1 | DeepSomatic-TO+LongPhase-TO | 0.681 | 0.812 | 0.741 |
| COLO829_PacBio | 0.2 | ClairS-TO | 0.497 | 0.540 | 0.517 |
| COLO829_PacBio | 0.2 | ClairS-TO+LongPhase-TO | 0.736 | 0.553 | 0.631 |
| COLO829_PacBio | 0.2 | DeepSomatic-TO | 0.441 | 0.816 | 0.573 |
| COLO829_PacBio | 0.2 | DeepSomatic-TO+LongPhase-TO | 0.653 | 0.711 | 0.681 |
| COLO829_PacBio | 0.4 | ClairS-TO | 0.597 | 0.774 | 0.674 |
| COLO829_PacBio | 0.4 | ClairS-TO+LongPhase-TO | 0.805 | 0.741 | 0.771 |
| COLO829_PacBio | 0.4 | DeepSomatic-TO | 0.465 | 0.875 | 0.607 |
| COLO829_PacBio | 0.4 | DeepSomatic-TO+LongPhase-TO | 0.686 | 0.783 | 0.731 |
| COLO829_PacBio | 0.6 | ClairS-TO | 0.628 | 0.826 | 0.713 |
| COLO829_PacBio | 0.6 | ClairS-TO+LongPhase-TO | 0.838 | 0.778 | 0.807 |
| COLO829_PacBio | 0.6 | DeepSomatic-TO | 0.475 | 0.885 | 0.619 |
| COLO829_PacBio | 0.6 | DeepSomatic-TO+LongPhase-TO | 0.705 | 0.787 | 0.744 |
| COLO829_PacBio | 0.8 | ClairS-TO | 0.652 | 0.835 | 0.732 |
| COLO829_PacBio | 0.8 | ClairS-TO+LongPhase-TO | 0.857 | 0.713 | 0.779 |
| COLO829_PacBio | 0.8 | DeepSomatic-TO | 0.489 | 0.883 | 0.630 |
| COLO829_PacBio | 0.8 | DeepSomatic-TO+LongPhase-TO | 0.713 | 0.689 | 0.700 |
| COLO829_PacBio | 1 | ClairS-TO | 0.680 | 0.838 | 0.751 |
| COLO829_PacBio | 1 | ClairS-TO+LongPhase-TO | 0.656 | 0.728 | 0.690 |
| COLO829_PacBio | 1 | DeepSomatic-TO | 0.524 | 0.866 | 0.653 |
| COLO829_PacBio | 1 | DeepSomatic-TO+LongPhase-TO | 0.514 | 0.702 | 0.593 |
| HCC1954_CASTLE | 0.2 | ClairS-TO | 0.055 | 0.160 | 0.082 |
| HCC1954_CASTLE | 0.2 | ClairS-TO+LongPhase-TO | 0.225 | 0.204 | 0.214 |
| HCC1954_CASTLE | 0.2 | DeepSomatic-TO | 0.148 | 0.601 | 0.237 |
| HCC1954_CASTLE | 0.2 | DeepSomatic-TO+LongPhase-TO | 0.516 | 0.532 | 0.524 |
| HCC1954_CASTLE | 0.4 | ClairS-TO | 0.144 | 0.464 | 0.220 |
| HCC1954_CASTLE | 0.4 | ClairS-TO+LongPhase-TO | 0.425 | 0.519 | 0.467 |
| HCC1954_CASTLE | 0.4 | DeepSomatic-TO | 0.196 | 0.843 | 0.318 |
| HCC1954_CASTLE | 0.4 | DeepSomatic-TO+LongPhase-TO | 0.593 | 0.760 | 0.666 |
| HCC1954_CASTLE | 0.6 | ClairS-TO | 0.201 | 0.686 | 0.311 |
| HCC1954_CASTLE | 0.6 | ClairS-TO+LongPhase-TO | 0.510 | 0.714 | 0.595 |
| HCC1954_CASTLE | 0.6 | DeepSomatic-TO | 0.209 | 0.910 | 0.340 |
| HCC1954_CASTLE | 0.6 | DeepSomatic-TO+LongPhase-TO | 0.614 | 0.826 | 0.704 |
| HCC1954_CASTLE | 0.8 | ClairS-TO | 0.234 | 0.801 | 0.362 |
| HCC1954_CASTLE | 0.8 | ClairS-TO+LongPhase-TO | 0.560 | 0.792 | 0.656 |
| HCC1954_CASTLE | 0.8 | DeepSomatic-TO | 0.214 | 0.927 | 0.347 |
| HCC1954_CASTLE | 0.8 | DeepSomatic-TO+LongPhase-TO | 0.625 | 0.831 | 0.713 |
| HCC1954_CASTLE | 1 | ClairS-TO | 0.258 | 0.849 | 0.396 |
| HCC1954_CASTLE | 1 | ClairS-TO+LongPhase-TO | 0.584 | 0.676 | 0.626 |
| HCC1954_CASTLE | 1 | DeepSomatic-TO | 0.229 | 0.923 | 0.367 |

| Sample | Fraction | Variant caller | Precision | Recall | F1-score |
|---|---|---|---|---|---|
| HCC1954_CASTLE | 1 | DeepSomatic-TO+LongPhase-TO | 0.643 | 0.648 | 0.645 |
| HCC1937_CASTLE | 0.2 | ClairS-TO | 0.199 | 0.310 | 0.242 |
| HCC1937_CASTLE | 0.2 | ClairS-TO+LongPhase-TO | 0.473 | 0.349 | 0.402 |
| HCC1937_CASTLE | 0.2 | DeepSomatic-TO | 0.271 | 0.658 | 0.384 |
| HCC1937_CASTLE | 0.2 | DeepSomatic-TO+LongPhase-TO | 0.567 | 0.574 | 0.570 |
| HCC1937_CASTLE | 0.4 | ClairS-TO | 0.312 | 0.559 | 0.401 |
| HCC1937_CASTLE | 0.4 | ClairS-TO+LongPhase-TO | 0.611 | 0.580 | 0.595 |
| HCC1937_CASTLE | 0.4 | DeepSomatic-TO | 0.318 | 0.832 | 0.460 |
| HCC1937_CASTLE | 0.4 | DeepSomatic-TO+LongPhase-TO | 0.617 | 0.738 | 0.672 |
| HCC1937_CASTLE | 0.6 | ClairS-TO | 0.364 | 0.682 | 0.474 |
| HCC1937_CASTLE | 0.6 | ClairS-TO+LongPhase-TO | 0.667 | 0.689 | 0.678 |
| HCC1937_CASTLE | 0.6 | DeepSomatic-TO | 0.332 | 0.898 | 0.485 |
| HCC1937_CASTLE | 0.6 | DeepSomatic-TO+LongPhase-TO | 0.630 | 0.798 | 0.704 |
| HCC1937_CASTLE | 0.8 | ClairS-TO | 0.426 | 0.757 | 0.546 |
| HCC1937_CASTLE | 0.8 | ClairS-TO+LongPhase-TO | 0.723 | 0.722 | 0.722 |
| HCC1937_CASTLE | 0.8 | DeepSomatic-TO | 0.343 | 0.925 | 0.501 |
| HCC1937_CASTLE | 0.8 | DeepSomatic-TO+LongPhase-TO | 0.628 | 0.775 | 0.694 |
| HCC1937_CASTLE | 1 | ClairS-TO | 0.523 | 0.808 | 0.635 |
| HCC1937_CASTLE | 1 | ClairS-TO+LongPhase-TO | 0.612 | 0.763 | 0.679 |
| HCC1937_CASTLE | 1 | DeepSomatic-TO | 0.425 | 0.930 | 0.583 |
| HCC1937_CASTLE | 1 | DeepSomatic-TO+LongPhase-TO | 0.485 | 0.832 | 0.613 |
| H2009_CASTLE | 0.2 | ClairS-TO | 0.767 | 0.547 | 0.639 |
| H2009_CASTLE | 0.2 | ClairS-TO+LongPhase-TO | 0.889 | 0.591 | 0.710 |
| H2009_CASTLE | 0.2 | DeepSomatic-TO | 0.800 | 0.880 | 0.838 |
| H2009_CASTLE | 0.2 | DeepSomatic-TO+LongPhase-TO | 0.903 | 0.780 | 0.837 |
| H2009_CASTLE | 0.4 | ClairS-TO | 0.840 | 0.853 | 0.846 |
| H2009_CASTLE | 0.4 | ClairS-TO+LongPhase-TO | 0.918 | 0.843 | 0.879 |
| H2009_CASTLE | 0.4 | DeepSomatic-TO | 0.806 | 0.951 | 0.873 |
| H2009_CASTLE | 0.4 | DeepSomatic-TO+LongPhase-TO | 0.901 | 0.865 | 0.882 |
| H2009_CASTLE | 0.6 | ClairS-TO | 0.854 | 0.908 | 0.880 |
| H2009_CASTLE | 0.6 | ClairS-TO+LongPhase-TO | 0.927 | 0.881 | 0.903 |
| H2009_CASTLE | 0.6 | DeepSomatic-TO | 0.800 | 0.958 | 0.872 |
| H2009_CASTLE | 0.6 | DeepSomatic-TO+LongPhase-TO | 0.893 | 0.870 | 0.881 |
| H2009_CASTLE | 0.8 | ClairS-TO | 0.878 | 0.914 | 0.895 |
| H2009_CASTLE | 0.8 | ClairS-TO+LongPhase-TO | 0.942 | 0.809 | 0.871 |
| H2009_CASTLE | 0.8 | DeepSomatic-TO | 0.797 | 0.953 | 0.868 |
| H2009_CASTLE | 0.8 | DeepSomatic-TO+LongPhase-TO | 0.881 | 0.767 | 0.820 |
| H2009_CASTLE | 1 | ClairS-TO | 0.912 | 0.917 | 0.915 |
| H2009_CASTLE | 1 | ClairS-TO+LongPhase-TO | 0.912 | 0.840 | 0.875 |
| H2009_CASTLE | 1 | DeepSomatic-TO | 0.833 | 0.924 | 0.876 |
| H2009_CASTLE | 1 | DeepSomatic-TO+LongPhase-TO | 0.796 | 0.195 | 0.314 |
| H1437_CASTLE | 0.2 | ClairS-TO | 0.551 | 0.351 | 0.429 |
| H1437_CASTLE | 0.2 | ClairS-TO+LongPhase-TO | 0.775 | 0.379 | 0.509 |
| H1437_CASTLE | 0.2 | DeepSomatic-TO | 0.581 | 0.564 | 0.573 |
| H1437_CASTLE | 0.2 | DeepSomatic-TO+LongPhase-TO | 0.797 | 0.500 | 0.615 |
| H1437_CASTLE | 0.4 | ClairS-TO | 0.651 | 0.529 | 0.584 |
| H1437_CASTLE | 0.4 | ClairS-TO+LongPhase-TO | 0.829 | 0.528 | 0.645 |
| H1437_CASTLE | 0.4 | DeepSomatic-TO | 0.604 | 0.622 | 0.613 |
| H1437_CASTLE | 0.4 | DeepSomatic-TO+LongPhase-TO | 0.810 | 0.561 | 0.663 |
| H1437_CASTLE | 0.6 | ClairS-TO | 0.675 | 0.576 | 0.622 |
| H1437_CASTLE | 0.6 | ClairS-TO+LongPhase-TO | 0.846 | 0.565 | 0.677 |
| H1437_CASTLE | 0.6 | DeepSomatic-TO | 0.609 | 0.636 | 0.622 |
| H1437_CASTLE | 0.6 | DeepSomatic-TO+LongPhase-TO | 0.811 | 0.574 | 0.672 |
| H1437_CASTLE | 0.8 | ClairS-TO | 0.709 | 0.589 | 0.643 |
| H1437_CASTLE | 0.8 | ClairS-TO+LongPhase-TO | 0.868 | 0.534 | 0.661 |
| H1437_CASTLE | 0.8 | DeepSomatic-TO | 0.613 | 0.639 | 0.626 |
| H1437_CASTLE | 0.8 | DeepSomatic-TO+LongPhase-TO | 0.801 | 0.517 | 0.628 |
| H1437_CASTLE | 1 | ClairS-TO | 0.773 | 0.598 | 0.674 |
| H1437_CASTLE | 1 | ClairS-TO+LongPhase-TO | 0.778 | 0.524 | 0.627 |
| H1437_CASTLE | 1 | DeepSomatic-TO | 0.678 | 0.623 | 0.649 |
| H1437_CASTLE | 1 | DeepSomatic-TO+LongPhase-TO | 0.677 | 0.511 | 0.583 |

**Supplementary Table 12**: Somatic indel calling performance across DNA fraction levels for the six PacBio HiFi cell line datasets, with and without LongPhase-TO recalibration. Fraction is the tumor DNA fraction. The triplet-graph coefficients and thresholds were those fitted on the nanopore panel and were applied without refitting.

| Sample | Fraction | Variant caller | Precision | Recall | F1-score |
|---|---|---|---|---|---|
| HCC1395_PacBio | 0.2 | ClairS-TO | 0.020 | 0.014 | 0.017 |
| HCC1395_PacBio | 0.2 | ClairS-TO+LongPhase-TO | 0.051 | 0.016 | 0.024 |
| HCC1395_PacBio | 0.2 | DeepSomatic-TO | 0.114 | 0.173 | 0.137 |
| HCC1395_PacBio | 0.2 | DeepSomatic-TO+LongPhase-TO | 0.249 | 0.153 | 0.190 |
| HCC1395_PacBio | 0.4 | ClairS-TO | 0.135 | 0.103 | 0.117 |
| HCC1395_PacBio | 0.4 | ClairS-TO+LongPhase-TO | 0.298 | 0.107 | 0.157 |
| HCC1395_PacBio | 0.4 | DeepSomatic-TO | 0.275 | 0.479 | 0.349 |
| HCC1395_PacBio | 0.4 | DeepSomatic-TO+LongPhase-TO | 0.484 | 0.421 | 0.450 |
| HCC1395_PacBio | 0.6 | ClairS-TO | 0.317 | 0.248 | 0.279 |
| HCC1395_PacBio | 0.6 | ClairS-TO+LongPhase-TO | 0.565 | 0.248 | 0.345 |
| HCC1395_PacBio | 0.6 | DeepSomatic-TO | 0.369 | 0.627 | 0.465 |
| HCC1395_PacBio | 0.6 | DeepSomatic-TO+LongPhase-TO | 0.602 | 0.537 | 0.568 |
| HCC1395_PacBio | 0.8 | ClairS-TO | 0.448 | 0.330 | 0.380 |
| HCC1395_PacBio | 0.8 | ClairS-TO+LongPhase-TO | 0.675 | 0.313 | 0.427 |
| HCC1395_PacBio | 0.8 | DeepSomatic-TO | 0.460 | 0.706 | 0.557 |
| HCC1395_PacBio | 0.8 | DeepSomatic-TO+LongPhase-TO | 0.663 | 0.574 | 0.616 |
| HCC1395_PacBio | 1 | ClairS-TO | 0.494 | 0.383 | 0.431 |
| HCC1395_PacBio | 1 | ClairS-TO+LongPhase-TO | 0.482 | 0.408 | 0.442 |
| HCC1395_PacBio | 1 | DeepSomatic-TO | 0.560 | 0.766 | 0.647 |
| HCC1395_PacBio | 1 | DeepSomatic-TO+LongPhase-TO | 0.565 | 0.741 | 0.641 |
| COLO829_PacBio | 0.2 | ClairS-TO | 0.012 | 0.017 | 0.014 |
| COLO829_PacBio | 0.2 | ClairS-TO+LongPhase-TO | 0.026 | 0.015 | 0.019 |
| COLO829_PacBio | 0.2 | DeepSomatic-TO | 0.042 | 0.128 | 0.063 |
| COLO829_PacBio | 0.2 | DeepSomatic-TO+LongPhase-TO | 0.100 | 0.112 | 0.105 |
| COLO829_PacBio | 0.4 | ClairS-TO | 0.090 | 0.138 | 0.109 |
| COLO829_PacBio | 0.4 | ClairS-TO+LongPhase-TO | 0.209 | 0.137 | 0.166 |
| COLO829_PacBio | 0.4 | DeepSomatic-TO | 0.080 | 0.246 | 0.120 |
| COLO829_PacBio | 0.4 | DeepSomatic-TO+LongPhase-TO | 0.181 | 0.217 | 0.198 |
| COLO829_PacBio | 0.6 | ClairS-TO | 0.153 | 0.231 | 0.184 |
| COLO829_PacBio | 0.6 | ClairS-TO+LongPhase-TO | 0.329 | 0.236 | 0.275 |
| COLO829_PacBio | 0.6 | DeepSomatic-TO | 0.091 | 0.268 | 0.136 |
| COLO829_PacBio | 0.6 | DeepSomatic-TO+LongPhase-TO | 0.205 | 0.231 | 0.217 |
| COLO829_PacBio | 0.8 | ClairS-TO | 0.188 | 0.269 | 0.221 |
| COLO829_PacBio | 0.8 | ClairS-TO+LongPhase-TO | 0.354 | 0.240 | 0.286 |
| COLO829_PacBio | 0.8 | DeepSomatic-TO | 0.091 | 0.236 | 0.132 |
| COLO829_PacBio | 0.8 | DeepSomatic-TO+LongPhase-TO | 0.163 | 0.166 | 0.164 |
| COLO829_PacBio | 1 | ClairS-TO | 0.206 | 0.289 | 0.241 |
| COLO829_PacBio | 1 | ClairS-TO+LongPhase-TO | 0.184 | 0.318 | 0.233 |
| COLO829_PacBio | 1 | DeepSomatic-TO | 0.071 | 0.169 | 0.100 |
| COLO829_PacBio | 1 | DeepSomatic-TO+LongPhase-TO | 0.072 | 0.161 | 0.100 |
| HCC1954_CASTLE | 0.2 | ClairS-TO | 0.002 | 0.004 | 0.003 |
| HCC1954_CASTLE | 0.2 | ClairS-TO+LongPhase-TO | 0.006 | 0.004 | 0.005 |
| HCC1954_CASTLE | 0.2 | DeepSomatic-TO | 0.013 | 0.019 | 0.015 |
| HCC1954_CASTLE | 0.2 | DeepSomatic-TO+LongPhase-TO | 0.044 | 0.017 | 0.025 |
| HCC1954_CASTLE | 0.4 | ClairS-TO | 0.018 | 0.027 | 0.021 |
| HCC1954_CASTLE | 0.4 | ClairS-TO+LongPhase-TO | 0.048 | 0.028 | 0.036 |
| HCC1954_CASTLE | 0.4 | DeepSomatic-TO | 0.070 | 0.107 | 0.084 |
| HCC1954_CASTLE | 0.4 | DeepSomatic-TO+LongPhase-TO | 0.199 | 0.095 | 0.129 |
| HCC1954_CASTLE | 0.6 | ClairS-TO | 0.064 | 0.093 | 0.076 |
| HCC1954_CASTLE | 0.6 | ClairS-TO+LongPhase-TO | 0.185 | 0.105 | 0.134 |
| HCC1954_CASTLE | 0.6 | DeepSomatic-TO | 0.128 | 0.197 | 0.155 |
| HCC1954_CASTLE | 0.6 | DeepSomatic-TO+LongPhase-TO | 0.353 | 0.176 | 0.235 |
| HCC1954_CASTLE | 0.8 | ClairS-TO | 0.129 | 0.182 | 0.151 |
| HCC1954_CASTLE | 0.8 | ClairS-TO+LongPhase-TO | 0.334 | 0.197 | 0.248 |
| HCC1954_CASTLE | 0.8 | DeepSomatic-TO | 0.175 | 0.254 | 0.207 |
| HCC1954_CASTLE | 0.8 | DeepSomatic-TO+LongPhase-TO | 0.436 | 0.218 | 0.291 |
| HCC1954_CASTLE | 1 | ClairS-TO | 0.193 | 0.269 | 0.225 |
| HCC1954_CASTLE | 1 | ClairS-TO+LongPhase-TO | 0.377 | 0.218 | 0.276 |
| HCC1954_CASTLE | 1 | DeepSomatic-TO | 0.190 | 0.261 | 0.219 |

| Sample | Fraction | Variant caller | Precision | Recall | F1-score |
|---|---|---|---|---|---|
| HCC1954_CASTLE | 1 | DeepSomatic-TO+LongPhase-TO | 0.396 | 0.186 | 0.253 |
| HCC1937_CASTLE | 0.2 | ClairS-TO | 0.018 | 0.025 | 0.021 |
| HCC1937_CASTLE | 0.2 | ClairS-TO+LongPhase-TO | 0.037 | 0.024 | 0.029 |
| HCC1937_CASTLE | 0.2 | DeepSomatic-TO | 0.104 | 0.252 | 0.147 |
| HCC1937_CASTLE | 0.2 | DeepSomatic-TO+LongPhase-TO | 0.257 | 0.224 | 0.239 |
| HCC1937_CASTLE | 0.4 | ClairS-TO | 0.134 | 0.188 | 0.156 |
| HCC1937_CASTLE | 0.4 | ClairS-TO+LongPhase-TO | 0.291 | 0.193 | 0.232 |
| HCC1937_CASTLE | 0.4 | DeepSomatic-TO | 0.219 | 0.521 | 0.308 |
| HCC1937_CASTLE | 0.4 | DeepSomatic-TO+LongPhase-TO | 0.464 | 0.464 | 0.464 |
| HCC1937_CASTLE | 0.6 | ClairS-TO | 0.245 | 0.320 | 0.278 |
| HCC1937_CASTLE | 0.6 | ClairS-TO+LongPhase-TO | 0.463 | 0.302 | 0.365 |
| HCC1937_CASTLE | 0.6 | DeepSomatic-TO | 0.282 | 0.615 | 0.387 |
| HCC1937_CASTLE | 0.6 | DeepSomatic-TO+LongPhase-TO | 0.536 | 0.506 | 0.521 |
| HCC1937_CASTLE | 0.8 | ClairS-TO | 0.323 | 0.375 | 0.347 |
| HCC1937_CASTLE | 0.8 | ClairS-TO+LongPhase-TO | 0.541 | 0.320 | 0.403 |
| HCC1937_CASTLE | 0.8 | DeepSomatic-TO | 0.364 | 0.614 | 0.457 |
| HCC1937_CASTLE | 0.8 | DeepSomatic-TO+LongPhase-TO | 0.583 | 0.455 | 0.512 |
| HCC1937_CASTLE | 1 | ClairS-TO | 0.350 | 0.423 | 0.383 |
| HCC1937_CASTLE | 1 | ClairS-TO+LongPhase-TO | 0.347 | 0.415 | 0.378 |
| HCC1937_CASTLE | 1 | DeepSomatic-TO | 0.409 | 0.516 | 0.457 |
| HCC1937_CASTLE | 1 | DeepSomatic-TO+LongPhase-TO | 0.415 | 0.497 | 0.452 |
| H2009_CASTLE | 0.2 | ClairS-TO | 0.096 | 0.030 | 0.046 |
| H2009_CASTLE | 0.2 | ClairS-TO+LongPhase-TO | 0.197 | 0.033 | 0.057 |
| H2009_CASTLE | 0.2 | DeepSomatic-TO | 0.283 | 0.174 | 0.216 |
| H2009_CASTLE | 0.2 | DeepSomatic-TO+LongPhase-TO | 0.473 | 0.156 | 0.235 |
| H2009_CASTLE | 0.4 | ClairS-TO | 0.468 | 0.230 | 0.308 |
| H2009_CASTLE | 0.4 | ClairS-TO+LongPhase-TO | 0.655 | 0.233 | 0.344 |
| H2009_CASTLE | 0.4 | DeepSomatic-TO | 0.505 | 0.429 | 0.464 |
| H2009_CASTLE | 0.4 | DeepSomatic-TO+LongPhase-TO | 0.702 | 0.385 | 0.497 |
| H2009_CASTLE | 0.6 | ClairS-TO | 0.677 | 0.436 | 0.530 |
| H2009_CASTLE | 0.6 | ClairS-TO+LongPhase-TO | 0.834 | 0.406 | 0.546 |
| H2009_CASTLE | 0.6 | DeepSomatic-TO | 0.579 | 0.499 | 0.536 |
| H2009_CASTLE | 0.6 | DeepSomatic-TO+LongPhase-TO | 0.765 | 0.422 | 0.544 |
| H2009_CASTLE | 0.8 | ClairS-TO | 0.776 | 0.579 | 0.663 |
| H2009_CASTLE | 0.8 | ClairS-TO+LongPhase-TO | 0.884 | 0.461 | 0.606 |
| H2009_CASTLE | 0.8 | DeepSomatic-TO | 0.601 | 0.431 | 0.502 |
| H2009_CASTLE | 0.8 | DeepSomatic-TO+LongPhase-TO | 0.758 | 0.308 | 0.438 |
| H2009_CASTLE | 1 | ClairS-TO | 0.791 | 0.633 | 0.703 |
| H2009_CASTLE | 1 | ClairS-TO+LongPhase-TO | 0.778 | 0.659 | 0.714 |
| H2009_CASTLE | 1 | DeepSomatic-TO | 0.571 | 0.295 | 0.389 |
| H2009_CASTLE | 1 | DeepSomatic-TO+LongPhase-TO | 0.604 | 0.104 | 0.178 |
| H1437_CASTLE | 0.2 | ClairS-TO | 0.029 | 0.013 | 0.017 |
| H1437_CASTLE | 0.2 | ClairS-TO+LongPhase-TO | 0.062 | 0.014 | 0.023 |
| H1437_CASTLE | 0.2 | DeepSomatic-TO | 0.097 | 0.113 | 0.104 |
| H1437_CASTLE | 0.2 | DeepSomatic-TO+LongPhase-TO | 0.259 | 0.100 | 0.144 |
| H1437_CASTLE | 0.4 | ClairS-TO | 0.281 | 0.141 | 0.188 |
| H1437_CASTLE | 0.4 | ClairS-TO+LongPhase-TO | 0.472 | 0.146 | 0.223 |
| H1437_CASTLE | 0.4 | DeepSomatic-TO | 0.239 | 0.255 | 0.246 |
| H1437_CASTLE | 0.4 | DeepSomatic-TO+LongPhase-TO | 0.481 | 0.228 | 0.309 |
| H1437_CASTLE | 0.6 | ClairS-TO | 0.457 | 0.243 | 0.317 |
| H1437_CASTLE | 0.6 | ClairS-TO+LongPhase-TO | 0.672 | 0.239 | 0.352 |
| H1437_CASTLE | 0.6 | DeepSomatic-TO | 0.329 | 0.295 | 0.311 |
| H1437_CASTLE | 0.6 | DeepSomatic-TO+LongPhase-TO | 0.580 | 0.252 | 0.351 |
| H1437_CASTLE | 0.8 | ClairS-TO | 0.574 | 0.297 | 0.391 |
| H1437_CASTLE | 0.8 | ClairS-TO+LongPhase-TO | 0.752 | 0.246 | 0.371 |
| H1437_CASTLE | 0.8 | DeepSomatic-TO | 0.386 | 0.257 | 0.308 |
| H1437_CASTLE | 0.8 | DeepSomatic-TO+LongPhase-TO | 0.601 | 0.186 | 0.284 |
| H1437_CASTLE | 1 | ClairS-TO | 0.609 | 0.333 | 0.430 |
| H1437_CASTLE | 1 | ClairS-TO+LongPhase-TO | 0.598 | 0.346 | 0.438 |
| H1437_CASTLE | 1 | DeepSomatic-TO | 0.407 | 0.200 | 0.268 |
| H1437_CASTLE | 1 | DeepSomatic-TO+LongPhase-TO | 0.407 | 0.191 | 0.260 |

**Supplementary Table 13**: Per-mixture tumor DNA fraction estimates for the PacBio HiFi panel, comparing ASCAT, PURPLE and LongPhase-TO across six cell-line datasets and five DNA-fraction levels (30 mixtures). ASCAT and PURPLE values are cellular purities converted to DNA fraction with each tool's fitted ploidy (see Methods). The LongPhase-TO regression coefficients were those fitted on the nanopore panel and were applied without refitting. LP-TO (C) and LP-TO (D) denote LongPhase-TO applied to somatic variants from ClairS-TO and DeepSomatic-TO, respectively. The expected value is the achieved read-level mixing fraction, which differs slightly from the nominal target where coverage was limiting. Entries marked n.a. indicate that the method returned no estimate.

| Dataset | Expected | ASCAT (T+N) | ASCAT (TO) | PURPLE (TO) | LP-TO (C) | LP-TO (D) |
|---|---|---|---|---|---|---|
| HCC1395_PacBio | 0.200 | 1.000 | 0.937 | 0.340 | 0.146 | 0.148 |
| HCC1395_PacBio | 0.400 | 0.406 | 1.000 | 0.389 | 0.383 | 0.389 |
| HCC1395_PacBio | 0.600 | 0.609 | 0.598 | 0.598 | 0.580 | 0.580 |
| HCC1395_PacBio | 0.800 | 0.809 | 0.807 | 0.798 | 0.742 | 0.704 |
| HCC1395_PacBio | 1.000 | 1.000 | 1.000 | 0.946 | 1.000 | 1.000 |
| COLO829_PacBio | 0.200 | 0.221 | 0.971 | 0.197 | 0.121 | 0.165 |
| COLO829_PacBio | 0.400 | 0.408 | 1.000 | 0.390 | 0.409 | 0.403 |
| COLO829_PacBio | 0.600 | 0.604 | 0.610 | 0.604 | 0.693 | 0.656 |
| COLO829_PacBio | 0.800 | 0.810 | 0.810 | 0.797 | 0.833 | 0.821 |
| COLO829_PacBio | 1.000 | 1.000 | n.a. | 0.994 | 1.000 | 0.989 |
| HCC1954_CASTLE | 0.200 | 1.000 | 1.000 | 0.231 | 0.619 | 0.186 |
| HCC1954_CASTLE | 0.400 | 0.411 | 0.445 | 0.399 | 0.250 | 0.289 |
| HCC1954_CASTLE | 0.600 | 0.598 | 0.619 | 0.590 | 0.307 | 0.429 |
| HCC1954_CASTLE | 0.800 | 0.791 | 0.809 | 0.793 | 0.478 | 0.552 |
| HCC1954_CASTLE | 1.000 | 0.982 | 0.581 | 0.567 | 0.725 | 0.651 |
| HCC1937_CASTLE | 0.200 | 0.214 | 1.000 | 0.369 | 0.030 | 0.176 |
| HCC1937_CASTLE | 0.400 | 0.404 | 0.402 | 0.399 | 0.304 | 0.378 |
| HCC1937_CASTLE | 0.600 | 0.602 | 0.602 | 0.606 | 0.554 | 0.590 |
| HCC1937_CASTLE | 0.800 | 0.801 | 0.806 | 0.793 | 0.791 | 0.736 |
| HCC1937_CASTLE | 1.000 | 0.995 | 0.935 | 0.619 | 0.928 | 1.000 |
| H2009_CASTLE | 0.200 | 1.000 | 1.000 | 0.364 | 0.153 | 0.155 |
| H2009_CASTLE | 0.400 | 0.384 | 0.395 | 0.399 | 0.369 | 0.374 |
| H2009_CASTLE | 0.600 | 0.584 | 0.597 | 0.600 | 0.617 | 0.620 |
| H2009_CASTLE | 0.800 | 0.791 | 0.798 | 0.790 | 0.816 | 0.783 |
| H2009_CASTLE | 1.000 | 0.983 | 0.932 | 0.940 | 1.000 | 0.844 |
| H1437_CASTLE | 0.200 | 0.216 | 0.956 | 0.353 | 0.165 | 0.156 |
| H1437_CASTLE | 0.400 | 0.392 | 0.679 | 0.680 | 0.385 | 0.406 |
| H1437_CASTLE | 0.600 | 0.590 | 0.604 | 0.604 | 0.622 | 0.616 |
| H1437_CASTLE | 0.800 | 0.787 | 0.802 | 0.800 | 0.795 | 0.751 |
| H1437_CASTLE | 1.000 | 0.986 | 0.936 | 0.994 | 1.000 | 1.000 |

**Supplementary Table 15**: Somatic variant callers used for benchmarking.

| Somatic Variant Callers |
|---|
| ClairS-TO v0.3.0 (ssrs) |
| ClairS-TO v0.3.0 (ss) |
| DeepSomatic v1.8.0 (tumor-only) |

**Supplementary Table 14**: Fixed parameters for directional clipping and LOH classification. Distances expressed in positions refer to the ordered series of clip-bearing genomic positions, not base pairs.

| Parameter | Value | Function |
|---|---|---|
| Minimum clipping length | $> 5$ bases | Exclude very short clipping operations. |
| Directional rolling window | 100 clip-bearing positions | Consolidate dispersed directional clipping support. |
| Directional rolling passes | 2 | Generate the second-pass profile used to localize lower-amplitude candidate peaks. |
| Transformed peak threshold | $\geq 0.25$ | Retain distributed-signal candidates. |
| Peak consolidation distance | 100 clip-bearing positions | Represent nearby candidates by the larger peak. |
| Direct-signal exclusion span | 200 clip-bearing positions | Avoid duplicating a direct breakpoint as a distributed-signal candidate. |
| Direct signal/event initiation | $\geq 5$ clips | Retain a direct signal and initiate interval pairing. |
| Cumulative-profile smoothing window | 100 clip-bearing positions | Reduce local variation before estimating candidate support. |
| Candidate-support offset | $\pm 10$ clip-bearing positions | Measure the local change in the smoothed cumulative profile. |
| Maximum SGE pairing span | 10 kb | Restrict paired opposite-orientation signals to localized events. |
| Opposite-orientation support | $\geq \lfloor c/4 \rfloor$ | Complete an SGE interval relative to initiating-orientation support $c$. |
| Terminal LGE boundaries | First and last clip-bearing positions | Bound evaluation at the ends of each analyzed chromosome. |
| Homozygous-variant threshold | VAF $\geq 0.8$ or input homozygous flag | Classify variants for regional allele-state counting. |
| LOH threshold | $R_{\text{het}} < 0.09$ | Classify an LGE-bounded region as LOH. |

**Supplementary Table 16**: PacBio HiFi cell line datasets used to assess cross-platform generalization of somatic recalibration. All datasets were generated on the PacBio Revio platform and the same benchmark truth sets as the nanopore panel were applied. Four datasets were obtained from CASTLE, and the HCC1395 and COLO829 pairs from the public PacBio Revio 2023Q2 release. Five of the six cell lines, all except COLO829, belong to the tumor-normal cohort assembled for DeepSomatic; that cohort also includes Hs578T, which was not analyzed here.

| Dataset | Cancer Type | Source | Benchmark | Normal | Tumor |
|---|---|---|---|---|---|
| HCC1395_PacBio | Breast ductal carcinoma | PacBio | SEQC2 | 44x | 62x |
| COLO829_PacBio | Malignant melanoma | PacBio | NYGC | n.r. | 41x |
| HCC1954_CASTLE | Breast ductal carcinoma | CASTLE | DeepSomatic | 67x | 67x |
| HCC1937_CASTLE | Breast ductal carcinoma | CASTLE | DeepSomatic | 60x | 64x |
| H2009_CASTLE | Non-small-cell lung cancer | CASTLE | DeepSomatic | 73x | 70x |
| H1437_CASTLE | Non-small-cell lung cancer | CASTLE | DeepSomatic | 76x | 69x |

All libraries were prepared by Megaruptor 3 shearing to approximately 15–20 kb followed by SMRTbell preparation, sequenced on PacBio Revio with 24-hour movies with HiFi 5mC calls, and aligned to GRCh38 (no-alt) with `pbmm2`; the reported read N50 ranges from 15 to 18 kb. The HCC1937, HCC1954, H1437 and H2009 tumor-normal pairs were obtained from CASTLE, and the HCC1395 and COLO829 pairs, with their matched HCC1395-BL and COLO829-BL normals, from the public PacBio Revio 2023Q2 release. The COLO829 release does not report coverage and has been used elsewhere as an approximately 50× PacBio benchmark. In our alignments the pure COLO829 tumor sample reached approximately 41×, the value listed in the table, rather than the 50× used for the other mixtures, and its 1.0 DNA fraction point was evaluated at that coverage. The coverage of the matched COLO829-BL normal is not reported (n.r.).

### 5 Command-line arguments

#### 5.1 Read Alignment

##### 5.1.1 Alignment

**minimap2** (https://github.com/lh3/minimap2)

```
minimap2 --MD -t {THREADS} -a {REFERENCE} {INPUT_FASTA} -o {OUTPUT_SAM}
samtools view -@ {THREADS} -bS {OUTPUT_SAM} > {OUTPUT_BAM}
samtools sort -@ {THREADS} {OUTPUT_BAM} -o {OUTPUT_BAM_SORTED}
samtools index {OUTPUT_BAM_SORTED}
```

##### 5.1.2 Downsampling

**mosdepth** (https://github.com/brentp/mosdepth)

```
mosdepth -t {THREADS} -n -x --quantize 0:15:150: {SAMPLE_BAM}
# get depth profile to determine the downsampling fraction
samtools view -@ {THREADS} -bh --subsample {FRACTION} {SAMPLE_BAM} -o {DOWN_SAMPLED_BAM}
samtools index {DOWN_SAMPLED_BAM}
```

#### 5.2 Variant Calling

##### 5.2.1 ClairS-TO v0.3.0

(https://github.com/HKU-BAL/ClairS-TO)

```
# nanopore MODEL = ont_r10_dorado_sup_5khz_ssrs (default) or ont_r10_dorado_sup_5khz_ss
# PacBio HiFi MODEL = hifi_revio_ssrs (default) or hifi_revio_ss
docker run -i --rm \
  -v ${INPUT_DIR}:${INPUT_DIR} \
  -v ${OUTPUT_DIR}:${OUTPUT_DIR} \
  -u $(id -u):$(id -g) \
  hkubal/clairs-to:v0.3.0 \
  /opt/bin/run_clairs_to \
  --tumor_bam_fn ${TUMOR_BAM} \
  --ref_fn ${REF} \
  --threads ${THREADS} \
  --platform ${MODEL} \
  --output_dir ${OUTPUT_DIR}
```

**DeepSomatic v1.8.0** (https://github.com/google/deepsomatic)

```
docker run -i --rm --gpus all \
  -v ${INPUT_DIR}:${INPUT_DIR} \
  -v ${OUTPUT_DIR}:${OUTPUT_DIR} \
  -u $(id -u):$(id -g) \
  google/deepsomatic:1.8.0-gpu \
  run_deepsomatic \
  --model_type ${MODEL_TYPE} \
  # MODEL_TYPE = ONT_TUMOR_ONLY (nanopore) or PACBIO_TUMOR_ONLY (PacBio HiFi)
  --ref ${REF} \
  --reads_tumor  ${TUMOR_BAM} \
  --output_vcf  ${OUTPUT_DIR}/${OUTPUT_VCF_FILE_PATH} \
  --sample_name_tumor ${SAMPLE_NAME_TUMOR} \
  --num_shards ${THREADS} \
  --logging_dir ${OUTPUT_DIR}/logs \
  --intermediate_results_dir ${OUTPUT_DIR}/intermediate_results_dir \
  --use_default_pon_filtering=true
```

#### 5.3 Phasing

**LongPhase-TO v1.0.0** (https://github.com/CCU-Bioinformatics-Lab/longphase-to)

```
# CALLER = clairs_to_ss, clairs_to_ssrs or deepsomatic_to
# platform preset: --ont (nanopore) or --pb (PacBio HiFi); --ont was used throughout this study
# add --loh to write the detected LOH segments as a BED file
longphase-to phase \
-b ${TUMOR_BAM} \
-r ${REF} \
-s ${VCF} \
-t ${THREADS} \
--caller ${CALLER} \
--pon-file ${PON_FILE} \
--strict-pon-file ${STRICT_PON_FILE} \
--ont \
-o ${OUTPUT}
```

LongPhase-TO provides a `--pb` preset for PacBio HiFi input. In the cross-platform analysis (Supplementary Figs. 4, 5 and 6), the PacBio HiFi datasets were nevertheless processed with `--ont`, so that the nanopore-trained triplet-graph models were applied under exactly the settings in which they were fitted. ClairS-TO and DeepSomatic-TO were run with their PacBio HiFi models on the same data, so both the somatic recalibration and the tumor DNA fraction regression were evaluated off their training platform. This isolates the transfer of the trained models from any effect of the platform preset, and the resulting accuracy is therefore a lower bound on what `--pb` would provide.

**LongPhase v1.7.3** (https://github.com/twolinin/longphase)

```
longphase phase \
-b ${TUMOR_BAM} \
-r ${REF} \
-s ${VCF} \
-t ${THREADS} \
--ont \
-o ${OUTPUT}
```

**WhatsHap v2.1** (https://github.com/whatshap/whatshap)

```
whatshap phase \
--ignore-read-groups \
-o ${OUTPUT}.vcf \
-r ${REF} \
${VCF} \
${TUMOR_BAM}
```

**HapCUT2 v1.3.4** (https://github.com/vibansal/HapCUT2)

```
HapCUT2/build/extractHAIRS \
--ont 1 \
--bam ${TUMOR_BAM} \
--VCF ${VCF} \
--out ${OUTPUT}.fragment_file \
--ref ${REF} \
&& \
HapCUT2/build/HAPCUT2 \
--fragments ${OUTPUT}.fragment_file \
--VCF ${VCF} \
--output ${OUTPUT}
```

#### 5.4 Tumor DNA Fraction Estimation

**ASCAT v3.2.0** (https://github.com/VanLoo-lab/ascat)

```
R_SCRIPT=$(mktemp --suffix=.R)

cat <<EOF > $R_SCRIPT
library(ASCAT)

setwd("${OUTPUT_PREFIX}")

ascat.prepareHTS(
  tumorseqfile = "${TUMOR_BAM}",
  normalseqfile = "${NORMAL_BAM}",
  tumorname = "tumor",
  normalname = "normal",
  allelecounter_exe = "${ALLELE_COUNTER}",
  skip_allele_counting_normal = FALSE,
  skip_allele_counting_tumor = FALSE,
  alleles.prefix = "${ALLELES_PREFIX}",
  loci.prefix = "${LOCI_PREFIX}",
  gender = "${gender}",
  genomeVersion = "hg38",
  nthreads = 50,
  tumorLogR_file = "Tumor_LogR.txt",
  tumorBAF_file = "Tumor_BAF.txt",
  normalLogR_file = "Germline_LogR.txt",
  normalBAF_file = "Germline_BAF.txt",
  loci_binsize = 500,
  min_base_qual= 10,
  additional_allelecounter_flags="-f 0")

ascat.bc = ascat.loadData(
  Tumor_LogR_file = "Tumor_LogR.txt",
  Tumor_BAF_file = "Tumor_BAF.txt",
  Germline_LogR_file = "Germline_LogR.txt",
  Germline_BAF_file = "Germline_BAF.txt",
  gender = "${gender}",
  genomeVersion = "hg38"
)

ascat.plotRawData(ascat.bc, img.prefix = "Before_correction_")

ascat.bc = ascat.aspcf(ascat.bc)
ascat.plotSegmentedData(ascat.bc)
ascat.output = ascat.runAscat(ascat.bc, write_segments = TRUE)
EOF

# run R script
Rscript $R_SCRIPT
```

**PURPLE v4.0, with AMBER v4.0 and COBALT v1.16.0** (https://github.com/hartwigmedical/hmftools)

Purity and ploidy were estimated with PURPLE from AMBER B-allele frequencies and COBALT read-depth ratios. All three tools were run from the container distributed for the official PacBio HiFi-somatic-WDL pipeline, pinned by image digest so that the exact build can be recovered, and the Hartwig Medical Foundation reference bundle `hmf_pipeline_resources.38_v2.3.0` (https://storage.googleapis.com/hmf-public/HMFtools-Resources/pipeline/oncoanalyser/2.3/38/) supplied the AMBER germline site list, the COBALT GC profile and the Ensembl annotation directory. Both segmentation steps used a piecewise-constant-fit penalty of $\gamma = 1000$, the value adopted by that pipeline for long-read data: AMBER was run from a build with this value fixed (the jar named `amber.gamma1000.jar`) and COBALT through `-pcf_gamma 1000`. The purity search was bounded to the interval $[0.08, 0.99]$.

All three tools were run in tumor-only mode, which each of them selects by the absence of a reference sample, so that PURPLE is evaluated under the same tumor-only constraint as LongPhase-TO. In place of the read-depth baseline that a matched normal would otherwise provide, COBALT was given the consensus diploid-regions file `DiploidRegions.38.bed.gz` from the same reference bundle, through `-tumor_only_diploid_bed`. That file is a fixed cohort-level resource built by the tool's authors from a panel of reference samples, recording the genomic windows that typically remain diploid, and is thus the copy-number analogue of a panel of normals: it is identical for every sample, is independent of the tumor under analysis, and introduces no matched-normal information. PURPLE was additionally supplied with the ClairS-TO v0.3.0 somatic SNV calls used throughout this study, through `-somatic_vcf`, which lets the purity fit draw on somatic allele frequencies in addition to the AMBER and COBALT profiles.

```
# Container pinned by digest. The image reference is a single argument; the
# digest is wrapped here for display only.
docker pull quay.io/pacbio/purple@sha256:
    9074d56ad46f3d6804f1ae6e45a2ac8c300effc5efe120020d3dcd23edb3b063

# AMBER: B-allele frequencies at germline heterozygous sites (tumor-only)
java -jar amber.gamma1000.jar \
  -tumor ${TUMOR} -tumor_bam ${TUMOR_MIX_BAM} \
  -ref_genome ${REF} -ref_genome_version V38 \
  -loci AmberGermlineSites.38.tsv.gz \
  -output_dir ${AMBER_DIR}

# COBALT: read-depth ratios (tumor-only; the consensus diploid regions supply
# the depth baseline that a matched normal would otherwise provide)
java -jar cobalt.jar \
  -tumor ${TUMOR} -tumor_bam ${TUMOR_MIX_BAM} \
  -ref_genome ${REF} -gc_profile GC_profile.1000bp.38.cnp \
  -tumor_only_diploid_bed DiploidRegions.38.bed.gz \
  -pcf_gamma 1000 -validation_stringency SILENT \
  -output_dir ${COBALT_DIR}

# PURPLE: joint purity and ploidy fit (tumor-only)
java -jar purple.jar \
  -tumor ${TUMOR} \
  -amber ${AMBER_DIR} -cobalt ${COBALT_DIR} \
  -ref_genome ${REF} -ref_genome_version 38 \
  -gc_profile GC_profile.1000bp.38.cnp \
  -ensembl_data_dir ${ENSEMBL_DIR} \
  -min_purity 0.08 -max_purity 0.99 \
  -somatic_vcf ${TUMOR}.somatic.snv.vcf.gz \
  -output_dir ${PURPLE_DIR}
```

#### 5.5 Benchmarking

**Som.py** (https://github.com/Illumina/hap.py)

```
docker run -i --rm \
  -v ${INPUT_DIR}:${INPUT_DIR} \
  -v ${OUTPUT_DIR}:${OUTPUT_DIR} \
  -e HGREF=${REF} \
  pkrusche/hap.py:latest \
  /opt/hap.py/bin/som.py \
  -N ${ANSWER_VCF} \
  ${DIFF_VCF} \
  -r ${REF} \
  -o ${OUTPUT_PATH} \
  --feature-table generic \
  ${BED_OPTION} \
  ${PASS_ONLY_OPTION}
```
